# The adaptor protein DAB2 cell-autonomously links APOE with disease-associated microglia (DAM)/TREM2 state dysregulation associated with Alzheimer’s disease

**DOI:** 10.64898/2026.11.12.766141

**Authors:** Melissa Barber, William D Andrews, Aleksej Kubelka, Heleni Singer, Silvia La Penna, Daria Gavriouchkina, Modesta Blunskyte-Hendley, Alexander Becalick, Xinran Hao, Timothy JY Birkle, Rebecca Northeast, Umran Yaman, Naciye Magusali, Zaynah Shazad, Lila Chapman, Richard C Siow, Dag Aärsland, Jennifer M Pocock, Brian W Howell, John G Parnavelas, John Hardy, Dervis A Salih

## Abstract

The neuron-secreted Reelin (RELN) protein and its adaptor DAB1 modify Alzheimer’s disease (AD) risk through genetic interaction with APOE4. While RELN and APOE4 exert antagonistic effects on amyloid and Tau pathology in mice, how risk is conferred in the human brain remains unclear. The related adaptor DAB2 endocytoses RELN/APOE receptors and, via LRP8, modulates non-canonical pro-inflammatory signalling in non-neuronal cells. We hypothesised that RELN and APOE signalling directly influence microglial function to modify AD risk. Integrating human brain transcriptomics, a *Reln*-deficient mouse, and *DAB1/DAB2* knock-down in *APOE* isogenic iPSC-microglia, we found that microglial *DAB1/DAB2* pathways were dynamically regulated across the lifespan, with *DAB2* dysregulated with increasing amyloid pathology. DAB2 transcriptional networks correlated with disease– and lipid-associated microglial states (DAM/LDAM) linked to networks expressing dementia-risk loci (TREM2, GRN, ACE, SORL1) and associated with endolysosomal and lipid-handling pathways. DAB2-knockdown cell-autonomously shifted microglia away from gene networks enriched in DAM/TREM2 states and towards pro-inflammatory cytokine-responsive states and TNFα-NFκB pathways. APOE4/E4 microglia showed elevated APOE and reduced DAB1pY232 under inflammatory stress, and blunted DAB2-knockdown transcriptomic responses, consistent with pathway-level genetic interaction. In vivo, RELN-deficiency reduced microglial ramification and increased CD68. These findings identify DAB2 as a regulator of microglial state transitions and a modifier of dementia-risk networks linking APOE genotype to inflammatory, phagocytic and amyloid-beta clearance pathways relevant to AD.

- The microglial endocytic DAB2 adaptor links APOE and RELN genetic risk pathways in the human brain and is dysregulated with increasing amyloid pathology in Alzheimer’s Disease
- DAB2 cell-autonomously sustains iPSC-microglial disease-associated DAM/TREM2 activated states which are associated with dementia-risk loci
- Loss of DAB2 shifts APOE iPSC-microglia away from protective DAM/TREM2 states and towards pro-inflammatory cytokine-responsive states
- DAB2 knockdown inflammatory/senescent responses are attenuated in an APOE4/E4 background suggesting genetic interaction at the pathway level

## Introduction

Reelin (RELN) is a neuron-secreted extracellular-matrix glycoprotein best known for orchestrating neuronal migration and cortical lamination during development^1,2^. RELN binds the LDL-receptor-family proteins LRP8 (ApoER2) and VLDLR to activate the intracellular adaptor protein DAB1, which regulates cytoskeletal organisation, neuronal positioning and synaptic plasticity^3–6^. Beyond development, genetic studies implicate RELN and DAB1 as modifiers of familial^7^ and sporadic Alzheimer’s disease (AD), through genetic interaction with the APOE locus^8^. Although mouse models show APOE4 and RELN exert opposing effects on amyloid and tau pathology^9–12^, the mechanisms by which these pathways modify AD-risk in the human brain remain poorly defined. Notably, the paralogous clathrin dependent adaptor DAB2 endocytoses the RELN and APOE receptor LRP8^13,14^, linking both AD-risk pathways; with RELN-LRP8 promoting non-canonical NFκβ and eNOS signalling in human endothelial cells via DAB2^15,16^. Whether DAB2 transduces RELN signalling is unclear^17^, however its overexpression in amyloid mouse models has been shown to activate anti-inflammatory TGFβ/SMAD signalling that attenuates microglial activation and amyloid pathology^18^ offering a non-canonical DAB2 route through which RELN and APOE receptor availability and inflammatory signalling could be modulated in human microglia.

Microglia are central to both brain development and neurodegeneration, contributing to synaptic pruning, immune surveillance and the response to amyloid^19–21^. In early AD they mount a protective response, limiting amyloid-related damage through phagocytosis, the removal of debris and amyloid-damaged spines, and by providing neurotrophic support and resolving inflammation via lipid signalling^22,23^. However, as disease advances, dysregulated transitions between microglial states drive chronic neuroinflammation and senescent states that exacerbate synaptic loss, tau phosphorylation and neurodegeneration^20,24–26^. Whether RELN signalling directly shapes these microglial state transitions, and how this intersects with APOE genotype, has not been examined before.

Here, we integrate transcriptomic analyses of human brain datasets with *DAB1/DAB2* genetic perturbation studies in APOE-isogenic iPSC-derived microglia and an in vivo Reln-deficient mouse model to define how RELN signalling, through its convergence with APOE, shapes microglial biology. We identify the endocytic adaptor DAB2 as a regulator of gene pathways underlying microglial DAM/TREM2 state transitions and show that its disruption shifts microglia toward pro-inflammatory, cytokine-responsive (CRM) states, corresponding to gene modules associated with dementia risk-loci and linking APOE genotype with dysregulated transcriptional profiles associated with phagocytosis, lipid handling and amyloid-processing pathways, relevant to AD progression.

## Results

### DAB2 signalling is dynamically regulated in microglia across the human lifespan and becomes dysregulated in AD

To determine whether RELN-DAB1 or RELN-DAB2 signalling is active in microglia within the human brain, we analysed publicly available microglial transcriptomic datasets spanning brain development^27^, ageing^28^ and AD^29^. These were statistically scored against curated RELN signalling gene sets derived from protein-protein interaction databases (Reactome, STRING, MSigDB), with convergent gene sets used as RELN-axis signalling readouts (Supplementary Table 1). Because LRP8 and VLDLR lipoprotein receptors bind both APOE and RELN, and DAB2 has also been shown to interact with shared receptor LRP8^13^ these pathways reflect convergent RELN/APOE signalling pathways on-DAB1 or non-canonical-DAB2 adaptor proteins (Fig. 1a). While RELN signalling has mostly been studied in a neuronal context, previous studies have suggested that microglia may express RELN receptors VLDLR^30^ and LRP8^31,32^ and adaptor DAB2^18,33^; with systemic RELN recently shown to interact with LRP8-DAB2 and promote pro-inflammatory NFκβ signalling in human endothelial cells^15^ (Fig. 1a). We hypothesised that DAB2 may modulate RELN– and APOE-inflammatory signalling in human microglia. To assess the RELN-axis in microglia in the human brain, we first looked at expression levels of canonical RELN receptors and the DAB1 and DAB2 adaptors within the healthy adult brain across the adult lifespan (young 34-57years; n=26 donors vs aged 62-102years; n=33 donors; Galatro 2017; GSE99074). This showed LRP8, VLDLR, and DAB1 were stably expressed at modest levels, whereas the APOE receptor LRP1 and the non-canonical adaptor DAB2 were expressed ∼13-fold and 9-fold higher than LRP8 and DAB1 respectively, with DAB2 upregulated with ageing (+1.078, *padj=0.0291) (Fig. 1b). RELN receptors were expressed at comparable levels to integrin receptor subunits (ITGB1, ITGB8), with ITGB1 corresponding to a known microglial receptor that endocytoses fibrillar amyloid-beta and promotes microglial activation^34,35^.

**Figure 1.**
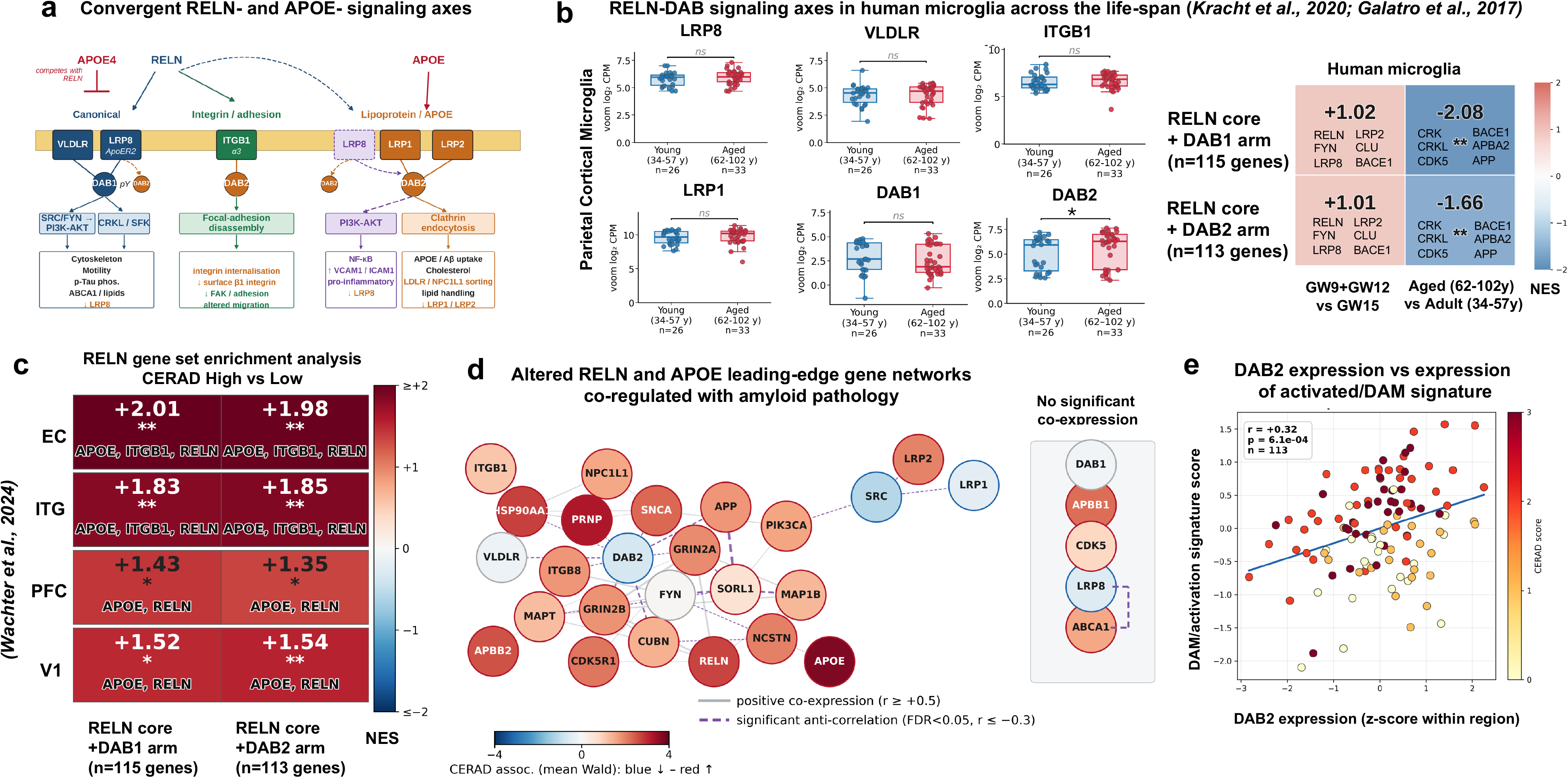
RELN-DAB and APOE-signalling components are expressed in human microglia across a lifetime and are coordinately dysregulated in Alzheimer’s Disease. **a) Convergence of canonical RELN-DAB1 and non-canonical RELN-DAB2 signalling with APOE.** Schematic of RELN signalling and its convergence with APOE signalling, derived from neuronal, brain endothelial, cardiomyocyte, retinal ganglion neurons and lymphatic endothelial cell studies. RELN engages i) the canonical lipoprotein-receptor-family members LRP8/VLDLR (blue arrows) to activate DAB1; with SRC/FYN-mediated tyrosine phosphorylation (pY) driving PI3K-AKT underlying classical effects of RELN on motility, cytoskeletal remodeling and inhibition of Tau-phosphorylation^1,3,50,127^; and activates FYN-mediated DAB1 phosphorylation with adaptors CRK/CRKL transducing downstream pathways that engage ii) integrin ITGB1 (***α***3***β***1) (green arrows) to alter DAB1 availability^36–38^, or matrix-adhesion also through N-cadherins^128^. APOE (red arrows) competitively bids with RELN at VLDLR/LRP8 and separately engages LRP1/LRP2 with LRP2/LRP8 predicted to be interact with DAB2^14,80,83,84,86^, and triggering clathrin-mediated endocytosis of APOE bound cargo and A***β***. The dashed orange arrow denotes RELN-LRP8-DAB2 signalling reported in brain endothelial cells showing convergence at the DAB2/endocytic adaptor level which activates non-canonical pro-inflammatory PI3K/AKT/NF-κβ signalling and eNOS signalling in human endothelial cells^15,16^. Microglial cell line studies highlight RELN-VLDLR-LRP8-DAB1pY also activates the cholesterol and lipid efflux transporter ABCA1 providing cross-talk between APOE cholesterol/lipid handling and RELN-dependent signalling^32^. **b) RELN and APOE signalling-axis genes are expressed in human microglia, with DAB2 upregulated with ageing.** Box plots show voom normalized log2 counts per million (CPM) of RELN receptors LRP8, VLDLR, ITGB1, the LRP1 APOE receptor and the adaptor proteins DAB1 and DAB2 in young adult donors (34-57years, n=26) versus aged adult donors (62-102years, n=33) in bulk microglia from the parietal cortex (Galatro and others 2017; GSE99074): individual transcripts were unchanged (ns) except for DAB2 which was upregulated with ageing. (LRP8: median young 5.96, median aged 5.98, Mann-Whitney q0.41; Aged vs Young log2FC-0.14, padj 0.60; VLDLR: median young 4.52, median aged 4.69, Mann-Whitney q0.39; log2FC –0.22, padj 0.55; ITGB1: median young 6.31, median aged 6.84, Mann-Whitney q0.41; –0.25, padj 0.32; LRP1: median young 9.66, median aged 10.18, Mann-Whitney q0.40; log2FC +0.49, padj 0.11; DAB1: median young 2.67, median aged 1.87, Mann-Whitney p0.41; log2FC –0.47, padj 0.44) DAB2: median young 5.91, median aged 6.29, Mann-Whitney p0.15; log2FC +1.08, *padj 0.029) **Heatmap** shows multilevel fgsea (infinite permutations; Korotkevich et al., 2021^123^) normalised enrichment scores (NES) for the curated RELN-core +DAB1 (n=115 genes) and +DAB2-(n=113 genes) protein-protein interaction datasets (STRINGv12;Reactome;MSigDB) tested in fetal human cortical microglia (sum aggregated expression levels across cells/sample) (GW9+GW12 n= 7 vs GW15 n= 3 donors; (Kracht et al., 2020^27^); exploratory given low n); and in aged (62-102y, n=33) compared with young (34-57years, n=26) parietal microglia (genes ranked by DESeq2 Wald-z). Leading edge genes of RELN-axis (left column) or APOE-axis (right column) contributing to NES are shown per cell. BH *padj<0.05, **<0.01, ***<0.001. Color-bar: NES range ± 3.0. **c) RELN and APOE signalling is dysregulated in microglia in the AD brain. Left heatmap** shows gene-set enrichment (fgsea multilevel) of the RELN-core signature resolved into its DAB1 and DAB2 arms, contrasted with genes ranked by Wald statistics between high (moderate/frequent) compared with low (none/sparse) CERAD neuritic amyloid-plaque scores in microglia from four regions of the AD progression atlas: entorhinal cortex (EC), inferior temporal gyrus (ITG), prefrontal cortex (PFC) and primary visual cortex (V1) (rows). Values show normalised enrichment score (NES); positive NES (red) indicates greater enrichment at higher CERAD stage. Asterisk denote Benjamini-Hochberg adjusted significance (*padj<0.05, **padj<0.01, ***padj<0.001). Italic labels list RELN/APOE axis leading edge genes showing APOE, RELN, ITGB1 driving the enrichment). **d) Co-expression network of leading-edge genes from the RELN-core (DAB1– and DAB2-arm) enrichment analyses.** Each node is a leading-edge gene, coloured by the direction and strength of its association with CERAD scores (mean DESeq2 Wald statistic across entorhinal cortex, inferior temporal gyrus, prefrontal cortex and visual cortex). Edges denote donor-level co-expression across microglial transcriptomes (n=113 donor x region samples): grey solid edges show positive co-expression (Pearson r≥+0.5); purple dashed edges, significant anti-correlation (Benjamini-Hochberg FDR <0.05, r≤-0.3). DAB2 shows modest decline with increasing CERAD scores at the single-gene level, but at the network level was part of a significant inverse co-expression hub, anti-correlating with the APP module and with its own RELN receptor VLDLR. Genes not connected with the main amyloid-associated module at these thresholds are shown in the grey box on the right including the canonical RELN transduction components DAB1 and LRP8, with LRP8 and ABCA1 however significantly anti-correlated with each other. **e) DAB2 transcript expression correlates with a microglial DAM/activation signature in the human brain (AD Progression Atlas).** Scatter plots show z-scored expression of DAB2 plotted against a microglial DAM/activation signature, defined as the mean within-region z-scored expression of 9 canonical disease-associated (DAM)/activated microglial markers (TREM2, TYROBP, GPNMB, AXL, LPL, CD9, CD68, CTSD, CAPG; core DAM genes from Keren-Shaul et al., (2017)^46^ and Mancuso et al., (2024)^26^ with CAPG from Monasor et al., (2020)^49^ & Pesämaa et al., (2023)^51^. Each dot is from a single donor and brain region (n=113 region-donor pseudobulk samples from 32 donors across entorhinal cortex (EC), inferior temporal gyrus (ITG), prefrontal cortex (PFC) and visual area 1 (V1), colour-coded by CERAD score (yellow to dark red = low to high scores). DAB2 positively correlates with DAM/activation score (Pearson r = +0.32, p=6.1×10^−4^). Blue line shows regression fit.

To more comprehensively assess RELN-signalling at the pathway level, we carried out a series of gene statistical enrichment analyses to determine whether genes in the RELN-DAB1 and RELN-DAB2 pathways are over-represented in pathways relevant to microglia and AD pathology compared with a random assortment of genes. This showed that the RELN-signalling axis was unchanged in microglia between early and mid-neurogenesis but was depleted in microglia with ageing (RELNcore-DAB1 NES –2.08, RELNcore-DAB2 NES –1.66, both p<0.01^27,28^) with leading-edge genes corresponding to CRK/CRKL adaptor proteins which are thought to relay pDAB1-signalling to downstream RAP1 that modulates integrin and N-cadherin dependent adhesion underlying cell motility^36–38^, or the CDK5 kinase which synergises RELN receptor-DAB1 signalling through positive phosphorylation of the DAB1p45 isoform^39–41^. Analysis of microglia in AD, showed an enrichment in RELN-pathways associated with increasing CERAD/neuritic amyloid-plaque burden (high: moderate/frequent vs low: none/sparse) and reflecting advancing AD pathology across brain regions, but most strongly enriched in the early-vulnerable entorhinal cortex followed by the inferior temporal gyrus and later-affected visual areas and prefrontal cortex (Fig. 1c). Contrasts with Braak staging (Tau pathology) showed enrichment in RELN-DAB1 and RELN-DAB2 pathways in microglia in early vulnerable Tau-affected entorhinal cortex (EC, NES +1.65) and ITG (NES +1.49; both p<0.01), with composite pathology scores (Braak+ CERAD + diagnosis) only significant in the ITG (Supplementary Fig. 1d). Leading-edge gene analysis showed enrichment with increasing plaque pathology comprised the APOE gene, with RELN and ITGB1 contributing across brain regions. To understand if this reflects a bona-fide enrichment in microglial RELN signalling, we carried out a leading-edge gene co-expression network from microglial transcriptomes, computing gene-gene correlations within each brain region across donors and correlated this with CERAD scores/neuritic-plaque burden (Fig. 1d. This showed that APOE, APP and other amyloid-associated genes were positively correlated with CERAD scores/neuritic plaque burden, whereas the canonical RELN-transduction LRP8 receptor and DAB1 adaptor showed no overall association with amyloid and lay outside the amyloid co-expression module (Fig. 1d). Although DAB2 also showed only a modest decline with CERAD scores at the single gene level, at the network level it was a significant inverse co-expression hub, anti-correlating with APP and associated amyloid-genes (CUBN, SNCA and PRNP) and, notably, with the RELN receptor VLDLR. Among APOE signalling components, APOE and its receptor LRP2 were upregulated with amyloid whereas LRP1 was reduced. To better resolve potential brain region-specific effects in RELN and APOE signalling, we looked at single-gene associations in an exploratory analysis by brain-region and APOE genotype. While this needs to be interpreted cautiously given there were few APOE4 carriers (APOE4/E4 n=1, APOE3/E4 n=6; APOE3/E3 n= 22 individuals), this showed microglial DAB1 declined with increasing amyloid, Tau/Braak and composite-stage pathology in a region-specific manner, strongest in the early-affected entorhinal cortex and later affected prefrontal cortex and predominantly in APOE3/E3 carriers; whereas DAB2 showed a region-specific decline with amyloid pathology, that in an exploratory genotype-stratified analysis, was most prevalent in APOE4 carriers (n=6-7 carriers; Fig. 1e & Supplementary Fig. 1a). Together, these findings suggest that components of the RELN– and APOE-response pathways are enriched in human microglia and are dynamically regulated across the healthy lifespan, with dysregulation of the APOE-DAB2 axis emerging with increasing amyloid pathology, especially in the AD-afflicted brain.

### RELN-signalling is enriched in specific developmental and AD relevant microglial states

Microglia are highly plastic cells which adopt distinct functional states essential for their pleiotropic functions, including mounting protective responses at the early stages of AD. To determine whether RELN signalling is associated with specific microglial states, we performed gene enrichment testing of RELN-curated lists with pseudo-bulk microglial RNA sequencing datasets spanning embryonic^42^, postnatal^43^, aged and AD datasets^26,44–47^. This showed that a few RELN/APOE pathway genes significantly overlapped with early developmental Arg1+, metabolic, Spp1+/ATM axon-tract or Hmox1+/stress-responsive states (<=3 genes each, padj<0.01; Supplementary Fig. 2a). To assess relevance to AD, we compared RELN-and APOE-associated pathways with amyloid-responsive transcriptional signatures of APOE iPSC-microglia xenotransplanted into amyloid mouse models from Mancuso et al., (2024)^26^, and with curated AD-relevant microglial signatures from published data in the human brain^24,46,48^. This showed that the RELNcore-DAB1 (NES +1.38; p<0.05) and RELNcore-DAB2 (NES +1.45; p<0.05) arms were both enriched in disease-associated (DAM) microglia (Supplementary Fig. 2b). Gene overlap studies of lipid-droplet accumulating microglial states, recently reported to be enriched in APOE4/E4 patients with AD and shown to be important in promoting the transition of amyloid to Tau pathology^24,47^, showed significant enrichment in the RELNcore+DAB1 and RELNcore+DAB2 arms (LDAM, TREM2-lipid and LXR/cholesterol states; p<0.05 to <0.001) (Supplementary Fig. 2c).

In the human brain, advancing AD pathology (AD Progression Atlas^29^), was associated with a broad transcriptional enrichment of activated microglial states including DAM, HLA and dystrophic microglia across all brain regions, with ribosomal microglia enriched across all brain regions except for within the entorhinal cortex, and LDAM specifically enriched in the ITG and V1 cortex. This contrasted with a global depletion of cytokine-responsive states across all brain regions, with ribosomal and proliferating microglia specifically depleted within the early vulnerable entorhinal cortex (Supplementary Fig. 1b). We next correlated DAB1 and DAB2 expression per donor across brain regions with increasing CERAD scores/neuritic-plaque pathology with an average z-score expression of microglial activation signatures (TREM2, TYROBP, GPNMB, AXL, LPL, CD9, CD68, CTSD, CAPG)^26,46,49^. This showed that DAB2, but not DAB1, expression correlated with a microglial DAM/activation signature in the human brain with advancing neuritic plaque-burden/CERAD scores (r= +0.32, p=6.1×10^−4^, 113 total regions from n=32 donors) (Fig. 1e & Supplementary Fig. 1c). To further understand the relevance of RELN-response pathways enriched in DAM and LDAM/LAM microglia to AD-microglial states across the lifespan, we carried out a gene overlap analysis between embryonic and AD-relevant microglial transcriptomic states. Early fetal homeostatic, phagocytic, amyloid– and lipid-metabolism programmes (Hm C4/ATM, Hm C7b, Hm C1/Arg, Li C1/PAM; p<0.001) transcriptionally overlapped with activated disease– and lipid-associated states (DAM/LDAM), and amyloid responsive ARM states although this comprised few genes; while stress-responsive Li C6 (IEG), axon-tract HmC4 (ATM) and homeostatic (Hm C7b, Li C0) microglia showed significant gene overlap (p<0.05-000.1) with cytokine-responsive CRM-1 states (Supplementary Fig. 2d). Together, these findings suggest that RELN and APOE signalling is associated with human microglial states across the lifespan and may influence microglial disease-activated/DAM states in Alzheimer’s disease.

### RELN signalling components are expressed in human APOE iPSC-derived microglia

To directly test whether microglia express pathways able to respond to RELN-DAB1/DAB2 signalling and assess its interaction with APOE, we employed human APOE iPSC-microglial models. We first characterised the expression of RELN signalling pathway components in human iPSC-derived microglia across APOE genotypes using two independent cell lines BIONIO37-A/4 (EBiSC) and ioMG (Bit.bio). Both canonical receptors (LRP8, VLDLR) and DAB1/DAB2 adaptors were expressed at the RNA level at equivalent levels across genotypes, except for APOE, which was significantly ∼6-7 fold higher in APOE4/E4 BIONIO37A/A-4 compared with APOE3/E3 iPSC-microglia as assessed by RT-qPCR (Fig. 2a, Supplementary Fig. 3a, Supplementary Fig. 4a-b). Staining with anti-DAB1 antibody, previously validated in DAB1 knockout tissue^3,50^, or against its tyrosine-phosphorylated forms (Y232, Y198) which are indicative of functional RELN receptor-DAB1 engagement, showed both were detected at the protein level (Fig 2b, Supplementary Fig. 3b), despite its low transcript expression.

**Figure 2.**
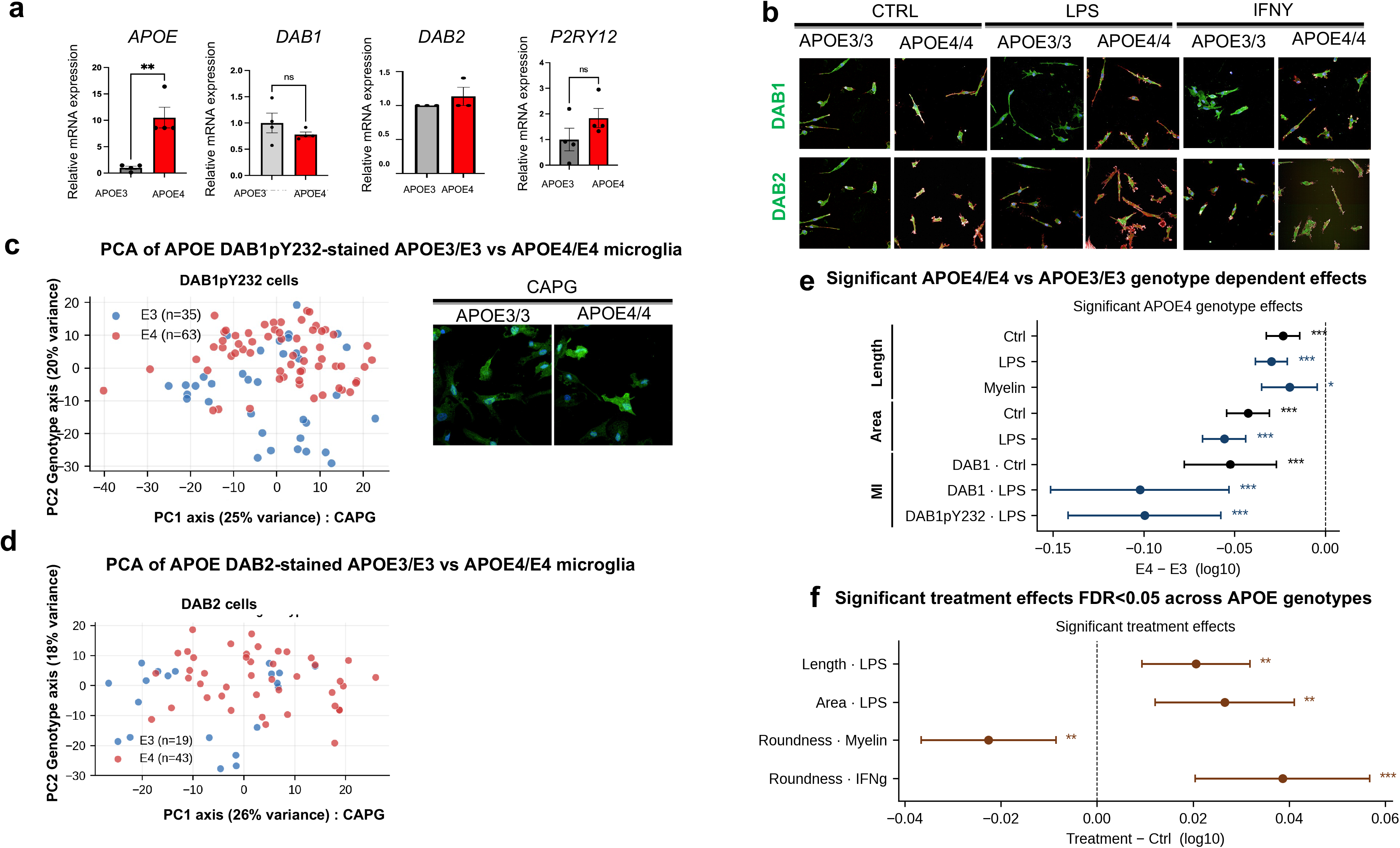
Human APOE iPSC-derived microglia express RELN-signalling components. **a)** Relative gene expression levels obtained through qRT-PCR of *APOE, DAB1, DAB2* Reelin signalling components and the microglial marker P2RY12. **b) DAB1 and DAB2 proteins are expressed in APOE iPSC-microglia.** Day 14 APOE3/3 (BIONIO37A) and APOE4/4 (BIONIO37A-4) differentiated isogenic iPSC-microglia immuno-stained for DAB1 and DAB2 and counter-stained with actin-labelling phalloidin under vehicle control (CTRL), or pro-inflammatory LPS and IFNγ conditions. **c) APOE genotype dependent effects on DAB1pY232 iPSC-microglial features co-stained with CAPG.** Principal component analysis (PCA; PC1 vs PC2) of automated high-content imaging features – including DAB1pY232 protein intensity, texture and cell-morphology parameters (cross-experiment, z-scored within plate) extracted by Cell Painting segmentation and quantified in Harmony software (Opera Phenix High-Content Imaging System). Genotypes separate along PC2 (20% variance, linear mixed model with experiment as random effect, p=1.4×10^−4^); APOE3/E3 n=3 experiments/35 wells/17,689 cells); APOE4/E4 n=4 experiments/63 wells/ 46,775cells), whereas PC1 (25% of variance) is driven by CAPG texture features. Each dot is a single well from one experiment. The bar plot shows PC2 loadings (the genotype axis), which is predominantly driven by DAB1pY232 texture features (purple). Inserts show immuno-staining for CAPG DAM marker in APOE iPSC-microglia. **d) APOE genotype dependent effects on DAB2 iPSC-microglial features co-stained with CAPG.** Principal component analysis (PCA; PC1 vs PC2) of automated high-content imaging features – including DAB2 protein intensity, texture and cell-morphology parameters (cross-experiment, z-scored within plate) extracted by Cell Painting segmentation and quantified in Harmony software (Opera Phenix High-Content Imaging System; “FinalPlate” pipeline, Methods) (APOE3/E3 n=3 independent experiments (19wells/9,804cells); APOE4/E4 n=5 independent experiments, 43wells/29,926cells). Each dot shows a single plotted well on PC1 versus the genotype-associated PC2 axes, coloured by APOE3/E3 (blue) or APOE4/E4 (red) genotypes. PC1 accounts for 26% of variance and PC2, the genotype axis, for 18%; APO3/E3 and APOE4/E4 iPSC-microglia separate along PC2 (linear mixed model with experiment as random effect, p=3.8×10^−5^). **e) APOE4/E4 microglia show reduced DAB1 signalling and smaller, shorter morphologies.** Forest plot of the log10 mean-fold difference (E4−E3) in single-cell DAB1 and phospho-DAB1 (DAB1-pY232) intensity and in whole-cell morphology (cell area and length) across treatments (Ctrl, IFNγ, LPS, Myelin 3 mg/mL); negative values indicate lower levels in APOE4/E4 than APOE3/E3. APOE4/E4 microglia had significantly lower DAB1 (Ctrl, LPS) and DAB1-pY232 (LPS) and were significantly smaller with shorter processes (reduced cell area at Ctrl and LPS; reduced cell length at Ctrl, LPS and Myelin); DAB2 did not differ significantly between genotypes. Points coloured by significance (dark blue FDR<0.01; light blue FDR<0.05; grey FDR≥0.05); error bars = 95% CI; Estimates come from a cross-plate linear mixed-effects model (experiment as random effect) with Benjamini–Hochberg FDR; using vehicle-treated wells as internal controls. n per condition (independent experiments, wells/cells; APOE3/E3 vs APOE4/E4): **DAB1** — Ctrl n=5 experiments, 11wells/3,723cells vs n=5 experiments, 15wells/8,773cells; LPS n=4 experiments, 10wells/3,179cells vs n=5 experiments, 17wells/11,185cells. **DAB1-pY232** — Ctrl n=4 experiments, 10wells/3,050cells vs n=4 experiments, 14wells/9,342cells; LPS n=3 experiments, 8wells/2,726cells vs n=4 experiments, 15wells/11,447cells. **Whole-cell morphology** (area, length, roundness; Ctrl n=10experiments, 78wells/30,027cells vs n=11 experiments, 73wells/31,493cells; IFNγ n=6 experiments, 24wells/9,225cells vs n=8 experiments, 51wells/23,834cells; LPS n=6 experiments, 28wells/8,990cells vs n=7 experiments, 51wells/31,796cells; Myelin n=8 experiments, 52wells/25,706cells vs n=9 experiments, 84wells/39,410cells). **f) Inflammatory and myelin challenges significantly alter APOE iPSC-microglial morphology.** Forest plot of significant treatment effects (FDR<0.05) from linear mixed models (treatment vs control, genotype-adjusted, experiment as random effect) across all imaging plates, APOE3/E3 and APOE4/E4 pooled samples sizes (n= independent experiments, wells/cells) (Ctrl: n=12 experiments, 151wells/61,250cells; LPS: n=8 experiments, 79 wells/40,786 cells; IFN*γ*: n=9 experiments, 75wells/33,059cells; Myelin: n=9 experiments, 136wells/65,116cells). Points are the estimated treatment-control log10 mean-fold difference (log_10_ scale for area and length; raw scale for roundness) with 95% confidence intervals; colour shows false-discovery rate FDR (dark p<0.01; light p<0.05). IFN*γ* increases roundness, LPS increases cell area and length, and myelin decreases roundness. Plots show significant findings only.

Previous GWAS studies have shown that APOE4 genetically interacts with RELN-DAB1^8^. In addition, our previous exploratory analyses of microglia in the human AD-afflicted brain suggest that DAB2 is downregulated in APOE4/E4 carriers and in response to increasing amyloid pathology and is positively correlated with microglial activation markers (Supplementary Fig. 1a, 1c). We therefore next asked if DAB1/DAB2 protein expression is altered in microglia in different APOE genetic backgrounds, using an automated single-cell segmentation algorithm of APOE iPSC-microglia co-labelled with immuno-localised DAB1 or DAB2 proteins and the activated microglial marker CAPG^49,51^ (Fig. 2c-d). This protein-level analysis captured 584 different intensity, texture and morphological features based on DAB1/DAB2 and CAPG staining. Principal component analysis (PCA) of the extracted features showed that the DAB1pY232 texture features distinguished APOE3/3 compared with APOE4/E4 microglia (PC2 20% variance) (Fig. 2c, Supplementary Fig. 3c), while differential DAB2 feature expressions based on intensity of immunostaining did not diverge across APOE genotypes (Fig. 2d). APOE4/E4 microglia showed reduced DAB1 intensity staining, and smaller area and shorter-process morphologies (Fig. 2e) under vehicle-control conditions (mean log10 fold-difference: E4-E3=-0.05 ie ∼11% lower) relative to APOE3/E3 iPSC-microglia, with the effect roughly doubling in response to LPS (DAB1:mean log10 fold-difference: LPS=-0.10 ie ∼21% lower; all p<0.001), and also resulting in decreased phosphorylated DAB1 (pY232-DAB1: mean log10 fold-difference: LPS=-0.10 staining) indicative of decreased functional RELN-DAB1 signalling under these conditions (Fig. 2e). In addition, pro-inflammatory mediators significantly altered microglial morphologies across APOE genotypes, with IFN*γ* promoting rounder/more amoeboid shapes whereas exposure to myelin, a CNS demyelinating damage-associated substrate that enhances disease-associated microglia^52^ and, within the aged brain also senescent microglial states^53^, had the opposing effect and promoted more elongated morphologies (Fig. 2f). Thus, APOE4 lowers DAB1 and its pY232 phosphorylated-form and drives an activated morphology, whereas DAB2 abundance is genotype-invariant at the protein level in iPSC-microglia.

### APOE4 microglia exhibit disrupted lipid-handling, extracellular matrix and inflammatory pathways linked to RELN and APOE signalling

APOE4 has been shown to directly influence the microglial response to AD pathology through a cell-autonomous impairment of their morphologies and phagocytosis of amyloid-beta that mediates early protection against AD pathology, and by impairing lipid handling required for their crosstalk with neurons^48,54^. To better understand how RELN and APOE pathways alter microglial function, we carried out bulk RNA-sequencing of isogenic APOE iPSC-microglia. This identified 169 upregulated and 69 downregulated genes in APOE4/E4 versus APOE3/E3 under basal conditions (vehicle-treated cells, n=4/genotype; Fig. 3a). Our analysis showed that APOE4/E4 microglia were enriched for genes expressed by disease-associated (NES +1.75) and lipid-droplet-accumulating (NES +1.58; both p<0.05) microglia (Fig. 3b). Cross-comparison with RELN-core signalling pathways showed that the canonical RELN-DAB1 arm was modestly but significantly enriched in APOE4 microglia (NES +1.44, p<0.05; 34/115 genes), which was partly due to the strong upregulation of APOE itself (+11.9 fold), but also by the increased expression of RELN receptor VLDLR (Fig. 3b) and effectors AKT3, FYN, SPON1 (Supplementary Table 2), consistent with an enrichment in both RELN and APOE signalling pathways. Differential expression and targeted pathway analysis revealed that APOE4/E4 microglia showed a transcriptional enrichment in gene networks associated with the TNFα-NFκβ pro-inflammatory states (NES +1.96, p<0.001), coupled to extracellular-matrix remodelling and integrin-adhesion programmes important for regulating microglial motility (Extracellular matrix organisation NES +2.28; ECM Receptor interaction NES +2.36), and with pathways important for microglial interaction with neurons and synaptic pruning (Neuron projection guidance NES +2.37; Synapse Organisation NES= +1.87; both p<0.001)(Fig. 3c). This was associated with the concurrent suppression of cholesterol biosynthesis (Reactome NES –2.25, p<0.0001), antigen-receptor mediated signalling (NES= –1.46, p<0.05), immune response pathways (NES –1.51, p<0.001) as well as blunted Toll-like receptor signalling (NES –1.69 p<0.05))(Fig. 3c; Supplementary Table 2). The integrin-extracellular matrix pathways are important in mediating cell-adhesive interactions that underlie morphological changes, consistent with our observations of more rounded APOE4/E4 microglia under basal conditions and in response to specific inflammatory mediators (Fig. 2e-f). Together, this suggests that APOE4/E4 microglia show dysregulation in lipid handling, neuronal interactions, inflammatory and matrix-adhesion pathways consistent with a shift to DAM/LDAM microglial states associated with their adoption of altered morphologies under inflammatory conditions. Thus while APOE4/E4 microglia are enhanced in DAM– and LDAM-like transcriptional states, typically associated with endolysosomal/phagocytic function, these pathways were impaired suggested they may adopt maladaptive DAM and LDAM states.

**Figure 3.**
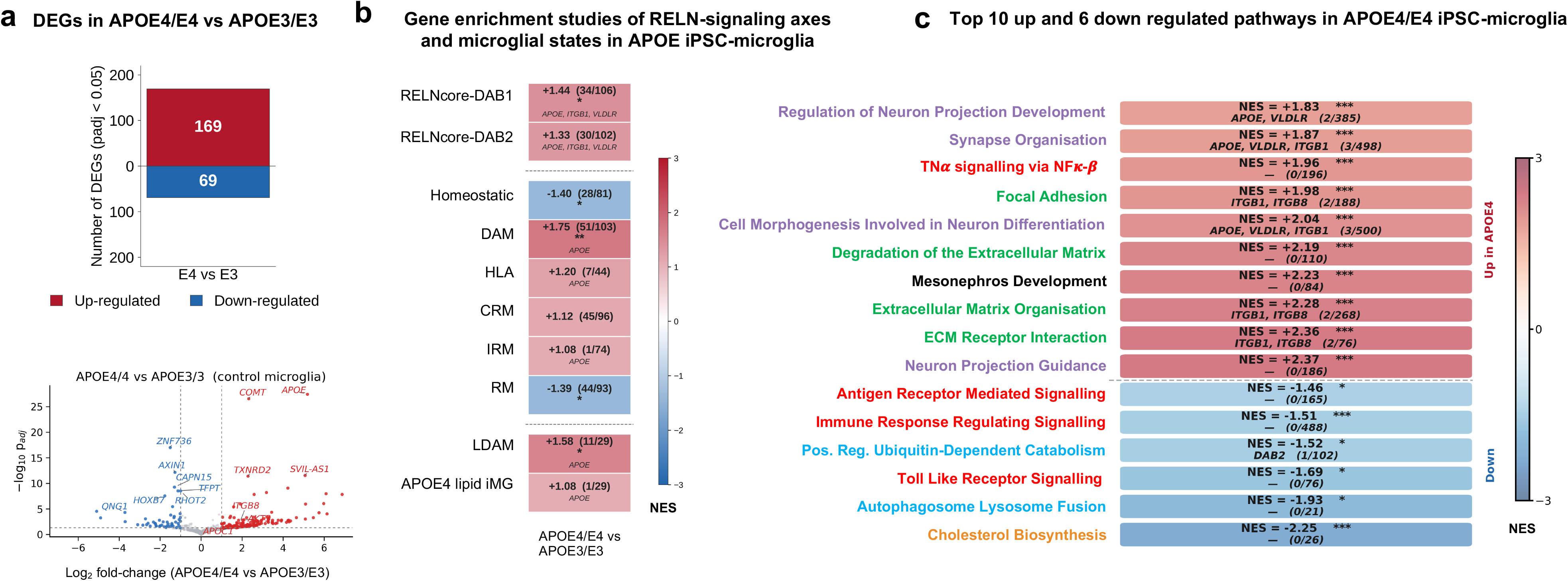
APOE4/E4 microglia are enriched in RELN signalling and activated microglial states but show altered lipid processing, extracellular matrix remodelling, phagocytic and inflammatory profiles. **a**) Up-(red) and down-(blue) regulated differentially expressed genes (DEGs)(padj<0.05) from DESeq2 (design ∼ Genotype) of Ctrl siRNA APOE4/E4 (n=4) vs APOE3/E3 (n=4) (BIONIO37A-4, BIONIO37A) microglia (169 up, 69 down). Right panel shows volcano plot of DEGs of APOE4/E4 vs APOE3/E3 Ctrl siRNA microglia with upregulated (Log2FC>1) genes shown in red and downregulated genes (Log2FC<-1) shown in blue. APOE is strongly upregulated in APOE4/E4 iPSC-microglia. b) Multilevel fgsea (10,000 permutations) of the RELN-signalling axes (RELNcore-DAB1 n=115 genes, RELNcore-DAB2 n=113 genes) and microglial AD state gene sets, enriched in APOE4/E4 (Ctrl siRNA, n=4) vs APOE3/E3 (Ctrl siRNA, n=4) microglia (BIONIO37-A4 vs BION37A, this study) and in APOE iPSC-microglia from Haney et al., 2024^24^. Fill = NES (±3). Brackets = leading-edge RELN genes/detected genes in the gene set (Wald-z-ranked, post-expression-filter). Bottom rows: fgsea NES of microglial AD-state gene sets (column 1, this study; column 2, Haney et al., 2024^24^). **c) Top 10 up– and 6 down-regulated microglial– and AD-focused pathways** (GO:BP, Reactome, KEGG or MSigDB Hallmark) from multi-level fgsea on the DESeq2 Wald-z-ranking of APOE4/E4 vs APOE3/E3 Ctrl siRNA microglia (n=4 per genotype). Fill=NES. Brackets = leading edge RELN/APOE genes/total pathway genes; pathway labels coloured by shared function (purple = microglial-neuronal interactions, red= inflammatory pathways, green = interaction with and remodelling of the extracellular matrix, blue = autophagosomal/endolysosomal function, and yellow = lipid-handling pathways).

### DAB2 siRNA knockdown in APOE iPSC-differentiated microglia shifts microglia from activated phagocytic states towards pro-inflammatory cytokine-responsive states

To directly test whether RELN/APOE signalling through DAB2 alters microglial states, DAB2 adaptor proteins were acutely silenced in siRNA knockdown experiments (Supplementary Figs. 5-6). Two major isoforms of DAB2, the prevalent DAB2-201 isoform (ENST00000320816.11), and the amyloid-responsive DAB2-202 isoform (ENST00000339788.11)^55^ were silenced through siRNA-96 that targets exon12 of DAB2-201 and exon11 of DAB2-202; whereas the DAB2-201 isoform was knocked-down with siRNA98 that targets its unique exon 10 (Supplementary Fig. 5a-c). This showed 659 upregulated and 330 downregulated genes in APOE3 iPSC-Microglia and 662 upregulated and 620 downregulated genes in APOE4 iPSC-microglia when the major DAB2-201 isoform was silenced (siRNA-98) (Supplementary Fig. 5c). To assess whether DAB2 knockdown response differed between APOE genotypes, the log2 foldchanges per gene were compared directly between APOE3/E3 and APOE4/E4 iPSC-microglia for each siRNA (Supplementary Fig. 5f). Among genes significantly regulated in both genotypes, foldchanges were concordant in direction and strongly correlated with magnitude (siRNA-98: Pearson r=0.86, Spearman ρ =0.90, n=324 genes; siRNA-96: Pearson r= 0.84, Spearman ρ= 0.67, n=23 genes), indicating that the DAB2-dependent transcriptional programme was qualitatively APOE-genotype-concordant, although the magnitude was different (Supplementary Fig. 5f). To understand whether DAB2 knockdown alters microglial states, a gene-enrichment analysis was carried out contrasting differentially expressed ranked genes with AD microglial states. This showed DAB2 depleted homeostatic (NES –2.38), DAM (NES –1.51), antigen-presenting HLA (NES – 2.22) and the lipid-droplet LDAM programme (NES –1.89), while enriching cytokine-responsive CRM (NES +2.15) and interferon-responsive IRM (NES +1.97) states in APOE3/E3 iPSC-microglia (all padj<0.01; Fig. 4a). APOE4/E4 microglia showed equivalent effects (homeostatic –1.93, DAM –2.02, HLA –1.78, LDAM –1.79; CRM +1.46, IRM +1.98). Knockdown of both isoforms of DAB2 (siRNA-96) downregulated gene networks associated with lysosomal and phagocytic pathways and amyloid-clearance pathways, and upregulated TNFα-NFκβ pro-inflammatory and senescence-associated signalling pathways (Fig. 4b), with concordant knockdown using both siRNAs also depleting the complement cascade (Supplementary Table 3). Pathway enrichment analysis between the two siRNA knockdown experiments showed that targeting both DAB2-201 and the amyloid-inducible DAB2-202 isoforms (siRNA-96) but not DAB2-201 alone (siRNA-98) showed a greater enrichment in extracellular matrix, amyloid-beta clearance, inflammatory response, lipid-handling pathways and in microglial state transitions towards antigen-presenting, interferon-responsive and disease associated states (Supplementary Fig. 5g; Supplementary Table 3). Targeting only DAB2-201 showed this enhanced the depletion of transcriptional pathways linked to DNA damage-induced senescence (Supplementary Fig. 5g). While this points to knockdown of different DAB2 isoforms affecting different microglial pathways, the observed effects may also be due to differences in absolute levels of DAB2. Differences in pathway-level NES between APOE3/E3 and APOE4/E4 iPSC-microglia were largely concordant although the magnitude was different with our interaction model showing DAB2 knockdown effects were blunted in an APOE4/E4 genetic background (Figure 4b, Supplementary Fig. 5f). To more closely assess for a potential APOE genotype interaction, we plotted the log2 foldchanges in leading-edge genes between APOE4/E4 and APOE3/E3 iPSC-microglia for the senescent/pro-inflammatory pathways predicted to be induced to a lower magnitude in APOE4/E4 iPSC-microglia. This showed that while APOE4/E4 iPSC-microglia engaged these pathways in the same direction as APOE3/E3 microglia, but they did so at a consistently lower magnitude (Fig. 4c), indicating a possible genotype interaction, and consistent with our pathway enrichment analysis (Fig. 4b). To understand how DAB2 knockdown in iPSC-microglia correlated with microglia in the human AD brain and their association with RELN-axis genes, we carried out a Spearman correlation gene overlap analysis between RELN-coreDAB2 enrichment scores with microglial states in APOE iPSC-microglia and human AD brain microglia (AD Progression Atlas^29^), AD microglia from APOE4 and APOE3 carriers in the prefrontal cortex and lipid-laden APOE iPSC-microglia from Haney et al., (2024)^24^). This showed that RELN-DAB2 axis scores were significantly correlated with activated amyloid-responsive (ARM, ***ρ*** = +0.79, **FDR=0.001), with homeostatic signatures (***ρ*** = +0.62, *FDR=0.011; ***ρ*** = +0.58, *FDR=0.018) and with lipid-droplet accumulating microglial states (LDAM ***ρ***= +0.64, *FDR =0.011) in microglia in the human AD-afflicted brain with advancing pathology and across APOE iPSC-microglia; whereas the RELN-DAB2 axis showed no trend with the cytokine-responsive CRM-1 (***ρ***=-0.14, FDR=0.587) and CRM-2 states (***ρ***= –0.16, FDR = 0.587) (Fig. 4d). In contrast, DAB2-knockdown in APOE iPSC-microglia (dark blue dots) occupied the low RELN-DAB2 extreme of this association, with a depletion in activated/homeostatic programs, and providing loss-of-function evidence that DAB2 signalling sustains activated AD-relevant microglial states observed in the human brain (Fig. 4d). This was consistent with our above leading-edge analyses, which suggested that while microglial DAB2 is downregulated with increasing neuritic-plaque burden/CERAD scores in the human brain (Fig.1c) and particularly in APOE4/E4 carriers (Supplementary Fig. 1a), DAB2 expression was maintained in activated microglial signatures (n=7 carriers; Supplementary Fig. 1c). Exclusive to this correlation were the microglia identified from the prefrontal cortex of APOE carriers with AD (green symbols) (control n=12, AD APOE3/E3 n=7, AD APOE4/E4 n=9 individuals) and the APOE4/E4 lipid-laden iPSC-microglia (yellow triangles) (APOE3/E3 n=47, APOE4/E4 n=38 iPSC-lipid bearing microglia)^24^(Fig. 4d). These signatures may not have been detected in the AD progression atlas human brain studies^29^ because of differences in brain-regions profiled (the prefrontal cortex is relatively spared in pathology), or because lipid-laden microglia comprise a novel subtype under-represented in the AD spatial atlas given the low number of APOE4/E4 carriers in that study (n=7 carriers), or because of differences in microglial isolation (single-nuclear^29^ vs single-cell RNA^24^). Consistent with this, microglia from the prefrontal cortex from the AD progression atlas showed the weakest enrichment in RELN-DAB2 signalling of the four brain regions profiled (arrows in Fig. 4d). In addition the APOE4/E4 compared with APOE3/E3 contrasts were enriched in RELN-DAB2, ARM, and LDAM states but were depleted in homeostatic signatures (blue triangle shown by asterisks in Fig. 4d).

**Figure 4.**
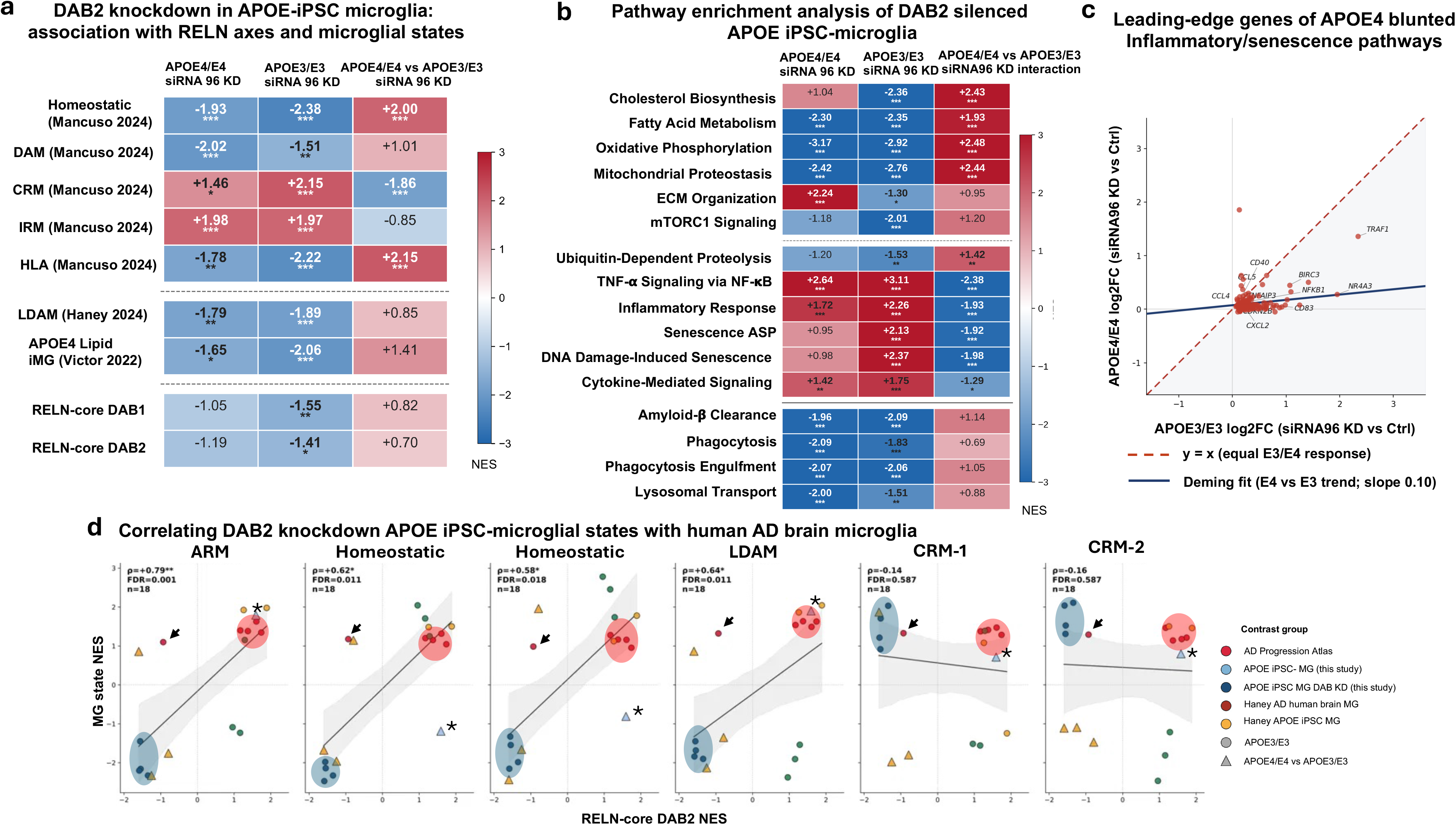
DAB2 knockdown alters microglial phagocytic and lipid pathways, reduces the RELN axis together with activated/homeostatic microglial states and pushes microglia towards cytokine-responsive states. **a**) RELN-axis and microglial AD-state enrichment following DAB2 knockdown. Heatmap of multilevel fgsea (infinite permutations) NES for microglial Alzheimer’s-disease (AD) state gene sets and (bottom rows) RELN-axis sets (RELNcore+DAB1 n=115, RELNcore+DAB2 n=113) across the same DAB2KD vs Ctrl contrasts (within APOE4/E4, APOE3/E3 genotypes and contrasting APOE4/E4 vs APOE3/E3 genotypes; columns as in a). AD microglial states reference sets were obtained from Mancuso et al., 2024^26^ (Homeostatic, Disease-associated (DAM), Cytokine-responsive (CRM), Interferon responsive (IRM), antigen presenting (HLA)), Haney et al., 2024^24^, Lipid-accumulating microglia (LDAM) and APOE4 Lipid-laden iPSC-microglia (Victor et al., 2022^48^). Fill = NES (±3); BH padj *≤0.05, **p ≤0.01, ***p ≤0.0001. b) Heatmap of normalised enrichment scores (NES) from pre-ranked pathway enrichment (fgsea multilevel; gene sets from GO:BP,KEGG,Reactome and MSigDB Hallmark) on the DESeq2 Wald-z gene ranking of DAB2-knockdown vs Ctrl-siRNA iPSC-microglia. DAB2 was silenced with siRNA96 targeting both DAB2-201 and DAB2-202 isoforms, versus Ctrl siRNA (n=4 differentiations per genotype), in APOE4/E4 (96siRNA n=3 differentiations) and APOE3/E3 lines (96siRNAn= 3 differentiations). Columns: DAB2-KD vs Ctrl (within APOE4/E4, within APOE3/E3, and genotype x siRNA96 interaction in APOE4/E4 versus APOE3/3 iPSC-microglia (positive NES indicates the knock-down response is weaker in APOE4/E4 vs E3/E3 (n=3 per genotype). Fill = NES (scale ±3). fgseaMultilevel infinite permutations). BH padj *≤0.05, **p ≤0.01, ***p ≤0.0001. **c) The transcriptional response to DAB2 knockdown in inflammatory and senescence-associated leading-edge genes is blunted in APOE4/E4 iPSC-microglia.** Plotting of per gene log2 foldchange (DAB2 siRNA-96 vs control) for the leading-edge genes of inflammatory and senescent pathway pathways (n=254 genes; TNFα-NFκβ, inflammatory response, DNA-damage/telomere stress senescence, cytokine-mediated signalling, SASP) plotted for APOE4/E4 vs APOE3/E3 iPSC-microglia. Red dashed line shows the hypothetical perfect linear relationship between both genotypes, with the blue line showing the Deming (orthogonal regression) slope of this correlation (slope= 0.10), with DAB2 knockdown in APOE4/E4 iPSC-microglia showing 10% of the APOE3/E3 microglial response ie an ∼90% blunting. Labelled points are the most genotype-divergent genes. **d) Correlation of DAB2-silenced APOE iPSC-microglia with microglia from the Alzheimer’s Disease afflicted human brain.** Spearman correlation (n=20 contrasts per panel) between RELN-core-DAB2 (113 genes) NES (x-axis) and microglial state NES (y-axis), across BIONIO37A/A-4 iPSC-microglia (Ctrl siRNA, light blue dots; DAB2 knockdown, dark blue dots), human microglia isolated from 4 brain regions of the AD progression atlas (Wachter et al., 2024^29^)(bright red dots); human microglia from APOE3/E3 and APOE4/E4 Advanced vs Early stages of AD in the prefrontal cortex (Haney et al., 2024^24^)(green); and lipid-laden iPSC-APOE microglia (Haney et al., 2024^24^) (yellow). Triangles show APOE4/E4 vs APOE3/E3 contrasts; circles mark all others. The six microglial states are amyloid/activated-response microglia (ARM; Sala-Frigerio et al., 2019^45^), homeostatic (consensus, Mancuso et al., 2024^26^), homeostatic (de Mathys et al., 2024^106^), lipid-droplet-accumulating (LDAM; Haney et al., 2024^24^), and cytokine-responsive (CRM-1/CRM-2; Mancuso et al., 2024^26^). RELN-axis NES scores correlate positively with ARM (***ρ*** = +0.79, ***FDR=0.001), both homeostatic signatures (***ρ*** = +0.62, *FDR=0.011 and ***ρ*** = +0.58, *FDR=0.018) and LDAM (***ρ***= +0.64, *FDR=0.011) with no trend observed with the cytokine-responsive CRM-1 (***ρ***=-0.14, FDR=0.587) and CRM-2 states (***ρ***= –0.16, FDR = 0.587). DAB2-knockdown microglia (dark blue dots) show coordinately reduced RELN-axis and activated/homeostatic states and increased pro-inflammatory cytokine-responsive (CRM1/CRM2) scores. Brain regions from the AD Progression atlas: prefrontal cortex, inferior temporal gyrus, entorhinal cortex, visual areas 1 & 2.

### DAB1 siRNA knockdown in APOE iPSC-differentiated microglia results in a depletion of HLA/IRM microglia-associated pathways

In contrast to DAB2, DAB1 knockdown (KD) targeted with two siRNAs (893, 894) yielded few differentially expressed genes, consistent with low DAB1 expression in iPSC-microglia and with only a trend downregulation in DAB1 following knockdown (Supplementary Fig. 6a-b) (siRNA-893, p=0.054; siRNA-894, p=0.056). Convergent DAB1 effects across genotypes upregulated RNA splicing/methylation and downregulated mitochondrial ATP generation and oxidative phosphorylation pathways (Hallmark OxPhos NES –1.62 to –2.55, p=3.2×10^−4^) (Supplementary Table 4). Filtering for microglial/immune categories, DAB1 knockdown, like DAB2, resulted in a depletion of transcriptional networks comprising phagocytosis, amyloid clearance, lysosomal transport as well as the complement pathway required for synaptic pruning but concurrently upregulated TNFα-NFκβ, pro-inflammatory and senescence responses (Supplementary Fig. 6c). DAB1 showed a strong transcriptional concordance with DAB2-modulated pathways (Spearman rho coefficient = 0.762) (Supplementary Fig. 6d), and we also found a significant DAB1KD-APOE4 interaction that showed blunted downregulation of mTORC1 signalling and a blunted response in pro-inflammatory TNFα-NFκβ responses and senescent pathways in APOE4/E4 microglia. DAB1 thus shares a pathway-level trend with DAB2 knockdown, but at a lower magnitude; Spearman correlation of NES (DAB1-KD vs DAB2-KD, pooled APOE genotypes) indicated that while DAB1 knockdown showed coordinated changes in endolysosomal pathways this did not recapitulate significant downregulation of endolysosomal, amyloid beta processing and phagocytic genes at the gene-level like DAB2 knock-down (Supplementary Fig. 6e). While there was strict transcriptional concordance between DAB1 and DAB2 knockdown, direct comparison of NES scores suggested that DAB1 knockdown was significantly more enriched in interferon-responsive and LXR cholesterol efflux microglial states, whereas DAB2 knockdown was more strongly enriched for transcriptional networks associated with transitioning cytokine-responsive CRM and disease-associated (DAM) states (Supplementary Fig. 6f-g), although larger studies are needed to confirm this. Thus, under basal conditions DAB1 has modest effects on pathways largely concordant with DAB2.

### Chronic RELN deficiency is associated with altered microglial morphology and CD68 upregulation in the postnatal mouse cortex

To assess how RELN signalling impacts microglia in vivo, we assessed for microglial changes in a germline Reln-knockout mouse (Supplementary Fig. 7). We first carried out an expression analysis of RELN in relation to IBA1+ immunostained or CX3CR1-Gfp genetically traced microglia across development until postnatal stages, given that these mice do not survive into adulthood. This showed that RELN proteins, but not transcripts, localised to IBA1+ or CX3CR1-GFP microglia in the embryonic (E14.5) and postnatal (P0.5) cerebral cortex (Fig. 5a), and that *Vldlr* and *Apoer2/Lrp8* transcripts were detected in Cd11b+ microglia (Supplementary Fig. 7a), consistent with microglia competent to respond to RELN signalling and with our human brain microglial transcriptomic and iPSC-microglial data. We next quantified the total number and distribution of microglia to investigate if Reln-deficiency impairs microglial recruitment into the brain. This showed that microglial density was unchanged between Reln-KO and WT mice at E14.5 and P22 (Supplementary Fig. 7f). However, at P22 Reln-KO cortical microglia were hypotrophic, with significantly reduced process length (Process Length: WT 268.0+/−26.9 um, n=5 mice; KO 152.3+/−12.2, n=4; p=0.009) and branch level (WT 4.14+/−0.13; KO 3.53+/−0.15; p=0.018; Fig. 5b), without changes in surface area or volume (Supplementary Fig. 7g). As these morphological changes are reminiscent of microglia changing from homeostatic surveillant to activated states^56^, we next asked if microglia were more activated based on their expression of the endolysosomal marker CD68 which is associated with phagocytic microglia^57^ and is elevated in the human AD-brain^58,59^. This showed that hypotrophied microglia in the Reln-deficient P22 cortex was accompanied by their increased CD68^+^ lysosomal volume (WT 11.5+/−1.8; KO 25.7+/−3.9; p=0.038; Fig. 5c), with similar hippocampal trends (Supplementary Fig. 7h-i). As these mice exhibit severe defects in cortical lamination and neuronal positioning (Supplementary Fig. 7d-e), we cannot exclude the possibility that the observed microglial deficits are secondary to disrupted Reln-dependent neuronal signalling^1,60^. Nonetheless, our observations are consistent with chronic RELN deficiency being associated with microglial altered morphologies and their increased activation based on their upregulation of CD68. Together, this suggests the altered endolysosomal/phagocytic function independent of microglial cell number, and consistent with endolysosomal/phagocytic transcriptomic pathways dysregulated in our DAB1 and DAB2 knockdown APOE iPSC-microglial findings.

**Figure 5.**
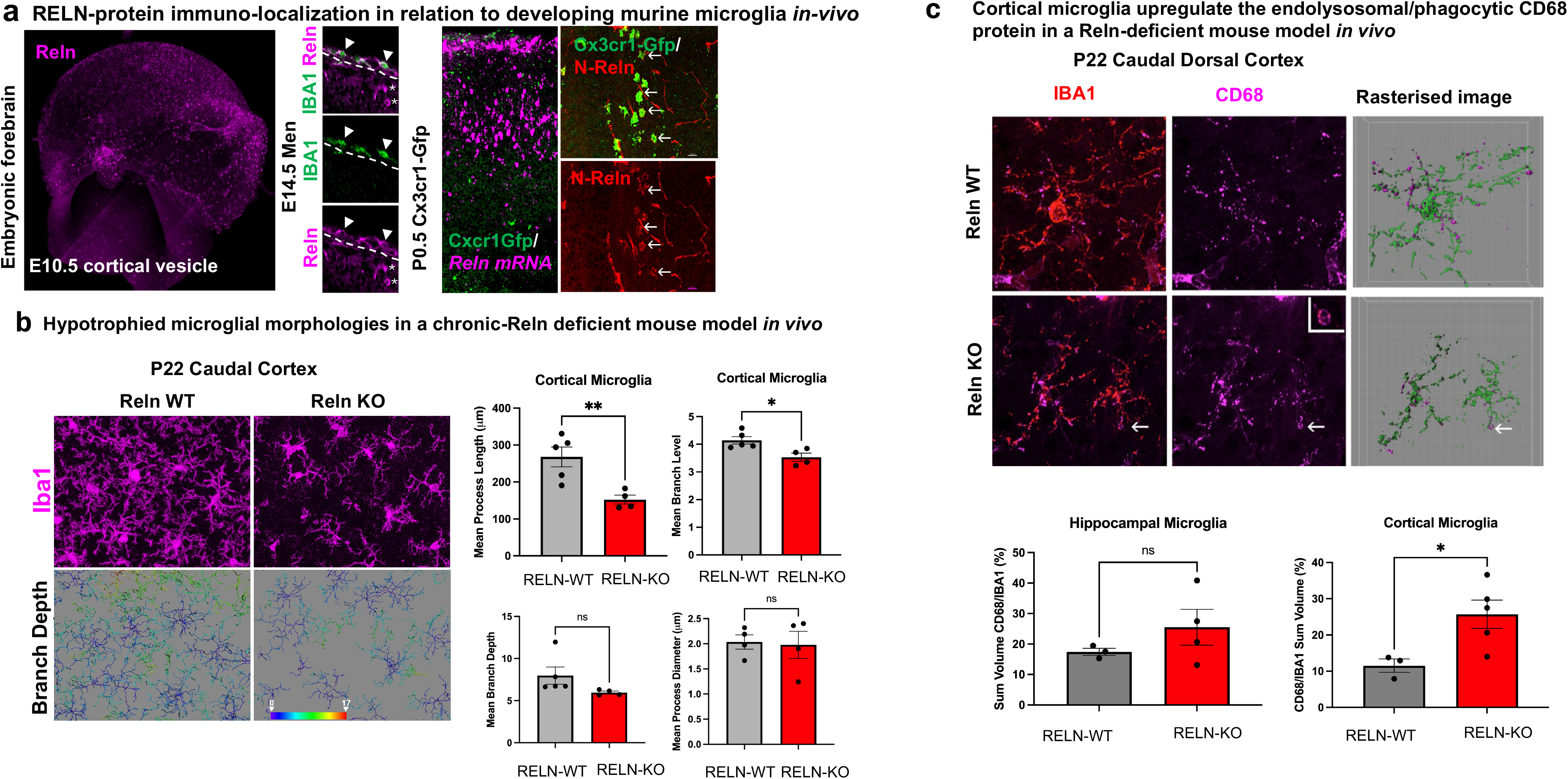
Analysis of microglia in a chronic-Reelin deficient mouse model results in the altered microglial morphologies and in their upregulation of endolysosomal protein CD68 in the P22 cortex. **a**) Whole-mount immunostaining of E10.5 mouse wildtype embryonic vesicle for RELN N-terminal proteins (mG10 antibody), with middle panels showing high magnification through the marginal zone of the E14.5 mouse cortex co-labelled for RELN N-terminal proteins and the microglial/macrophage marker IBA1. RELN proteins colocalise with IBA1+ microglia located within the pia/meninges at the border with the marginal zone (dashed white line), with RELN-expressing Cajal-Retzius cells denoted by the asterixes. Right panels show *in situ* hybridization of *Reln* transcripts (magenta) in a P0.5 CX3CR1-Gfp mouse cortex with microglia genetically traced with Gfp (green) showing *Reln* transcripts are not expressed by microglia. b) High magnification (x40) of the caudal dorsal-lateral cortex of P22 ActinCre+/−;Reln+/+ (Reln-WT, left) and ActinCre+/−;Relnfl/fl (Reln-KO, right) mice, immunostained for the ionised calcium binding protein IBA1 (magenta) expressed by microglia and macrophages. Lower panels show filamentous rendering of microglia reconstructed with Imaris software and colour coded according to their degree of process branching. Graphs show mean values ± SEM, with dots denoting number of mice analysed (n= total of 806-831 cells from n=4-5 mice Reln-WT, n= 559-801 cells from n=4 mice Reln-KO). (*p = 0.0184, **p = 0.0089 Unpaired T-Test). c) Indications of increased phagocytosis of cortical microglia in P22 Reln deficient mice. Panels show high magnification of representative microglia residing in the caudal dorsal-lateral cortex of P22 ActinCre+/− Reln+/+ (Reln-WT, top) and ActinCre+/−;Relnfl/fl (Reln-KO, bottom) mice, co-immunostained for the ionised calcium binding protein Iba1 (red) and the lysosomal marker CD68 (magenta). Right panels show 3D surface reconstruction of CD68+ structures (magenta) localised within each Iba1+ microglia (green). Graphs show mean values ± SEM of the percentage (%) volume of CD68+ structures per hippocampal (left graph) and cortical (right graph) IBA1+ microglial sum volumes, quantified from surface reconstructions, with each dot corresponding to a single mouse (27 cortical microglia from n=3 Reln-WT, 48 cortical microglia from n=5 Reln-KO; 9 hippocampal microglia from n=3 Reln-WT, 18 microglia from n=5 Reln-KO)(*p = 0.0381, Unpaired T-test).

### DAB2 knockdown is negatively associated with TREM2-DAM-associated lysosomal/phagocytic networks in microglia

To identify overarching transcriptional relationships between our experimental perturbations and published microglial-state signatures, we performed principal component analysis (PCA) on our fast gene set enrichment analysis (fgsea) studies, based on normalised enrichment scores (NES) computed across all contrasts (APOE genotype, DAB1/DAB2 knockdown, published lipid/amyloid iPSC-microglia, developmental subtypes and human AD microglia) against a panel of ten independent microglial-state signatures (Fig. 6, Supplementary Table 5). Placing the pathway-enrichment profiles of our DAB1 and DAB2 knockdown and APOE-isogenic iPSC-microglia alongside these independent microglial signatures from the human AD brain and from other APOE4 iPSC-microglial studies on a common set of summary axes, allowed us to map our APOE iPSC-microglial experiments onto a human microglial AD brain landscape. As expected, the RELN-DAB1 and RELN-DAB2 axes (green arrow heads) pointed opposite to the DAB1/DAB2-knockdown iPSC-microglia (open and closed blue circles), which occupied the cytokine-responsive microglial states space. Principal component analysis 1 showed that the x-axis (PC1) accounted for 46.7% of variance and separated activated DAM/LDAM and APOE4 programmes (positive) from cytokine-responsive states (negative; Fig. 6a-b) (MG-DAM +0.41, LXR cholesterol efflux +0.41, APOE4-lipid +0.40, LDAM +0.37, MG-HLA +0.35, RELN-DAB1 +0.27, MG-IRM +0.26, RELN-DAB2 +0.22, MG-HM +0.15, MG-CRM –0.15). Principal component two (PC2) ie the y-axis explained 19.7% of variance and resolved interferon/cytokine states (CRM –0.29, IRM –0.36) from the RELN-signalling axes (RELN-DAB2 +0.56, RELN-DAB1 +0.52), the latter which modestly co-loaded with homeostatic microglia (HM +0.29) (Fig. 6a,c). Accordingly, microglia isolated from the human brain with increasing AD-pathology (red circles) occupied the positive PC1 DAM/lipid pole, and APOE4/E4 iPSC-microglia mapped toward the same DAM/LDAM/IRM programmes (Fig. 6a), whereas DAB1– and DAB2-deficient APOE iPSC-microglia sat in the opposite cytokine-responsive space (blue circles). Not all datasets were uniform: microglia isolated from the human AD-afflicted prefrontal cortex (Haney et al., (2024)^24^ single-nucleus, red triangles) occupied a negative-PC1 and high PC-2 space closer to homeostatic/developmental microglia, than to the activated pole (APOE3/E3-AD PC1-0.42, PC2 +1.67; APOE4/E4-AD-0.36, +2.85; APOE4/E4-AD vs APOE3/E3-AD PC1-1.27; PC2 +2.93); and APOE4-lipid laden iPSC-microglia (yellow squares) spanned the CRM-DAM boundary (APOE4/E4 vs APOE3/E3 microglia PC1-0.26, PC2-1.03 and APOE4/E4 vs APOE3/E3 lipid-droplet microglia PC1-1.08, PC2-1.38, while lipid droplet microglia vs lipid-droplet free microglia sat at the DAM pole (APOE3/E3 +2.71, +0.80; APOE4/E4 +2.53, +0.10), likely reflecting region (relatively spared prefrontal cortex) and platform differences (single nucleus under-capture of activated/lipid states). Consistent with this, the prefrontal cortex contrast was also the least activated with the AD Progression Atlas itself (PC1 +0.56 versus inferior temporal gyrus +3.49). Homeostatic states also co-loaded with increasing developmental maturation (green triangles) on PC3 (MG-HM +0.58) (Supplementary Fig. 8), with postnatal/adult mouse microglia. Overall, we propose that DAB2 cell-autonomously primes microglia toward homeostatic and protective DAM/activated states with its deficiency pushing microglia toward pro-inflammatory and cytokine-responsive states, positioning microglial DAB2-deficient signalling as a convergent driver of maladaptive AD-relevant microglial states.

**Figure 6.**
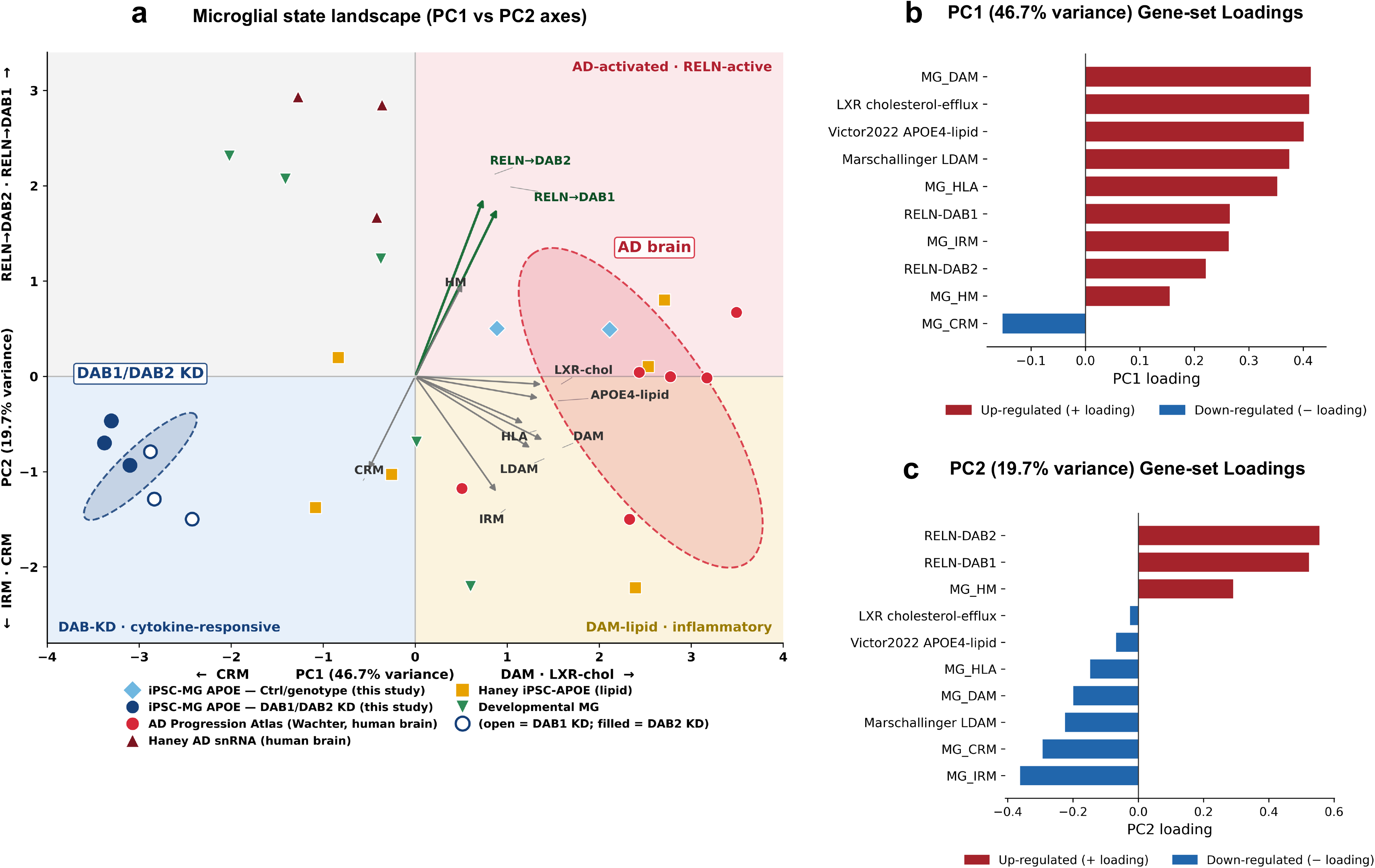
Mapping APOE iPSC-microglia on the APOE4/AD-associated human microglial landscape in the human brain shows DAB2 knockdown microglia are transcriptionally depleted in RELN-activated DAM/LDAM/HM/HLA states and instead transcriptionally resemble cytokine-responsive microglial states observed in the human AD brain. **a**) Principal component analysis (PCA) of the normalized enrichment score (NES) matrix (human microglial reference states and RELN-axis gene sets contrasted with 28 experimental datasets from DeSEQ2 analyses in 6 experimental groups: including APOE-iPSC microglia (BIONIO37A/A4, light blue diamonds) and their DAB1/DAB2 siRNA knockdown (blue circles); 10 human microglial gene sets i.e. lipid-laden iPSC-APOE microglia (Haney et al., 2024^24^, orange squares), human AD microglia, late vs early CERAD pathology from the AD Progression Atlas (Wachter et al., 2024^29^; red circles) or from pre-frontal cortical microglia at advanced compared with early stages of AD (Haney and others 2024^24^, red triangles) or from ageing microglia (Galatro et al., 2017^28^, green triangles) or early fetal and postnatal human (Kracht et al., 2020^27^ early GW9 vs late GW12-15; Mouse Ostrem et al., 2024^124^ (P14 vs E13, P14 vs E15). Red arrows show gene-set loadings tracking homeostatic to AD-relevant activated DAM (+0.41), LDAM (+0.37), APOE4-lipid (+0.40), HLA (+0.35), IRM (+0.26) and homeostatic (+0.15), microglial states, or towards pro-inflammatory cytokine responsive (CRM, –0.15) states, with RELN-core+DAB1 (+0.27) and +DAB2 (+0.22) loadings shown by green arrows that track the opposite pole to CRM. Quadrant labels denote the activated-AD/RELN-active microglial pole (PC1>0) versus the cytokine and DAB-knockdown poles (PC1<0). PC1= 46.7%, PC2= 19.7% (cumulative = 66.4%). Dotted line is the hypothetical interaction between DAB2 KD and APOE4 in reversing APOE4/AD-associated states. **b)** Bar plots show PC1 loadings contributing to 46.7% variance. PC1 is driven positively (red bars) by activated phagocytic and lipid-accumulating microglia (DAM (+0.41), LDAM (+0.37)) and APOE4-lipid (+0.40), LXR cholesterol efflux (+0.41) and antigen-presenting (HLA +0.35) states, and negatively only by the cytokine-responsive microglia (blue bars) ie PC1 separates CRM1 (–0.15) from all other states. c) PC2 loadings (19.7% variance). PC2 is led positively by the RELN axes (RELNcoreDAB1 (+0.52), RELNcoreDAB2(+0.56) and homeostatic microglia(+0.29) (red bars), and negatively by the activated/inflammatory states (APOE4-lipid (–0.07), HLA (–0.15), DAM (–0.20), LDAM (–0.22), CRM (–0.29), IRM(–0.36) (blue bars) – the RELN axes of variation. DAB2-knockdown contrasts map to the developmental/ageing pole with CRM microglial states, indicating RELN-axis activity is inversely associated with the pro-inflammatory state program and a shift of DAB2 KD microglia towards CRM states. DAB2 knockdown iPSC-microglia correlated modestly with postnatal mouse microglial maturation (P14 vs embryonic; Spearmean **ρ** ∼+0.3-0.38), but this reflected a shared loss in DAM/lipid activation programmes. Homeostatic signatures are positively correlated both maturation and ageing, and DAB1/DAB2 knockdown moved microglia in the opposite direction.

To resolve how DAB2 reshapes microglial states at the network level, we performed weighted gene co-expression network analysis (WGCNA) on the top 10,000 most variable genes following DAB2-knockdown in APOE iPSC-microglia (Fig. 7, Supplementary Table 6). This yielded fourteen co-expression modules including two modules positively associated with DAB2 knockdown (brown, n=927 genes; green, n=589 genes) and five negatively associated modules (Fig. 7a & Supplementary Figure 9a-b). The integrin receptor ITGB8 was a member of the positively associated brown module which was enriched in cytokine-responsive microglial signatures and in pro-inflammatory TNFα-NF-κβ pathways (Fig. 7a-c, Supplementary Fig. 9c). This showed enrichment of several chemokine receptors (CCR5, CCR7, CCR2), chemokine ligands (CCL2, CXCL3, CXCL8, CCL20), TNF receptors (TNFRSF9, TNFRSF18), of colony stimulating factor (CSF3), interferons and their receptors (IFNL3, IFNAR2) and interleukins and their receptors (IL23A, IL17C, IL10RB) (Supplementary Table 6). The positively associated green module was also enriched for TNFα-NF-κβ pathways and circadian rhythms but did not correspond to a specific microglial state. The RELN/APOE receptor LRP8 was a hub gene of the negatively associated blue module and contained the LRP1 APOE receptor. This blue module (n= 1,001 genes) was associated with gene networks underlying resting and antigen-presenting HLA, homeostatic, resting and APOE4-lipid microglial states, and was downregulated by DAB2 knockdown (Fig. 7b & Supplementary Figure 9c), suggesting that DAB2 loss disrupts an LRP8-centered receptor network relevant to microglial state transitions relevant to AD. Pathway enrichment analysis for this module was dominated by ribosome biogenesis, chromosome organisation and translation (Fig. 7c). DAB2 was a member of the negatively associated purple module (n= 150 genes), which comprised gene networks underlying activated DAM, TREM2-LAM, TREM2-lipid, LDAM and LXR cholesterol-efflux-activated microglial states and was transcriptionally enriched in lipid-metabolic and phagocytic pathways (Fig.7b-c). Within the negatively-associated purple module, DAB2 was co-expressed with a coordinated phagocytic-endolysosomal programme: receptor-mediated phagocytosis (the scavenger receptors CD36, MSR1, TREM2; the complement receptor-3 subunits ITGAM and ITGB2, the inhibitory receptor CD300A, and the endosomal sorting receptor SORL1, lysosomal acidification (the V-ATPase accessory subunit ATP6AP2), and hydrolytic degradation (LIPA, GUSB, MAN2B1, TPP1 and the hydrolase co-factors (PSAP, SUMF1). Downregulation of this module with DAB2 deficiency therefore implicates impaired phagocytic uptake and endolysosomal degradation, including of amyloid-beta and complement-tagged synaptic proteins (Supplementary Table 6), and consistent with broader suppression of the lysosomal and phagocytic programmes seen across the DAB2-knockdown transcriptome (Supplementary Table 3). In addition, this module was associated with amyloid-beta degradation (ACE), clearance (CD36, MSR1, TREM2, ITGAM, ITGB2), and intracellular breakdown in lysosomes (TPP1)(Supplementary Table 6). The RELN receptor VLDLR and integrin receptor ITGB1 mapped to the turquoise module (n=1,514 genes) which was not associated with a specific microglial state. Other downregulated modules included the salmon homeostatic module (n= 101 genes) associated with innate-immune and endolysosomal pathways; and the magenta homeostatic module (n= 232 genes), associated with innate-immune/complement pathways (Supplementary Table 6). Together this indicates that DAB2 knockdown regulates gene-module networks implicating both RELN and APOE signalling components. To independently confirm that DAB2 regulated gene module was associated with DAM/TREM2 states we carried out Fisher’s exact gene overlap tests between the WGCNA DAB2 siRNA knockdown co-expression gene modules with independent human^61–64^ and mouse^62,65^ transcriptomic signatures. This confirmed that the DAB2 purple module correlated with TREM2-DAM signatures, but not with TREM2-independent homeostatic, pro-inflammatory cytokine and interferon responsive or dystrophic states (Supplementary Fig. 9e). Comparison of gene modules associated with microglial transcriptomes across increasing CERAD or Braak stages (Supplementary Fig. 9d) however showed no significant association of gene modules with brain-region specific pathology. At the protein level, TREM2 was reduced by DAB2 knockdown in APOE3/E3 but not APOE4/E4 microglia (Fig. 7d), consistent with our enrichment studies suggesting that APOE4 blunts the response to DAB2 knockdown within an APOE4/E4 genetic background and with our pathway network analysis showing DAB2 regulates gene networks associated with a TREM2/DAM response (purple module). To test whether these modules overlapped with dementia-risk loci, they were contrasted with a panel of 91 genes comprising 35 known-locus genes and 55 prioritised genes from the 75 genome-wide significant loci of Bellenguez et al., (2022)^66^, plus APOE, of which 37 were present in the 10,000-gene WGCNA network. This showed that the purple/DAM/TREM2 module was enriched 7.2-fold (4/150 genes) consisting of lipid/endolysosomal dementia-risk genes TREM2 (kME 0.9, lead), GRN (0.84), SORL1 (0.66), ACE (0.71) (hypergeometric test p=0.002, q_BH=0.033) and the (salmon) homeostatic module enriched 8^−^ fold with complement/membrane innate-immune risk genes (3/101 genes) (CR1, kME 0.84), and the MS4A4A gene which has been shown to negatively regulate TREM2 (0.84), and also including the WAVE complex subunit ABI3 gene (0.64) (p=0.006, q_BH=0.045) which is an important activator of the Arp2/3 complex important for maintaining microglial morphological homeostatic surveillant states^57,67^ (Fig. 7e & Fig. 8, Supplementary Table 7). Collectively, these data establish DAB2 as a regulator of gene networks underlying microglial transcriptomic state transitions and of the endolysosomal, lipid-handling, phagocytic and complement pathways relevant to dementia-risk.

**Figure 7.**
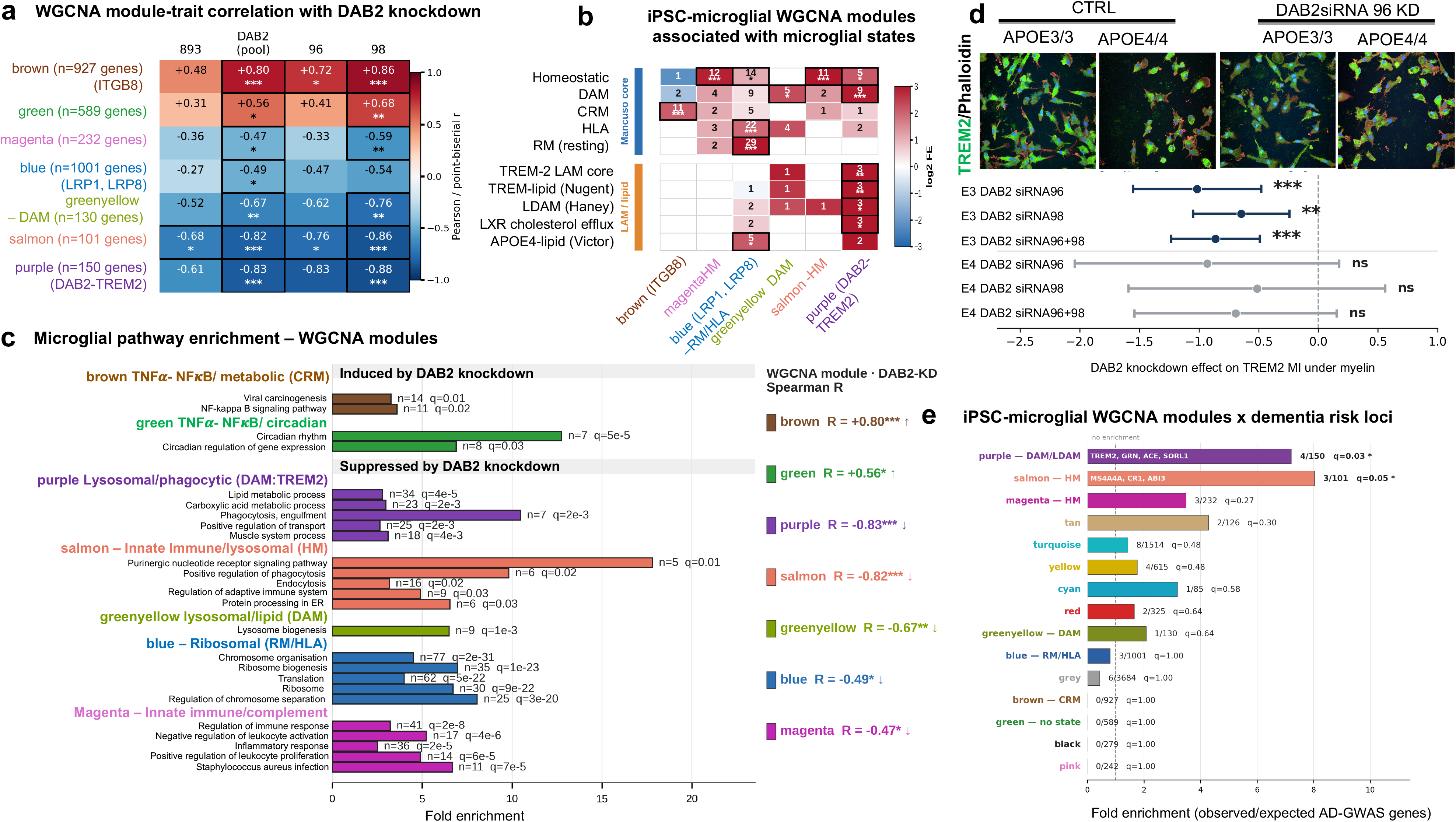
DAB2 knockdown remodels microglial-state co-expression modules in isogenic-APOE iPSC-microglia, elevating the cytokine-response (CRM, brown) module and suppressing the disease-associated/lipid (DAM/LDAM, purple; DAM, greenyellow) and homeostatic (HM, salmon) modules. **a**) WGCNA module–trait correlation with DAB2 knockdown. Heatmap of Pearson’s/point-biserial r between the eigengenes of the seven trait-significant iPSC-microglial WGCNA modules and DAB2-knockdown traits. WGCNA was performed on the top 10,000 most-variable genes of the variance-stabilised (VST; DESeq2 v1.46), limma-batch-corrected expression matrix of n = 39 BIONIO37A (APOE3/3) and BIONIO37A-4 (APOE4/4) iPSC-microglia. A soft-thresholding power β = 12 (scale-free-topology R² = 0.846) was applied to build a signed-hybrid weighted adjacency matrix; the signed Topological Overlap Measure was computed and genes clustered by the dynamic tree-cut algorithm (minModuleSize = 30, deepSplit = 2, mergeCutHeight = 0.25; single block). This yielded 14 modules plus grey, with 3,684 of 10,000 genes (36.8%) unassigned (grey). The seven trait-significant modules displayed are brown (cytokine-response, CRM; 927 genes), green (no discrete microglial state; 589), magenta (homeostatic, HM; 232), blue (ribosomal/antigen-presenting, RM/HLA; 1,001), greenyellow (disease-associated, DAM; 130), salmon (homeostatic, HM; 101) and purple (disease-associated/lipid-droplet-accumulating, DAM/LDAM; 150); microglial states were assigned by marker over-representation (panel c). Module membership (MM = signed kME) is the Pearson correlation between each gene and its module eigengene (the first principal component of the module’s expression across the 39 samples). Only trait columns reaching BH-significance in ≥1 module are displayed: the genotype-pooled DAB2 knockdown (siRNA-96 + siRNA-98, n = 14 vs control siRNA n = 8), the genotype-pooled siRNA-96 (n = 6) and siRNA-98 (n = 8) effects, and DAB1 siRNA-893. siRNA-96 targets both DAB2-201 and DAB2-202 isoforms whereas siRNA-98 targets DAB2-201 only, so effects convergent across both siRNAs implicate DAB2-201. DAB2 knockdown elevated the CRM (brown, r = +0.80) and green (r = +0.56) modules and suppressed the DAM/LDAM (purple, −0.83), DAM (greenyellow, −0.67), homeostatic (salmon −0.82; magenta −0.47) and RM/HLA (blue −0.49) modules, indicating a shift away from disease-associated/lipid and homeostatic programmes toward the cytokine-response state. The APOE-genotype and DAB1-pooled trait columns showed no significant module correlations and are omitted (full matrix in the supplementary figure). Stars: BH-adjusted *p < 0.05, **p < 0.01, ***p < 0.001. **b) Overlap of WGCNA-module genes with curated microglial-state signatures**. Heatmap of one-sided Fisher exact gene-overlap tests between each module (columns: brown, magenta, blue, greenyellow, salmon, purple) and curated microglial-state gene sets (rows); fill = log2 fold enrichment, the number in each cell = overlapping gene count, stars = BH-FDR (*p < 0.05, **p < 0.01, ***p < 0.001). States: homeostatic, disease-associated (DAM), cytokine-responsive (CRM), interferon-responsive (IRM), antigen-presenting (HLA) and ribosomal/resting (RM) (Mancuso et al., 2024^26^); TREM2-LAM core (Jaitin et al., 2019^129^); TREM2-lipid microglia (Nugent et al., 2020^44^); lipid-droplet-accumulating microglia (LDAM; Haney et al., 2024^24^; Marschallinger et al., 2020^47^); LXR cholesterol-efflux microglia; and APOE4-lipid iPSC-microglia (Victor et al., 2022^48^). Module-state assignments: brown = CRM, magenta and salmon = HM, blue = RM/HLA, greenyellow and purple = DAM (purple additionally LDAM/TREM2-LAM); green showed no significant marker over-representation. Universe = the network gene set. **c) Microglia/AD-relevant pathway enrichment per module.** Bars show fold enrichment from one-sided Fisher exact over-representation analysis of each module’s genes against a curated microglial/AD-relevant functional gene-set panel (GO Biological Process, KEGG and MSigDB Hallmark terms), filtered to BH-significant terms (q < 0.05, ≥ 4 overlapping module genes), with redundant terms collapsed by overlap-gene Jaccard > 0.4 and the top 6 per module shown; each bar is annotated with the overlapping module-gene count (n) and BH-adjusted p (q). Enrichment is shown for the brown (CRM), green, purple (DAM/TREM2), salmon (HM), blue (RM/HLA), greenyellow (DAM) and magenta (Innate Immune/Complement) modules. **d) TREM2 is downregulated by DAB2 siRNA knockdown in myelin-treated APOE iPSC-microglia.** APOE3/3 and APOE4/4 iPSC-microglia (BIONIO37A/A-4) immuno-stained for TREM2 (green) and actin-labelled with phalloidin (red) in CtrlsiRNA and DAB2siRNA96 knockdown of both DAB2-201/-202 isoforms. Forest plot showing mean-intensity z-scores for: APOE3/E3 DAB2 siRNA-96 knockdown (9 wells, 2,246 cells, 3 experiments) vs Ctrl siRNA (n=3 independent experiments, 11 wells, 4,335 cells); DAB2 siRNA-98 knockdown (n=3 experiments, 9 wells, 4,979 cells) vs Ctrl siRNA (n=3 experiments, 11 wells, 4,335 cells); DAB2 siRNA-96+98 knockdown (n=3 experiments, 18 wells, 7,225 cells) vs Ctrl siRNA (n=3 experiments, 11 wells, 4,335 cells); APOE4/E4 DAB2 siRNA-96 (n=4 experiments, 13 wells, 4,571 cells) vs Ctrl siRNA (n=4 experiments, 15 wells, 5,494 cells); DAB2 siRNA-98 knockdown (n=4 experiments, 15 wells, 5,203 cells) vs Ctrl siRNA (n=4 experiments, 15 wells, 5,494 cells); DAB2 siRNA-96+98 (n=4 experiments, 28 wells, 9,774 cells) vs Ctrl siRNA (n=4 experiments, 15 wells, 5,494 cells). Negative z-scores indicate reduced TREM2 with knockdown; significance from a linear mixed model (value ∼ siRNA + (1|experiment)) with BH-FDR (**FDR<0.01, ***FDR<0.001). **e) The DAM/LDAM (purple) and homeostatic (HM, salmon) WGCN modules of isogenic APOE iPSC-microglia are enriched for Alzheimer’s Disease genetic risk factors (Bellenguez et al., 2022**^66^). Bars show the fold-enrichment of the overlap between each WGCNA module and the 75 dementia-risk gene reference list from Bellenguez et al., 2022^66^ which were detected within the 10,000 WGCN gene network. The reference list comprised the 35 lead genes at the 33 known loci (Supplementary Table 5 from Bellenguez et al., 2022)^66^ and the 55 Tier 1 and Tier 2 prioritised genes at the 42 new loci (Supplementary Table 20 from Bellenguez et al., 2022)^125^, plus APOE, = 91 genes in total, of which 37 were present in the 10,000 WGCNA gene network tested. Fisher’s Exact test, BH padj values *q<0.05. Purple/DAM module AD-risk genes: TREM2 (kME 0.9, lead), GRN (0.84), SORL1 (0.66), ACE (0.71)(7.2x enrichment (hypergeometric p=0.0022, q_BH=0.033, 7.21x enrichment); and the salmon/homeostatic module:CR1 (0.84), MS4A4A (0.84), ABI3 (0.64) (8.0x enrichment, p=0.006, q_BH=0.045).

**Figure 8.**
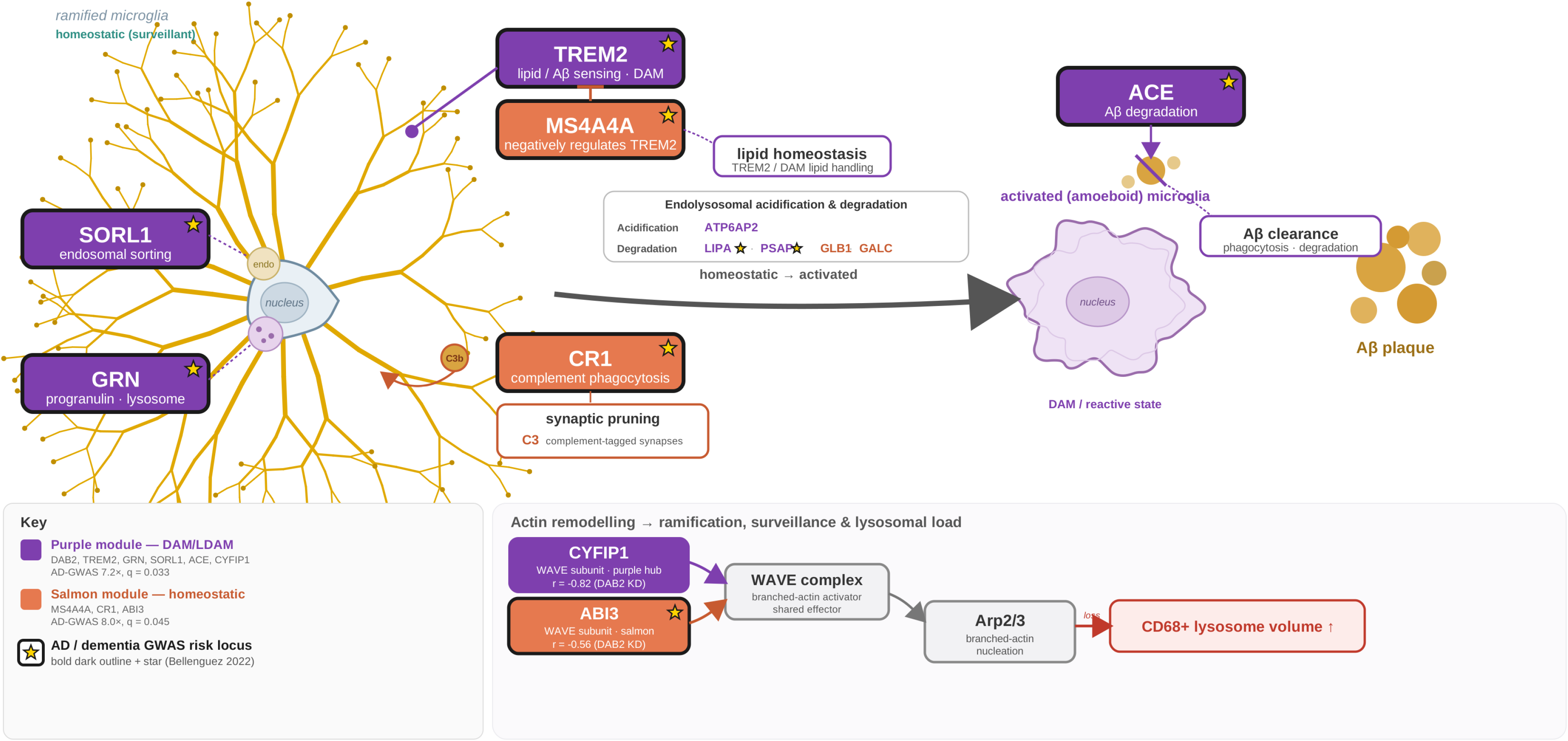
DAB2 knockdown remodels microglial-state co-expression modules associated with dementia-risk loci. Schematic illustrating pathway analysis of DAB2-knockdown negatively associated purple (n=150 genes) and salmon (n=101 genes) gene modules associated with microglial endosomal sorting (SORL1^130,131^), endolysosomal homeostasis (GRN^132–134^), endolysosomal acidification (ATP6AP2; GO:0007035), endolysosomal organisation (LIPA, PSAP, GLB1; GO:0007040) and degradation (GALC; GO:0004336), TREM2/DAM lipid handling and activation of DAM states (TREM2^135,136^, MS4A4A^137^), synaptic pruning (CR1^20,138^), amyloid-beta degradation (ACE^139,140^) and actin remodelling important for microglial morphological changes from their homeostatic surveillance states to more activated states which are coupled with lysosomal load (ABI3, CYFIP1^57^). Genes which correspond to the 75 identified dementia-risk loci (Bellenguez et al., 2022^66^) identified in gene modules and pathway analysis are denoted by a star and black outline.

## Discussion

APOE4 is the greatest genetic risk factor for sporadic AD^68^ that partly increases AD-risk through its exacerbation of amyloid pathology^11^ and impaired microglial function^24,48,54,69^, yet the receptor-adaptor machinery that couples APOE to microglial state transitions in the human brain remains poorly defined. Here, we identify the endocytic adaptor DAB2 as a cell-autonomous regulator of microglial states associated with dementia risk, and a candidate point of convergence between RELN and APOE AD-risk pathways^8^. In isogenic iPSC-microglia, acute loss of DAB2 depleted the gene networks that define disease-associated/TREM2 (DAM), lipid-droplet accumulation (LDAM), and homeostatic states, and shifted microglia toward cytokine-responsive (CRM), and TNFα-NFκ-β-driven states. The DAB2-containing DAM/LDAM co-expression module negatively associated with DAB2 knockdown was enriched for known lipid and endolysosomal dementia-risk genes (TREM2, GRN, SORL1, ACE), and a second homeostatic (salmon) module suppressed by DAB2-knockdown was enriched for complement and membrane innate-immune risk genes (CR1, MS4A4A, ABI3). In the human brain, microglial DAB2 declined with amyloid burden in APOE4/E4 carriers, while correlating positively with a DAM/TREM2 signature across donors, suggesting that DAB2 promotes and maintains DAM/TREM2 microglial states, considered to be protective in early AD^46,62,64^. Together, these findings place DAB2-regulated microglial transcriptomic states directly on the landscape of human AD genetic risk.

Here we identify DAB2 as an APOE4-sensitive receptor-adaptor machinery that promotes DAM/TREM2 disease-associated states, corresponding to gene modules enriched in dementia risk loci that we predict are protective at early stages of AD. Our network (WGCNA) analysis in APOE iPSC-microglia placed DAB2 directly within the human DAM/TREM2 program. The DAB2-containing (purple) module was associated with endolysosomal, phagocytic, neuronal crosstalk, and amyloid-beta processing pathways and with DAM/TREM2 phagocytic microglial states enriched for DAM and lipid-handling genes including TREM2, GRN, SPP1, PSAP and MSR1, and was depleted by DAB2 knockdown. Tested against independently defined human microglial state signatures from the human and mouse brain^61–64^, this module was specifically and strongly enriched in the DAM/Neurodegenerative disease (MGnD) state, with TREM2, SPP1 and CD63 among the overlapping genes, but did not overlap with homeostatic, interferon or cytokine-responsive states, providing cross-dataset support that the DAB2 module supports the DAM/TREM2 program rather than this comprising a culture-specific signature (Supplementary Fig. 9e). TREM2 was the most central member of this module (module membership kME =0.9, exceeding DAB2’s own 0.71), and TREM2 protein was decreased following DAB2 knockdown in APOE3/E3 microglia. Because the DAM transition is thought to be TREM2-dependent^26,46^ and because APOE, upregulated in both our iPSC-microglia and in the AD brain, is a TREM2 ligand, our work positions DAB2 as a regulator of the APOE-TREM2 axis that cell-autonomously ^62^governs and maintains DAM acquisition. Loss of DAB2 lowers TREM2 and depletes the DAM/TREM2 network it helps sustain. Together, these findings place DAB2 as a central node in the AD-genetic risk landscape.

Several lines of evidence point to a dual role of DAB2 in restraining inflammation while promoting microglial activated states. An anti-inflammatory role for microglial DAB2 is consistent with its function in peripheral monocytes/macrophages, where DAB2 binds the TNF receptor-associated factor 6 (TRAF6) to restrain NF-κβ nuclear translocation and signalling, with myeloid DAB2 deficiency worsening inflammation^70^. In addition, DAB2 overexpression in the amyloid APP/PS1 mouse model, stabilised TGFßR2 to activate SMAD2/3, attenuating microglial pro-inflammatory mediators IL1ß and TNFα, and protected against amyloid deposition, hippocampal neuronal loss and memory impairment^18^. DAB2’s effects however may be region– or context-dependent; in an experimental model of autoimmune encephalomyelitis, stress-induced DAB2 in Cd11b+ microglia/macrophages and in astrocytes exacerbated inflammation^71^. Together with our in-vitro studies, this suggests that DAB2 puts a brake on inflammation while supporting microglial activation. Notably, the TREM2-DAB2-regulated DAM/LDAM network was enriched for human dementia-risk loci. These included GRN, previously shown to suppress DAM and neurodegenerative microglial signatures in mice (APOE, Clec7a, CD68 and Ccl4) and to promote homeostatic signatures in an opposing direction to TREM2^72^. The co-membership of these functionally opposing genes suggests that DAB2-associated modules capture the regulatory balance of DAM transitions associated with dementia-risk in the human brain, rather than a single direction of activation, and consistent with microglia existing as a continuum of activation states rather than discrete states^73^. DAB2 knockdown also depleted transcriptional networks associated with lipid-droplet associated microglial states typically linked to ageing and APOE4 carriers^24,47^ and impaired cholesterol biosynthesis and lipid-handling pathways. Although this seems to contradict a protective role for DAB2, lipid-handling is double-edged: maladaptive lipid-droplet overload produces dysfunctional, pro-inflammatory microglia and secrete APOE-dependent factors that promote Tau-phosphorylation and neurotoxicity^24^, yet the capacity to mobilise lipids is necessary for microglia to coordinate neuronal activity and mount reactive, potentially protective AD states^48^. Consistent with this, DAB2 signalling axes correlated with LDAM signatures, and DAB2 loss on an APOE4 genetic background showed a blunted but directionally similar response.

We found that acute loss of DAB2 induced cytokine-responsive (CRM) states in iPSC-microglia under otherwise unstressed conditions, whereas CRM states were depleted and DAM states enriched with advancing AD-pathology in the human brain. These apparent differences likely reflect context-dependent effects of DAB2 given that the iPSC-microglial DAB2 knockdown experiments were acute and were carried out in the absence of AD-stressors and their surrounding multi-cellular environment. The direction-consistent finding across both systems is that DAB2 supports the activated DAM/TREM2 program: removing DAB2 depletes this program in-vitro, and in the human brain the microglia that reach DAM states retain DAB2 (accounting for the positive DAB2-DAM correlation), while bulk DAB2 falls with APOE4/E4 genotype. Set against the xenograft microglial trajectory described above, in which homeostatic microglia transitioning to protective disease-associated-(DAM) or amyloid-responsive microglia (ARM) or following a parallel pro-inflammatory pathway along the cytokine-responsive CRM/CRM-1/CRM-2 axes, and predicting that DAM-like microglia can additionally transition towards CRM states^22,26^; we interpret the in-vitro phenotype as a relief of DAB2-mediated inhibition of pro-inflammatory pathways. In-vitro, acute DAB2 loss stalls the DAM transition and shifts microglia towards cytokine-responsive states, whereas in the chronic, amyloid-laden brain, the same axis may instead convert homeostatic and cytokine-responsive microglia into DAM/LDAM states, depleting CRM states as pathology advances. The in-vitro CRM phenotype may therefore reveal a cell-intrinsic function of DAB2 that is not fully recapitulated at the population level in the human disease trajectory. The sustained DAB2 expression in DAM-enriched microglia in advanced disease could also reflect microglia adopting maladaptive DAM states over time^62^. Whether DAB2 directly transduces APOE and RELN signalling or modulates these pathways by recycling their receptors remains to be determined.

Our gene network analyses of iPSC-microglia point to LRP1-LRP8-DAB2, rather than the canonical neuronal RELN-DAB1 pathway as the prominent axis dysregulated in microglia. This is directly relevant to AD as LRP1 is a principal microglial receptor for amyloid-beta clearance and APOE signalling^74^. We found that DAB2 was expressed ∼9-fold above DAB1 in adult human microglia with knockdown of DAB2, but not DAB1, resulting in robust transcriptional changes. The RELN/APOE receptor LRP8 however was a hub gene of the (blue) ribosomal/antigen-presenting module of which the APOE receptor LRP1 was also a member and corresponded to co-regulated gene pathways involved in ribosome biogenesis, translation and chromosome organisation. This gene module was further associated with homeostatic, antigen-presenting, resting and APOE4-lipid microglial states, all of which were suppressed by DAB2 knockdown. LRP1 receptor has dual endocytic and signal transduction roles in microglia essential for amyloid-beta clearance and for inhibiting pro-inflammatory NF-κβ, JNK and MAPK signalling pathways^74–76^ consistent with LRP1 restraining the adoption of pro-inflammatory CRM states. The LRP8 receptor has previously been shown to lock microglia in maladaptive homeostatic states in AD by promoting APOE4-LRP8-TGFβ upregulation of homeostatic check points which impair microglia from adopting protective neurodegenerative/activated DAM states^77^. Additionally, RELN signalling through LRP8-DAB2 modulates NF-κβ pro-inflammatory signalling in endothelial cells^16^, and chronic Reln-deficiency exacerbates microglial inflammation in amyloid mouse models^10^ nominating a possible convergence of APOE and RELN signalling through which microglial LRP8-DAB2^15^ could modulate microglial states and counteract pro-inflammatory pathways.

DAB2 has dual functions as an endocytic clathrin-adaptor protein and as a transducer of TGFß signalling^14,78–82^. Our re-analysis of human microglial transcriptomes showed that components of the APOE and RELN receptor-adaptor machinery changed in opposite directions with increasing amyloid pathology in AD: APOE and its receptor LRP2 rose with CERAD/amyloid scores, whereas VLDLR, LRP1 and DAB2 declined. One inferred interpretation is that rising APOE ligand becomes uncoupled from the endocytic machinery that normally internalises it in an amyloid-rich environment. Because DAB2 mediates clathrin-dependent endocytosis of LDL-receptor-family members including LRP2 and LRP1^80,83–86^, but the amyloid-responsive DAB2 isoform lacks several clathrin binding domains and shows reduced endocytic activity^55,85,87–89^, reduced DAB2 (or the preferential splicing of DAB2-202 in an amyloid environment^55^) could therefore either prolong receptor residence at the plasma membrane and sustain APOE signalling and/or impair uptake of APOE and amyloid-beta cargo; our data do not distinguish these. The same logic applies to the RELN receptors, integrins and Toll-like receptors (TLR2/4) whose surface availability DAB2 also regulates^90–94^. If DAB2 deficiency represents a gain of function in microglial RELN-LRP8 signalling, its attenuated response on an APOE4 background could be explained by the known role of APOE4 in sequestering VLDLR/LRP8 receptors in endosomes^6^, which reduces RELN receptor availability. Whether sequestration is sufficient to impair signalling in the absence of the exogenous addition of RELN remains to be tested; however, both our iPSC-microglia and our human brain analyses showed that APOE4/E4 microglia strongly upregulated APOE, consistent with a role for APOE4 impairing RELN receptor signalling. As our readouts are transcriptional, these changes may reflect feedback from altered receptor trafficking or the preferential splicing of the DAB2-202 isoform which shows attenuated endocytic function and is proposed to be translocated to the nucleus where this acts as a transcriptional co-activator^55,85,87,88^. LRP8 is also cleaved by gamma-secretase following RELN-binding^95,96^, and its intracellular domain can translocate to the nucleus where this regulates transcription including the downregulation of RELN^97,98^. Notably, DAB2 knockdown downregulated nicastrin, a gamma-secretase component, which would be predicted to limit LRP8 processing. Direct measurement of receptor surface levels and recycling in DAB2-deficient microglia, and exposure of APOE iPSC-microglia to exogenous RELN, in additional isogenic lines, will be needed to resolve these mechanisms.

Microglia play a multiple roles in their response to amyloid pathology, by phagocytosing Aß aggregates, through the removal of damaged amyloid-laden synapses to mitigate amyloid build-up^23^, and by transitioning from homeostatic states to amyloid-associated states necessary for their protective functions^26,45,46,49^. DAB2 has previously been shown to endocytose lipoprotein receptors^81^ pointing to a potential interaction with LRP1 which binds and mediates microglial amyloid-beta clearance^22,99^. Our transcriptomic analyses show that DAB2-deficient microglia downregulate the nicastrin enzyme and endolysosomal proteins important for amyloid-beta degradation, suggesting that microglial DAB2 signalling contributes to the clearance and degradation of amyloid-beta. Microglial phagocytosis also sculpts synapses through their initial protective removal of amyloid-laden synapses^23^, but which over time becomes maladaptive with overactivated microglia leading to excessive synaptic clearance which is thought to contribute to cognitive decline in AD^20^. Our data show that DAB2 knockdown downregulated lysosomal transport/phagocytic pathways, together with integrin–extracellular matrix and complement-activation pathways, whereas CD68 is upregulated in microglia of Reln-deficient mice *in vivo*, implicating that RELN– and DAB2-signalling in endolysosomal dysfunction, which alongside altered complement signalling, could compromise synaptic pruning and phagocytosis. We predict that RELN–APOE signalling via DAB2 could alter endolysosomal/phagocytic function and contribute to synaptic deficits in AD via impaired microglial motility and their impaired C3-tagging of synapses and defective interactions with neurons^2,24,28,29^. Consistent with this, DAB2 limits integrin availability at the plasma membrane, predicted to reduce extracellular matrix cell adhesion^100^,and to affect the cytoskeletal and actin dynamics underlying microglial surveillance and cell motility. While we cannot exclude that the microglial defects in Reelin-deficient mice are secondary to ectopic neuronal positioning and signalling^1,60^, the altered morphologies and CD68 upregulation we observed at postnatal stages (∼P22) are consistent with endolysosomal changes and increased microglial activation arising in a chronic Reln-deficient environment. While DAB2 deficient iPSC-microglia and Reln-deficient murine microglia suggest opposing effects on endolysosomal function, this could be reconciled by our observations that genes important for lysosomal acidification and lysosomal degradation were downregulated in DAB2-deficient microglia, suggesting an impairment in the turnover of endolysosomes which could potentially result in the larger CD68 sum volumes observed in microglia within the Reln-deficient mice. Our network analysis places the dementia-risk gene ABI3, a WAVE regulatory complex subunit, in the homeostatic (salmon) module, and a second WAVE subunit CYFIP1, in the DAM/LDAM (purple) gene module alongside DAB2 and TREM2, where it was the most strongly suppressed by DAB2 knockdown (r= –0.82, padj =6.6 x 10^−4^). Arp2/3 activity maintains microglial morphological ramification and surveillance, and conditional depletion of Cyfip1 in mouse microglia reproduces these deficits while increasing CD68+ lysosomal volume^57^, paralleling the enlarged CD68+ compartment we observed in Reelin-deficient mice. Together, these observations suggest that RELN-DAB2 deficiency could impair both the uptake/phagocytosis of synaptic material and its subsequent lysosomal degradation, potentially resulting in enlarged lysosomes, although future studies are required to validate this. The enrichment of the dementia-risk locus CR1 in the salmon module is consistent with DAB2 deficiency altering microglial gene networks that govern complement-mediated phagocytosis of synapses, which may contribute to impaired synaptic function in AD, alongside microglial degradation of amyloid-beta through the dementia risk locus ACE (Fig. 8, Supplementary Table 7)^20^. Future studies exposing APOE iPSC-microglia to exogenous RELN or co-culturing microglia with RELN-expressing neurons in the presence of amyloid are needed to assess whether RELN signalling directly shapes microglial endolysosomal, phagocytic function and their interactions with neurons to compound synaptic deficits in an AD-like environment.

The two siRNAs used in DAB2 knockdown experiments targeted different DAB2 isoforms: siRNA-96 targeted both DAB2-201 (murine p96) and the shorter DAB2-202 (murine p67) isoform induced by amyloid^55^ which is the predominant isoform expressed during embryonic development^85^ that lacks the central clathrin– and AP-2 binding motifs and is therefore less efficient in mediating endocytosis, and is further upregulated in microglia under stress^71^. The siRNA-98 which targets the DAB2-201 alone suggested that this isoform inhibits DNA damage-induced senescent pathways. While DAB2 has not been directly linked to microglial senescence, DAB2 is upregulated in senescent pro-inflammatory lung cells^101^, and is a core TGFß/Smad adaptor through which TGFß promotes senescence^79,102^. Here, DAB2-loss induced the TNFα-NFκ-β pathway, which is a canonical driver of the senescence associated secretory phenotype^103,104^. While future studies specifically targeting the DAB2-202(murine p67 isoform in mice) are required, our data raise the possibility for differential roles for the endocytic DAB2-201 (murine p96 isoform in mice) and amyloid-induced DAB2-202 isoforms in modulating microglial states, lipid handling, endolysosomal/phagocytic, extracellular matrix, senescent, amyloid beta and inflammatory pathways. This also raises the possibility that the amyloid-induced DAB2-202 isoform represents an aberrant reactivation of the developmental isoform^55^, as has been observed for the complement pathway^20^. Nonetheless we cannot exclude the deeper knockdown achieved by targeting both isoforms could mediate these effects. Whether the two DAB2 isoforms are differentially regulated by amyloid in the human brain as has been shown in mice^55^, should be addressed in future studies.

APOE4 lowers the age of onset and increases the risk of dementia, partly through heightened amyloid deposition^11^, and increased microglial activation and neuroinflammation^24,69^; which we propose is relevant to microglial DAB2-mediated signalling. A prior genome-wide analysis of APOE4/E4 homozygotes identified RELN-DAB1 as a risk pathway genetically interacting with APOE4^8^. Our study nominates DAB2 as an additional node at which this interaction may occur, and points to a pathway-level interaction with APOE genotype. Although the transcriptional response to DAB2-knockdown was predominantly concordant across APOE genotypes (r=0.86), knockdown of DAB2 in an APOE4 background produced an attenuated depletion of homeostatic and antigen-presenting (HLA) states and a blunted shift toward pro-inflammatory cytokine-responsive (CRM) microglia relative to APOE3, whereas effects on DAM and interferon-responsive states did not differ by genotype. This was mirrored by an attenuated induction of TNFα-NFκ-β, inflammatory-response and senescence leading-edge genes in APOE4/E4 microglia. This is consistent with APOE4 microglia already occupying a more activated, homeostatic-depleted baseline, leaving less residual homeostatic identity for DAB2 to erode. In addition, we found that TREM2 protein was reduced by DAB2 knockdown in APOE3/E3 microglia in response to myelin, a known inducer of DAM-states in microglia^52^, but not in APOE4/E4 microglia in which TREM2 changes were more variable. TREM2 has been shown to underlie the transitioning of homeostatic to disease-activated and antigen presenting (HLA) states, with some evidence this may also be required to promote homeostatic towards some CRM states^26^. The increased variability of microglial TREM2 in an APOE4 background could potentially reflect an asynchronous progression towards distinct TREM2-dependent DAM/HLA or parallel CRM states in APOE4/E4 microglia^26^. Whether APOE4/E4 impairs the adoption of microglial protective DAM states early in disease, while leaving open the possibility that chronic pathology later locks microglia into maladaptive DAM/TREM2 states, remain to be tested. A potential explanation converges on DAB2 modulating APOE receptor availability and possibly on the reported nuclear localization and transcriptional activity of APOE^105^. APOE4/E4 microglia expressed significantly higher levels of APOE, and APOE4 is predicted to reduce APOE– and RELN-receptor availability, so a further perturbation through DAB2 knockdown could have less to act on.

An attractive, untested implication of this work is that the DAB2 axis could couple neuronal vulnerability to microglial states. Because RELN and APOE compete for the shared lipoprotein receptors (LRP8/VLDLR), a loss of neuronal RELN signalling in early AD from the first-affected RELN-expressing entorhinal cortical neurons^106,107^.would shift the receptor-adaptor balance toward unopposed APOE signalling. We predict that this destabilises microglial homeostasis and impairs microglial adoption of DAM/TREM2 states which impairs microglial clearance of amyloid, lipid-homeostasis and their phagocytosis of damaged synapses. Combined with APOE4-driven APOE upregulation and DAB2 decline we observed with advancing amyloid pathology, this could impair the DAB2-promoted transition into TREM2/DAM states precisely in early vulnerable neural circuits where synaptic deficits and Tau pathology begins^10,107,108^. *In vivo* the chronic combination of RELN loss with APOE4-driven APOE upregulation, would be predicted to exacerbate gliosis, amyloid deposition and Tau pathology in these vulnerable circuits^24,25,47,69^. Testing this will require co-culturing microglia with RELN-expressing neurons under amyloid challenge and spatially resolving DAB2-expressing microglial states relative to RELN-expressing neurons in the human entorhinal cortex.

A major limitation of this work is its primary focus on transcriptional readouts following DAB2 knockdown in a single isogenic iPSC-line, which may not be recapitulated at the functional or protein level beyond the TREM2 reduction we observed, and iPSC-line specific effects cannot be excluded. While our work reveals cell-autonomous roles of DAB2, knockdown was acute and performed without amyloid, RELN or inflammatory stressors. Our pathway enrichment approach also used RELN-core gene sets that also captured the APOE pathway, because the canonical RELN receptors bind both ligands, and because DAB2 also interacts with both canonical RELN receptor (LRP8)^13,14^ and APOE LDL-receptors^80,81,83,84,86^. Leading-edge analysis indicated that the ‘RELN’-signal in the human brain is driven primarily by APOE and amyloid networks, with DAB1 expressed at low levels. DAB1 knockdown achieved only trend-level reduction, consistent with low basal expression, but produced pathway-level changes concordant with DAB2. DAB1 could potentially contribute to microglial function under RELN stimulation that our basal cultures did not provide, and we found that APOE4/E4 microglia showed reduced DAB1 and pY232-DAB1 that was amplified under LPS, consistent with attenuated signalling under inflammatory stress. The human associations were pseudobulk and included few APOE4 carriers, and the Reln-deficient mouse is a germline model with severe lamination defects and did not isolate microglial RELN signalling. Future work combining exogenous RELN and amyloid-beta, co-culture with RELN-expressing neurons, subtype-specific microglial state markers, and multiplexed or spatial transcriptomic profiling of DAB1/DAB2 expressing microglia in the human brain will be needed to resolve these questions.

## Conclusions

We identify DAB2 as a cell-autonomous regulator of microglial DAM/TREM2 states associated with dementia risk. We find that RELN and APOE receptor expression are dysregulated in microglia in the human AD-afflicted brain in response to increasing AD-pathology, and that DAB2 loss in APOE iPSC-microglia depletes gene networks regulating DAM/TREM2-like transcriptomic states associated with endolysosomal and phagocytic function, lipid-handling and amyloid-beta degradation and clearance pathways. A loss in DAB2 signalling under basal conditions was sufficient to shift microglia towards pro-inflammatory cytokine-responsive states, that were associated with enhanced TNFα-NFκ-β inflammatory pathways, with the inflammatory arm attenuated on an APOE4/E4 background. Together we propose that APOE-DAB2 is a novel signalling axis and potential point of intervention for modulating the protective to maladaptive DAM transition, linking APOE genotype to the microglial state transitions that shape early AD.

## Materials and Methods

Ethical Compliance: All procedures performed in studies involving mice were performed in accordance with the UK Animals (Scientific Procedures) Act 1986 and the UCL institutional guidelines under Prof. Parnavelas PPL 70/8227. All human iPSC-microglia experiments were carried out in accordance with UCL UK DRI guidelines.

### Floxed exon-3 Reln conditional mice and transgenic mouse lines

All experimental procedures were performed in accordance with the UK Animals (Scientific Procedures) Act 1986 and the UCL institutional guidelines under Prof. Parnavelas PPL 70/8227. The conditional Reln mouse strain used for this research project was created from C57BL/6 mouse ES cell clones (DEPD00533_5_A03 and DEPD00533_5_F01), generated by the Mouse Biology Program (www.mousebiology.org), and made into Relntm1a(KOMP)Mbp ES cells using the KOMP Repository (www.komp.org) at the University of California Davis (U42RR024244)^109^. A promoter driven cassette was used to generate the ‘Knockout-first allele’ that targeted a critical region in exon 3 (22937-22970) of Reln Chr5:21884454-22344702 bp, common to all transcript variants that, when deleted, creates a frame-shift mutation. Conditional Reln^fl/+^ mice were obtained by their mating with Rosa26FLPe knock-in mice (B6.129S4-Gt(ROSA)26Sortm1(FLP1)Dym/RainJ) (Stock)^110^ to remove the short flippase recognition (Frt) sites; and their subsequent mating with ActinCre mice (B6N.FVB-Tmem163Tg(ACTB-cre)2Mrt/CjDswJ)^111^ for the ubiquitous depletion of Reelin. Cx3cr1-Gfp (B6.129P-Cx3cr1tm1Litt/J) transgenic mice obtained from Jackson Laboratories (Stock 005582) enabled the genetic tracing of microglia, dendritic cells, natural killer cells, and monocytes, but not differentiating macrophages, where this was downregulated. Tie2-Gfp mice from Jackson Laboratories (Stock Tg(TIE2GFP)287Sato/J, Stock Number 003658) were used for genetic tracing of the developing cerebrovasculature, in addition to a population of microglia^112,113^. Both the male and female postnatal and adult forebrains were used in this study. The day on which the vaginal plug was found was considered embryonic day (E) 0.5, and animals of both sexes were used in our experiments.

### iPSC-microglial differentiation and culture

iPSC-human microglia were differentiated from APOE3/E3 and APOE4/E4 isogenic BIONIO37 iPSCs generously gifted by JP and JH from EBiSC, through the generation of embryoid bodies and mesodermal induction prior to their differentiation along the myeloid lineage, according to Huang et al., (2024)^69^ and Washer et al. (2022)^114^ with some modifications. Briefly, APOE BIONIO37 iPSCs were expanded and maintained on vitronectin-coated substrate (Life Technologies) in supplemented StemFlex medium (Gibco) or E8 media (Gibco) respectively, and according to the manufacturer’s instructions until 80% confluency was reached. iPSCs were subsequently dissociated with Versene (Gibco) and Accutase (Gibco) and embryoid bodies were generated by seeding 15,000-20,000 iPSCs/well in ultra-low attachment U-shaped 96 well plates (Corning) in StemFlex media (Gibco) supplemented with mesoderm-inducing 20ng/mL SCF, 50ng/mL BMP4, 50ng/mL VEGF, 20ng/mL SCF, (all from Peprotech EC Ltd), and either 10µM RockInhibitor (Tocris) or 1:100 Revitacell (Gibco). Following seeding, embryoid bodies were replenished with previously described supplemented StemFlex media without Rockinhibitor on Day3 and two days later, they were transferred to gel-trex-coated (Gibco) 75 cm^2^ culture flasks in myeloid differentiation media, consisting of X-Vivo 15 media (Lonza) supplemented with 50µM of antioxidant beta-mercaptoethanol (Gibco), penicillin/streptomycin, 100ng/mL of MCSF and 25ng/mL of IL-3 (Peprotech EC Ltd) with 1x Glutamax (Gibco). The media was then replaced one week later and once a week with the first three passages of iPSC precursors discarded. In the fourth week, iPSC-microglia precursors were harvested and plated in DMEM/F12 Hepes-containing and phenol-red free media (Gibco) supplemented with 2% insulin transferrin selenium (Gibco), N2 supplement (Gibco), 200µM of Monothioglycerol (MTG), 1x Glutamax (Gibco), non-essential amino acids (Gibco), 5µg/mL human insulin (Merck), 100 ng/mL IL-34, 25ng/mL MCSF, and 25 ng/mL TGFβ (all from Peprotech EC Ltd.). iPSC-microglia were collected or fixed on day 14 for RNA or protein analysis. Immunostainings of iPSC-microglia were carried out using the anti-DAB1 B3 antibody (1:1000)^3^ and PY232 and PY198 DAB1 phosphorylated antibodies (1:350) (My Biosource). A minimum of 3 independent microglial differentiations were carried out for each APOE genotype in this study. All siRNAs were purchased from Ambion Life Technologies. For DAB1 knockdown experiments the following siRNAs were used: DAB1s3893 referred to as DAB1-893 (Cat 4457298/ASO2LDM9), DAB1s3894 (Cat 4457298/ASO2LDM8) referred to as DAB1-894. For DAB2 experiments DAB2s3896 referred to as DAB2-96 (4457298/ASO2P4CT) and DAB2s3898 referred to as DAB2-98 (4457298/ASO2P4CT) were used. siRNA was used to acutely knockdown DAB1 or DAB2 over a 48hour period, using Lipofectamine transfection (LifeTechnologies) for 5 minutes at room temperature in accordance with the manufacturer’s protocol.

### Bulk RNA sequencing analysis of ioMicroglia-deterministically programmed human iPSC-derived microglia

ioMicroglia are human microglia deterministically programmed from human iPSCs using optiox (optimised inducible overexpression) technology^115^ (bit.bio Ltd, UK). iomicroglia were cultured to day 10 following manufacturer’s protocol. Bulk RNA sequencing was performed on wildtype ioMicroglia Male (cat.no. io1021S, bitbio Ltd, Cambridge, UK) from an APOE3/E3 background, and two clones from genetically matched ioMicroglia with with the APOE4/E4 C112R/C112R mutation (CL69, cat. No. io1032 and CL83, cat. no, io1031. bitbio Ltd). All reprogramming was carried out by bit.bio. Total RNA was extracted using the All-prep RNA/DNA extraction kit (Qiagen), following the manufacturer’s instructions. RNA concentration was determined using the Qubit Broad Range RNA assay (Thermo Fisher Scientific) and total RNA integrity (RIN) assessed using the RNA ScreenTape assay (Agilent) ensuring RIN values greater than 8. An input of 150 ng of total RNA was used for each sample for the RNA-seq library preparation using the CORALL mRNA-seq V2 kit (Lexogen), prepared according to the manufacturer’s instruction for short inserts.

### High-content automated imaging and protein quantification

High-content automated single-cell imaging and protein quantification of DAB1, pY232-DAB1, pY198-DAB1 and DAB2 was performed on APOE3/E3 and APOE4/E4 iPSC-differentiated microglia cultured in 96-well plates under basal (control siRNA), LPS (100 ng/ml, 24 h) and IFNγ (20 ng/ml, 24 h), myelin (3mg/mL) stimulation conditions. Cells were fixed with 4% paraformaldehyde, permeabilised with 0.1% Triton X-100 and immunostained with validated primary antibodies. Images were acquired on an Opera Phenix Plus high-content confocal system (Revvity) using a 40x water-immersion objective. Automated image segmentation and single-cell feature extraction were performed in Harmony 4.9 software (Revvity). Per-cell intensity, morphology and texture features were extracted using the cell-paint masking algorithm resulting in 584 analysed feature readouts. A per well median across cells in that well was calculated, with population-level intensities then normalised to the mean of control-siRNA APOE3/E3 wells within each experiment plate. Each well value was standardised against the individual plate and Ctrl baseline using a robust z-score where median value + median absolute deviation was used. The robust Z-score was used in all downstream analyses with pooling across plates valid because each plate’s z was normalised to its own internal reference control. Linear mixed models (LMM, using lmerTest::lmer) were fitted in R using the lme4 package, with fixed effects for APOE genotype, stimulation condition and their interaction, and random effects for well and biological replicate.

### Bulk RNA-seq of APOE iPSC-microglia

Paired-end bulk RNA-sequencing reads from 39 samples were processed with the community-curated nf-core/rnaseq pipeline (v3.23.0) implemented in Nextflow (v25.10.4)^116–118^. In brief, raw FASTQ files were validated and quality-controlled (FastQC 0.12.1), adapter-and quality-trimmed (with Trim Galore! (0.6.10), a wrapper around Cutadapt^119^ and FastQC, using default settings), and aligned to the human reference genome with STAR (2.7.11b)^120^. All reads were aligned and quantified against the *Homo sapiens* GRCh38 primary assembly with Ensembl release 111 gene annotation. Resulting BAMs were coordinate-sorted and indexed with SAMtools (1.21 within STAR_ALIGN; 1.22.1 for post-processing)^121^. Then, gene-level count matrices were generated from STAR-aligned BAMs using featureCounts (v2.0.3). STAR overall mapping rates were consistently ∼99% (uniquely mapped ∼93%) and ribosomal RNA content was negligible (∼0%).

Differential gene expression analysis was performed using DESeq2^122^ implemented within the BIONmicroglia2026-01-26 analysis pipeline. Raw count data were normalised using median-ratio size factors. Genes were retained if these corresponded to more than 10 counts in at least 3 samples prior to analysis. Dispersion estimation and Wald tests were used to test differential expression. Samples corresponded to independent differentiations: APOE3/E3 CtrlsiRNA n=4, APOE4/E4 CtrlsiRNA n=4, APOE3/E3 DAB1-893siRNA n=5, APOE4/E4 DAB1-893siRNA n=4, APOE3/E3 DAB1-894siRNA n=5, APOE4/E4 DAB1-894siRNA n=3, APOE3/E3 DAB2-96siRNA n=3, APOE4/E4 DAB2-96siRNA n=3, APOE3/E3 DAB2-98siRNA n=5, APOE4/E4 DAB2-98siRNA n=3). Genotype effects in control samples were modelled using the design formula ∼Genotype (Q1; reference level: APOE E3/E3); siRNA/knockdown effects were modelled with ∼siRNA or ∼dab1_status (Q2); combined genotype-knockdown effects with ∼Genotype + siRNA or ∼Genotype + kd_status + Genotype:kd_status (Q3/Q4); and interaction effects were assessed using a likelihood ratio test (Q5). Log2 foldchange shrinkage was applied using apeglm for named coefficients or ashr for contrast-based comparisons. Differentially expressed genes (DEGs) were defined as padj < 0.05 (Benjamini-Hochberg correction) and |log2 foldchange| >= 0.58 (equivalent to ∼1.5-fold change). Volcano plots display lfcShrink-corrected estimates with significance thresholds indicated.

### fGSEA Gene Set Enrichment Analysis

Differential expression analysis was performed with DESeq2^122^, and a per-gene Wald statistic was extracted for each contrast to generate ranked gene lists. Duplicated genes were removed by selecting the gene with the maximum |Wald statistic| per HGNC symbol and ranked in descending order of the signed Wald statistic to produce a pre-ranked input list per contrast. Gene-set enrichment of these ranked lists was performed using the multilevel fgsea algorithm^123^, with infinite simple gene-set permutations, seed = 42. Only genes with ≥10 counts in ≥3 samples (set before DESeq2) were used, with a minimum of 10 genes and maximum of 500 genes per set. Four classes of gene sets were interrogated: (1) Mancuso et al. (2024)^26^ microglia subtype marker genes, derived from the top 100 genes per cluster (avg_log2FC > 0.1, pct.1 – pct.2 >= 0), ranked by log2FC x-log10(padj); (2) literature-curated microglial state programmes including homeostatic (Mathys et al. 2019), disease-associated microglia (DAM; Keren-Shaul et al. 2017), amyloid-responsive (ARM) and interferon-responsive (IRM) microglia (Sala-Frigerio et al. 2019), and lipid-droplet-associated microglia (LDAM; Marschallinger et al., 2020^47^); (3) RELN/DAB1/DAB2-seeded gene lists derived from STRING v12 interaction partners (required_score >= 400), MSigDB CP:REACTOME pathways, and MSigDB extended name-matched gene sets; and human microglia AD brain datasets (Alzheimer’s Disease Progression Atlas^29^, microglia isolated from APOE3/E3 healthy or APOE3/E4, APOE4/E4 AD-afflicted brains (GSE254205)^24^. Normalised Enrichment Scores (NES) and BH-adjusted p-values were calculated within each contrast. NES heatmaps display tile fill proportional to NES (diverging blue-white-red scale, range +/− 2-3.5); tiles with padj < 0.05 indicated by stars, with tile labels showing NES (top) and padj (bottom). For microglial-focused pathway enrichment analyses, a hypothesis-driven curated 58 pathways were generated apriori from MSigDB Hallmark, KEGG, Reactome and GP:BP databases based on six prior literature microglial-biology categories (lipid/cholesterol, phagocytic/lysosomal, AD/amyloid, cytokine/TLR-IFN/microglial, ECM/integrin, mitochondrial/proteostasis) defined by Keren-Shaul et al., 2017^46^, de Mathys et al., 2024^106^, Sala-Frigerio et al., 2019^45^; Wachter et al., 2024^29^). Redundant pathways within each category were collapsed based on Jaccard ≧ 0.4 between leading-edge gene sets and heatmaps displaying the top 2 pathways per category based on composite rank (within-category min BH-q + within-category mean |NES| across all contrasts). To rule out this curated focus excluded biologically significant categories, the full MSigDB Hallmark+ KEGG+ Reactome collection (2,075 sets) was tested as an unbiased control and are shown in **Supplementary Figures**, with top hits from unbiased analyses confirmed to belong to the same six categories.

### PCA of NES matrix scores from fgsea Gene Enrichment Studies

Principal component analysis was carried out on the matrix of normalised enrichment scores obtained through fgsea analyses of ranked gene lists for all 28 contrasted datasets (rows) from six experimental groups. This resulted in a NES matrix of 28-dimensional space (one dimension per gene set) with the following dimensions: rows: 28 differential expression contrasted ranked gene sets (rows) from the following experimental groups: BIONIO37A-/4 microglia, BIONIO37A-/4 + siRNA DAB1/DAB2 KD microglia, human brain AD APOE microglia snRNA seq (Haney et al., 2024^24^, GSE254205), iPSC-MG APOE4 microglia (Haney et al., 2024^24^, GSE254205), developmental microglial trajectories (Human: Galatro et al., 2017^28^ Aged vs Young; Kracht et al., 2020^27^ GW early vs late; GSE99074; Mouse Ostrem et al., 2024^124^ P14 vs E13, P14 vs E15), and microglia at advanced stages of AD from the AD progression Atlas (Wachter et al., 2024^29^). Columns (features) included 10 gene sets: 2 RELN-axis co-expression signatures (RELNcore-DAB1, RELNcore-DAB2) and 8 MG activation state gene sets (HM, DAM, HLA, CRM, IRM; LDAM markers (Haney et al., 2024^24^) & Victor et al., 2022^48^ APOE4_lipid_iMG). To avoid multiplication of near identical MG state signatures we used only one signature per microglial subtype to avoid multiplying the weight of a given microglial state – so redundant variants were collapsed into one consensus signature per state ie we chose the Mancuso et al., 2024^26^ states with two representative LDAM signatures from Marschallinger et al., 2020^47^ and Victor et al., 2022^48^, so as not to double reference the Haney et al., 2024^24^ gene set. The direction of all contrasts was specified as Disease UP vs Healthy/Control, Age UP vs Young, APOE4 UP vs APOE3, lipid-laden microglia LD+ UP vs LD-ve, across all datasets during fgsea analyses or prior to PCA. PCA code was generated with Claude and performed in R through Cursor using prcomp() with centring and unit-variance scaling (center =TRUE, scale =TRUE). Scaling ensures al 10 gene sets contribute equally to the matrix regardless of their absolute NES range. The SVD based algorithm was used (default for prcomp). No rotation was applied. Variance explained with developmental/ageing/CRM and homeostatic/DAM/LDAM microglial trajectories: PC1 = 46.7%; and PC2 = 19.7% resolving RELN-DAB signalling axes from CRM/IRM MG states; and PC3 = 13.9% resolving HLA-DAM from HM-CRM states. The first two PCs together capture 66.4% of total variance in the NES matrix and mapping onto 2D representation of the microglial contrast landscape during disease. The dashed blue curve labelled DAB1/DAB2 KD trajectory is an illustrative arc not a data-fitted curve.

### Co-expression network analysis of microglial transcriptomes from the Progression AD atlas (Wachter et al., (2024)^29^

Per-donor microglial pseudobulk profiles were reconstructed from Wachter et al., (2024)^29^ AD Progression Atlas single-nucleus data for the four profiled brain regions (entorhinal cortex (EC), inferior temporal gyrus (ITG), prefrontal cortex (PFC) and primary visual cortex (V1). For each region, raw microglial single-cell counts were summed per donor to form a genes x donor matrix, normalised to counts per million and log2 (CMP+1) transformed, resulting in 113 region x donor pseudobulk profiles (EC, n=23, ITG, n=31, PFC, n=28, V1 n=31). Each donor was annotated with numeric Braak stage (0-II, II-III, V, VI), neuritic-plaque/CERAD score (none =0, sparse =1, moderate =2, frequent =3) and composite pathology stage (Braak + CERAD + diagnosis). Co-expression among the RELN pathway leading-edge genes (together with DAB1 and DAB2) was computed in a region-specific manner and for each gene pair, a Pearson correlation was computed separately within each region, Fisher z-transformed, averaged across regions weighted by each region’s degree of freedom and then back-transformed to reflect genuine donor-to-donor co-expression for each region. The combined two-sided significance was derived from the weighted mean z score and Benjamini-Hochberg (BH) false discovery rate (FDR) corrected across all gene-pairs. The naively pooled correlation (n=113 regions) was computed in parallel to identify and discard region-driven artefactual edges. In the resulting network (Fig. 1d), nodes are colour-coded by each gene’s association with amyloid burden (mean DESeq2 Wald statistic for CERAD across the four regions; node size is proportional to Wald ranked statistic. Grey edges denote positive co-expression (r≥+0.5) and dashed edges indicate significant negative correlations (r≤-0.3, FDR <0.05).

### Correlating DAB1 and DAB2 microglial expression levels from the Progression AD atlas (Wachter et al., (2024) with defined microglial activation signatures

Using the same per-donor and within each region microglial pseudobulk profiles (log2 CPM +1), expression was standardised (Z-scored) within each region to remove region-level offsets. A disease-associated-microglial (DAM)/activation signature score was computed per donor as the mean z-score of defined DAM markers^26^, and the same mean-z scoring was applied to each additional curated microglial-state signature. Standardised DAB1 and DAB2 expression was correlated with the signature scores across all 113 region x donor profiles by Pearson correlation (donors coloured by CERAD score; Fig. 1e).

### Overlap enrichment of DAB2 siRNA-knockdown WGCNA modules against microglial state signatures defined from mouse^46,47,62^ and human^29,61–64^ AD transcriptomic studies

Overlap between DAB2 siRNA knockdown WGCNA co-expression modules (purple, DAB2/TREM2 (n=150genes); blue, LRP8/LRP1 (n=1001 genes); brown, ITGB8 (n=927 genes); salmon, MRC1/P2RY12 (n=101 genes); each summarised by its module eigengene) and microglial-state marker-gene signatures from mouse and human AD transcriptomic studies was tested with a two-sided Fisher’s exact test. For each module signature pair, a 2×2 table of shared versus non-shared genes was constructed against the set of genes represented in the WGCNA network as the background universe, and the overlap odds ratio recorded. Heatmap cells are colour coded from low (white) to high (red) odd’s ratio scale. Benjamini-Hochberg FDR *p<0.05, **p<0.01, ***p<0.001).

### WGCNA

Pre-processed DeSeq2 expression matrices (genes retained at ≥10 counts in ≥3 samples) from n=39 BIONIO37A/A-4 isogenic APOE iPSC-derived microglia (APOE4/E4 vs APOE3/E3 +/− DAB1siRNA893+894 or +/− DAB2siRNA96+98 KD samples) were variance-stabilised-transformed (VST) and batch-corrected (23,320 genes x 39 samples) (Supplementary Fig. 9a). The top 10,000 most variable genes were retained and QC-filtered (noisy genes with zero or near-zero variance were removed, genes with missing NA values, and samples with < 4 detected genes using the WGCNA goodSamplesGenes function). A signed-hybrid co-expression network was constructed with WGCNA and the soft-thresholding power of 12 selected targeting scale-free topology fit of R^2^≥0.85 (R^2^ =0.846, mean connectivity =6.4). Modules were detected with blockwiseModules function (network type= signed hybrid, TOMType= signed, Pearson correlation, minimumModuleSize of 30 genes, mergedCutHeight= 0.25, deepSplit=2, maxBlockSize=10,000, seed=42), with module eigengenes defined as the first principal component and module membership as signed kME. This yielded 14 modules (36.8% unassigned grey); the 7 modules with ≥1 BH-significant module-trait correlation are reported (number of symbol-annotated genes/ total number of genes) (brown 586/927, blue 854/1001, green 423/589, magenta 214/232, greenyellow 126/130, salmon 100/101 and purple 147/150 genes). Module eigengenes were correlated with sample traits (APOE genotype and DAB1/DAB2 knock-down) by Pearson correlation with BH FDR correction. Each module was assigned a microglial state by over-representation analysis (one-sided Fisher exact test, BH-FDR across the module x signature matrix, restricted to the WGCNA gene universe) against curated reference signatures (homeostatic/DAM/CRM/IRM/HLA/ribosomal states, Mancuso et al., (2024)^26^; lipid/LDAM signatures from Marschallinger et al., (2020)^47^, Nugent et al., (2020)^44^, and an LXR cholesterol-efflux set Victor et al., (2022)^48^. DAB2 and TREM2 co-localised to the purple module (DAM/LDAM; lipid metabolism) and ITGB8 was a hub gene of the brown CRM/pro-inflammatory module, with LRP8 and LRP1 part of the ribosomal/HLA module. Over-representation of dementia-risk genes within each WGCNA module was assessed by a one-sided hypergeometric test (equivalent to a one-tailed Fisher’s exact test). The reference list was built from the 75 dementia-risk loci reported by Bellenguez et al., 2022^66^ i.e. the 35 lead genes at the 33 known loci (Supplementary Table 5 from Bellenguez et al., 2022) and the 55 Tier 1 and Tier 2 prioritised genes at the 42 new loci (Supplementary Table 20 from Bellenguez et al., 2022)^125^, plus APOE, which lies outside the 75 loci but is the established dominant risk signal totalling 91 genes; 37 of the 91 were present in the 10,000-gene WGCNA network. Each of the 15 modules (including the unassigned grey module) was tested separately against a background of all 10,000 gene network genes, and p values were corrected across modules by Benjamini-Hochberg procedure.

### Use of AI (Claude AI, Cursor)

Anthropic Claude and Cursor was used to assist in writing and debugging analysis and plotting coding (for differential DESeq2 analysis (Love et al., 2014)^122^, gene set enrichment (fgsea) (Korotkevich et al., 2021^123^), principal component analyses (PCA) of both DESeq2 and protein cell-paint analyses, WGCNA, co-expression network analysis of AD Progression atlas microglial transcriptomes, overlap enrichment studies with defined microglial state signatures, and merging of experiments from high-content “FullPlate” image-analysis pipelines), to produce data figures from the authors’ own experimental and sequencing data, or from publicly available human brain transcriptomic datasets, and to assist in editing and condensing the manuscript text. No AI tools were used to generate or alter primary data or to create non-data images. All AI-assisted code, analyses, figures and text were reviewed, validated and edited by the authors, who take full responsibility for the accuracy and integrity of the work.

## Acknowledgements

An extra special thanks is given to U. Borello (U of Pisa) for critical reading of the manuscript, to B. J. Steventon (Cambridge University) for access to Imaris software, to A Barrios (CDB, UCL) for help with confocal funding, to A. Greig, T. Robson, M. Redd, V. Silio, and C. Thrasivolous (CDB, UCL) and N. Cade (DRI, UCL), for advice with image acquisition and Imaris analysis, and to A Eddaoudi for help and advice with FACS experiments, to A. Goffinet for generously gifting the anti-G10 RELN and C17 C-RELN antibodies. This work was funded by grants from the Biotechnology and Biological Sciences Research Council [J.G.P.; BB/N002105/1), the Alzheimer’s Society (M.B.; 614 (AS-PDF-22-030) and (D.S., JH UK DRI-1210).

## Author Contributions

MB, WDA, JGP, and DAS contributed to the design and implementation of the research, MB, carried out most of the experiments. MB, WDA, DAS to the analysis of the results and to the writing of the manuscript, AK, HS, SL, LC, ZS designed DAB1 and DAB2 primers and validated DAB2 knockdown, BWH provided guidance and expertise on DAB1 experiments. DG, MB, AB carried out bulk RNA sequencing experiments and generation of fastq and cell counts and advised on R scripts and analysis. XH and TB generated CellPaintAnalysis R scripts. RN generously gifted Bitbio Microglia. NM advised on iPSC-microglial differentiation protocol. UY helped with coding. JP and JH generously gifted APOE iPSC-cell lines (BIONIO37A/A-4). RCMS & DA supported mouse work. All authors reviewed the manuscript. MB, JGP, DAS, JH conceived the original and supervised the project. This work was funded by grants from the Biotechnology and Biological Sciences Research Council (JGP.; BB/N002105/1), the Alzheimer’s Society (MB; 614/AS-PDF-22-030), and the UK Dementia Research Institute primarily through the Medical Research Council (DAS, JH UKDRI-1210).

## Permission to reuse and Copyright

De Mathys et al., (2023^126^; 2024^106^) allowed us to re-analyse their published transcriptomic studies.

## Figure Legends

**Supplementary Figure. 1.** **a**) Spearman-correlation table relating the expression of canonical RELN-axis genes (RELN, DAB1, LRP8, ITGB1) and APOE-axis genes (LRP1, APOE, DAB2) to AD-pathology severity in human microglia from the entorhinal cortex (EC), inferior temporal gyrus (ITG), visual area 1 (V1) and prefrontal cortex (PFC) (AD Progression Atlas, Wachter et al., 2024^29^). Each cell shows Spearman’s correlation ρ between gene expression and the ordinal pathology measure: composite AD stage, Braak NFT stage (I–VI), and CERAD neuritic-plaque density, computed separately within APOE-genotype groups. Cell colour indicates the genotype in which the correlation was significant: APOE3/3 (blue, “E3”), APOE-ε4 carriers (red, “ε4”), both genotypes (purple), or neither (grey, ns). The canonical RELN adaptor DAB1 declines with pathology in EC and ITG (significant in APOE3/3) and in PFC (significant in ε4 carriers); the APOE-axis genes LRP1 and DAB2 decline in V1/PFC and EC/PFC respectively in ε4 carriers, whereas APOE and ITGB1 rise modestly in selected regions/genotypes. Significance: *p < 0.05, **p < 0.01, ***p < 0.001. **b) Advancing AD pathology is accompanied, across brain regions, by an enrichment in disease-associated, antigen-presenting, interferon responsive, ribosomal and dystrophic microglial states with depletion of cytokine-responsive (CRM) states across all brain regions.** Heatmap of multilevel fgsea testing enrichment of ten curated microglial-state gene sets in advanced-versus-early AD (PathAdv-vs-Early) pseudobulk microglia from four brain regions (EC, ITG, PFC, V1; AD Progression Atlas, Wachter et al., 2024^29^). Microglial states: disease-associated (DAM), antigen-presenting (HLA), cytokine-responsive (CRM), interferon-responsive (IRM), homeostatic (HM) and ribosomal (RM) (Mancuso et al., 2024^26^), dystrophic (Streit et al., 2004^141^; 2009^141,142^); proliferating (Olah et al., 2020^143,144^); senescence core (Saul et al., 2022^145^; Hernández-Segura et al., 2017^146^; Dehkordi et al., 2021^143^); lipid-droplet-accumulating microglia (LDAM up; Marschallinger et al., 2020^47^) and APOE4/E4 lipid-droplet accumulating microglia (APOE4-LD; Victor et al., 2022^48^). Fill = normalised enrichment score (NES; positive = enriched in advanced AD; saturated at ±3). The DAM/LDAM, HLA, IRM states are enriched across all or most regions; the cytokine-responsive CRM state is depleted across all regions; and homeostatic, ribosomal and proliferating microglia show region-dependent changes — homeostatic microglia are enriched in EC, ribosomal microglia show the reverse (depleted in the EC, enriched elsewhere), and proliferating microglia are depleted only in the EC. The senescence-core signature is largely unchanged. Significance: BH-adjusted *padj < 0.05, **padj < 0.01, ***padj < 0.001. **c) DAB1 expression does not correlate with a microglial DAM/activation signature in the human brain (AD Progression Atlas).** Scatter plots show z-scored expression of DAB1 plotted against a microglial DAM/activation signature, defined as the mean within-region z-scored expression of 9 canonical disease-associated (DAM)/activated microglial markers (TREM2, TYROBP, GPNMB, AXL, LPL, CD9, CD68, CTSD, CAPG; core DAM genes from Keren-Shaul et al., (2017)^46^ and Mancuso et al., (2024)^26^ with CAPG from Monasor et al., (2020)^49^ & Pesämaa et al., (2023)^51^. Each dot is from a single donor and brain region (n=113 region-donor pseudobulk samples from 32 donors across entorhinal cortex (EC), inferior temporal gyrus (ITG), prefrontal cortex (PFC) and visual area 1 (V1), colour-coded by CERAD score (yellow to dark red = low to high scores). DAB1 was not correlated with DAM signature scores (Pearson’s r= –0.07, p= 0.47). Blue line shows regression fit. **d)** Heatmaps showing fgsea enrichment analysis of the RELN-core+DAB1 (115 genes) and RELN-core+DAB2 (113 genes) signatures in per-donor pseudobulk microglial transcriptomes from 4 regions of the AD Progression Atlas (EC, ITG, PFC, V1; Wachter et al., 2024^29^). **Left:** late Braak (V-VI) versus early Braak (0-II) tau pathology. Intermediate Braak stage III was not included. **Right:** Advanced versus Early pathology stage. Genes were ranked by the DeSeq2 Wald statistic from a per-region pseudobulk model. Fill= NES positive (red) indicates enrichment at the advanced stage. Benjamini-Hochberg *padj<0.05, **padj<0.01, ***padj<0.001.(n=113 donors in total).

**Supplementary Figure 2.** RELN-signalling pathways are associated with specific microglial states in early life and in disease-associated (DAM) and lipid-accumulating (LDAM) microglial states in Alzheimer’s Disease. **a**) Heatmap summarizing Fisher’s exact gene overlap test between curated RELN protein signalling axes (RELNcore-DAB1 (n=115 genes) and RELNcore-DAB2 (n=113 genes) and murine embryonic/postnatal microglial states (Hammond et al., 2019^43^; Li et al., 2019^42^). Significant % gene overlap is shown in cells with total number of genes per gene set and microglial state shown in brackets. Fisher’s BH padj values: *p<0.05, **p<0.01, ***-<0.001. b) RELN-DAB1/DAB2 protein signalling axis are enriched in disease associated (DAM) and lipid-accumulating AD-microglial states. Heatmap shows results from gene enrichment studies of RELNcore-DAB1 (n=115 genes) and RELNcore-DAB2 (n=113 genes) signalling arms (curated from Stringv12;Reactome;MSigDB databases) with 20,000 full-ranked Wald z genes of microglial AD states from the Mancuso et al., 2024^26^. (Homeostatic, HM; Cytokine responsive microglia (CRM/CRM-1/CRM2), Disease-associated (DAM), Antigen-presenting (HLA), Interferon responsive (IRM) and Ribosomal microglia (RM). BH * padj <0.05. Color-code range NES ±3. **c)** Heatmap summarizing Fisher’s exact/gene overlap test of RELNcore+DAB1 (n=115) or +DAB2 (n=113) with lipid-droplet accumulating microglial (LDAM) states (Haney et al., 2024^24^ (human); (Marschallinger et al., 2020^47^ murine); TREM2-lipid (Nugent et al., 2020^44^, murine), TREM2-LAM (Jaitin et al., 2019^129^) and LXR/Cholesterol (Nugent et al., 2020^44^ composite gene list with curated LXR/TREM2 efflux targets). Brackets show total number of genes comprising microglial signatures and the numbers in heatmaps show the number of overlapping genes between developmental and AD microglial states. **d)** Heatmaps shows results from a Fisher’s exact gene overlap test comparing embryonic and postnatal microglia (Hammond et al., 2019^43^; Li et al., 2019^42^) with adult microglial states (Mancuso et al., 2024^26^). Total number of genes per gene set and microglial state are shown in brackets. Fisher’s BH padj values: *p<0.05, **p<0.01,***p<0.001.

**Supplementary Figure 3.** Canonical RELN-signalling components are largely unchanged across APOE genotypes in human iPSC-microglia, except for APOE. **a**) RT-qPCR of RELN-axis genes across APOE3/3 and APOE4/4 iPSC-microglia. Bar plots (mean ± SEM, individual differentiations overlaid) show the RELN receptors VLDLR, LRP8, integrin receptor ITGB1, the adaptor DAB1 and P2RY12 microglial marker are unchanged across genotype, except for significant **APOE upregulation in APOE4/4;** *n* =3-4 differentiations. **b)** Immunocytochemistry of the canonical RELN intracellular adaptor **DAB1** and its tyrosine-phosphorylated forms **pY232** and **pY198**, co-labelled with the microglial marker **IBA1**, in APOE iPSC-microglia. **c)** Bar plots show the top 12 features with the largest loadings on the genotype axis, coloured by feature class (texture (purple) or intensity (orange)) of DAB1pY232 (top) or DAB2 (bottom) immuno-stained iPSC-microglia. Top bar plot linked to Fig. 2c: PC1 axis is driven by Ring/perinuclear DAB1pY232 Compact 30% SERSpot texture features which separates along the genotype axis. Bottom plot linked to Fig 2d showing CAPG SER texture features and cell-width to length ratio rather than DAB2 intensity, indicating genotype separation reflects morphological/texture differences and not DAB2 abundance.

**Supplementary Figure 4.** Expression analysis of RELN-DAB1 signalling in APOE iPSC-microglia (BIONIO37A/A-4) (**a**) and in inducible transcription-factor differentiated ioMG (BitBito) (b). Graphs show gene expression in log2 foldchange transcripts per million (log2FC TPM) of RELN receptors (LRP8, VLDLR), the adaptor proteins DAB1, DAB2, RELN, APOE and integrin receptors (ITGB1, ITGB8), together with microglial markers (C1QA, P2RY12), and hub genes involved in microglial response to amyloid and tau (MITF, TYROBP) (n=4 independent culture experiments for each genotype) in a) APOE iPSC-microglia (BIONIO37A/A-4) and in b) transcription-factor inducible iPSC-Microglia in a single APOE3/E3 and two APOE4/E4 clones (Clone 69 and Clone83)(BitBio) (n=3 for each genotype).

**Supplementary Figure 5.** Validation of DAB2 siRNA knockdown in APOE iPSC-microglia. **a)** Immuno-localisation of DAB2 (green) with Hoechst-stained nuclei (blue) in APOE3/3 iPSC-microglia (BIONIO37A) after Ctrl-siRNA, DAB2 siRNA-96 (targets both DAB2-201 and DAB2-202 isoforms) or siRNA-98 (targets the ubiquitous DAB2-201 isoform only). **b)** DAB2 transcript level (log2 TPM+1) across Ctrl-siRNA, DAB1 knockdown (siRNA-893, –894) and DAB2 knockdown (siRNA-96, –98), in APOE3/3 (squares) and APOE4/4 (triangles) iPSC-microglia (*n* = 3 differentiations per genotype). DAB2 is reduced by the DAB2 siRNAs — most strongly by siRNA-96 — but not by the DAB1 siRNAs. One-way ANOVA with post-hoc test; **p < 0.01, ***p < 0.001. **c)** Transcriptional perturbation by DAB2 knockdown. Number of up– (red) and down– (blue) regulated log2 foldchange DEGs (DESeq2, padj < 0.05; model ∼Batch + siRNA_class) per siRNA × genotype (E3/96, E4/96, E3/98, E4/98), pooled across genotypes (Pooled 96, Pooled 98, and convergent (significant with both siRNAs). siRNA-98 produces more DEGs than siRNA-96 (Pooled 98: 2,534 up / 1,754 down vs Pooled 96: 335 / 190; convergent: 222 up). **d)** qRT-PCR of DAB2 transcripts using primers detecting both DAB2-201 and DAB2-202 isoforms (top) or the DAB2-201 isoform only (bottom), in APOE iPSC-microglia after siRNA-96 (both isoforms) or siRNA-98 (DAB2-201 only) and normalised to housekeeping gene ARPC5. Each dot = an independent differentiation (APOE3/3 *n* = 3, APOE4/4 *n* = 3). One-way ANOVA with multiple-comparison post-hoc test; *p < 0.05, **p < 0.01, ***p < 0.001. **e) Volcano plots showing significant up-regulated (red) and downregulated genes (blue) in APOE3/E3 and APOE4/E4 iPSC-microglia following DAB2 knockdown of both DAB2-201 and DAB2-202 isoforms (siRNA-96) or only targeting DAB2-201 isoforms (siRNA-98). f) DAB2 knockdown concordance across APOE genotypes.** Scatter plot shows the per gene log2 foldchange of DAB2 knockdown versus control siRNA in APOE3/E3 (x-axis) compared with APOE4/E4 (y-axis) iPSC-microglia, for siRNA-96 (targets both DAB2-201 and DAB2-202 isoforms) (left) and siRNA-98 (targets DAB2-201 isoform alone) (right). Blue dots are significantly and concordantly regulated genes across genotypes (DESeq2 padj<0.05), medium-grey dots are genes significant on one genotype only; red circled dots show the most genotype-divergent genes with pale grey dots showing all other detected non-significant genes. The blue line is the regression fit through genes significant in both. Knockdown effects were fully concordant in direction (100% significance agreement) and strongly correlated in magnitude between APOE genotypes (siRNA-98: Pearson r =0.86, n=324; siRNA-96: r=0.84, n=23). DAB2 itself (left) was reduced to a greater degree in APOE3/E3 than APOE4/E4 microglia when both isoforms were targeted. **g) DAB2 isoform-divergent pathway enrichment (siRNA-96 vs siRNA-98).** Differential pathway enrichment between the two DAB2 knockdowns in iPSC-microglia. siRNA-96 targets both DAB2 transcript isoforms (DAB2-201 and the amyloid-responsive DAB2-202) and achieves a stronger overall knockdown, whereas siRNA-98 targets DAB2-201 only, leaving DAB2-202 intact. Bars show the difference in normalised enrichment score (ΔNES = NES_siRNA-96 − NES_siRNA-98), pooled across APOE3/3 and APOE4/4 and adjusted for genotype, from fgsea (multilevel, infinite permutations; min size 5– 10, max 500) on genes ranked by the DESeq2 Wald statistic for the direct siRNA-96 vs siRNA-98 contrast. Only pathways significant at FDR < 0.05 are shown. Orange bars (ΔNES > 0) are pathways more strongly enriched with siRNA-96 (loss of both isoforms / stronger knockdown); blue bars (ΔNES < 0) are pathways more strongly enriched with siRNA-98 (DAB2-201 knockdown, DAB2-202 retained). Asterisks denote Benjamini–Hochberg FDR: *FDR < 0.05, **FDR < 0.01, ***FDR < 0.001. n = 14 pooled samples (siRNA-96 n = 6, siRNA– 98 n = 8; APOE3/3 and APOE4/4 combined).

**Supplementary Figure 6.** DAB1 knockdown produces concordant pathway-level changes — in phagocytic, oxidative phosphorylation, lysosomal and amyloid-β-processing programmes — and enriches pro-inflammatory cytokine signalling and CRM microglial states, mirroring DAB2 knockdown. **a**) DAB1 knockdown validation. DAB1 immunostaining (green) with nuclear Hoechst (blue) in APOE3/3 iPSC-microglia after 48h of Ctrl-siRNA, siRNA-893 or siRNA-894. b) left panel: boxplot of DAB1 transcript level (log2 TPM+1) across all siRNA conditions (Ctrl, 893, 894, 96, 98), with APOE genotype marked (E3 square, E4 triangle); DAB1 falls with the DAB1 siRNAs at trend level (Ctrl vs 893 p = 0.054; vs 894 p = 0.056; Wilcoxon test) but not with the DAB2 siRNAs (p = 0.282). Right **panel**: number of DESeq2 DEGs (Wald test, padj < 0.05, BH; model ∼Batch + siRNA_class) per contrast — APOE4/E4 siRNA-893: 2 up; APOE genotype convergent effects of siRNA-893: 14 up / 35 down; APOE genotypes convergent effects of siRNA-894: 3 up / 2 down (only contrasts with ≥1 DEG shown). DAB1 knockdown yields far fewer DEGs than DAB2 knockdown, consistent with DAB1’s low baseline expression and the modest knockdown efficiency. **Bottom panels:** Volcano plots show few significantly differentially expressed genes with DAB1 siRNA knockdown (siRNA-893 & siRNA-894) across pooled APOE genotypes. **c) Functional-pathway enrichment of pooled DAB1 siRNA KD effects in APOE iPSC-microglia.** Heatmap of fgsea NES (MSigDB Hallmark + Reactome + GO:BP, ∼9,600 gene sets) for DAB1 knockdown (Pooled siRNA-893 & siRNA-894 vs Ctrl within APOE3/E3 and APOE4/E4 genotypes, or for DAB1 KD APOE4/E4 vs APOE3/E3 CtrlsiRNA contrasts; significant pathways shown, coloured by NES. fgsea-multilevel (minSize = 15, maxSize = 500, permutation size infinite]); stars BH FDR *padj < 0.05, **< 0.01, ***< 0.001. TNFα-NFκβ signalling (APOE4/E4 +1.60, APOE3/E3 +1.90) and inflammatory response (APOE3/E3 +1.88) and cytokine-mediated signalling (APOE3/E3 +1.39) were significantly upregulated. In contrast, phagocytosis (APOE4/E4 –2.08), phagocytosis engulfment (APOE4/E4 –2.21), oxidative phosphorylation (APOE4/E4 –1.68, APOE3/E3 –2.57), mitochondrial proteostasis (APOE4/E4 –1.84, APOE3/E3 –2.15), complement cascade (APOE4/E4 –1.96), fatty-acid metabolism (APOE4/E4 –1.88, APOE3/E3 –1.99), and amyloid-β-clearance pathways (APOE4/E4 –2.11) were among the most significantly depleted. The genotype x knockdown interaction was negative for TNFα-NFκβ signalling (–2.10) and senescence associated secretory phenotype SASP (–1.86), indicating these inflammatory programmes were induced more strongly in APOE3/E3 iPSC-microglia. These changes mirror the DAB2-knockdown pathway signature (Fig. 4). **d) Transcriptional concordance of DAB1 and DAB2 silencing.** Per-gene scatter of sign(log2FC) × −log10(p) for pooled DAB1-KD-vs-Ctrl (x-axis) versus pooled DAB2-KD-vs-Ctrl (y-axis) across APOE genotypes (n = 20,526 genes); points coloured both-up (red), both-down (blue), discordant (grey), ns (light grey). Spearman ρ = 0.762 (pooled); concordance is greatest in APOE3/3 (ρ = 0.836; E4 ρ = 0.688). **e) Depletion of lysosomal and phagocytic genes following DAB1/2 KD.** Heatmap of DESeq2 log2FC (KD vs Ctrl) for endolysosomal/phagocytic genes (FUCA2, GLB1, NPC2, MAN2B1, MAN2B2, CTSA, PSAP), the γ-secretase/amyloid component NCSTN, and CD68 across DAB1-893, DAB1-894, DAB2-96 and DAB2-98 contrasts. DAB2 knockdown drives broad significant depletion; DAB1 knockdown phenocopies the direction but reaches significance only for MAN2B2 (siRNA-893; siRNA-894 shows essentially no change). Stars: *padj < 0.05, **< 0.01, ***< 0.001. **f) Functional-pathway NES preferential enrichment in DAB1 vs DAB2 pathways**. Scatter of fgsea NES for DAB1-KD-vs-Ctrl (y) versus DAB2-KD-vs-Ctrl (x), pooled across genotypes. Points: DAB1-enriched (orange, n = 29), DAB2-enriched (blue, n = 94), shared (grey, n = 53), ns. DAB1 knockdown shows enrichment of interferon-response pathways and depletion of lipoprotein-particle and mitochondrial-fatty-acid pathways, with DAB2 KD showing a depletion in antigen-presentation pathways. **g) DAB1 vs DAB2 microglial-state and RELN-axis preferential enrichment.** Scatter of fgsea NES (DAB1-KD-vs-Ctrl, y; DAB2-KD-vs-Ctrl, x) for curated microglial-state and RELN-signalling gene sets. Points: DAB1-enriched (orange, n = 3), DAB2-enriched (blue, n = 5), shared (black, n = 13), ns (n = 2). DAB1 knockdown preferentially shifts microglia toward interferon-responsive (IRM) with DAB2 knockdown shifting microglia towards transitioning cytokine-responsive states; DAB1 knockdown tracks IRM via the RELN-VLDLR-LRP8-DAB1 axis, while DAB2 knockdown negatively correlates with a broader range of lipid– and DAM-activated states (e.g. LXR/cholesterol-efflux, TREM2-lipid).

**Supplementary Figure 7.** Analysis of microglia in chronic Reln-deficient mouse model. **a**) Graphs show Reelin, ApoER2, Vldlr1 and ApoE transcript expression levels detected by RT-qPCR analysis of FACS Cd11b^+^ microglia and macrophages purified from the E18.5 rat forebrain meninges (Men), dorsal cerebral cortex (Cx) or subpallium (Sp) and normalised for housekeeping gene GAPDH. (n=2 experiments, each containing 10-20 pooled pups). **b-e) Validation of *Reln-deficient* mice.** Actin-Cre mediated depletion of Reelin in P0.5 transgenic ActinCre+/−;Relnfl/fl mice results in the global degradation of Reln protein isoforms and recapitulates reeler forebrain lamination defects. **b)** In-situ hybridisation for Reln mRNA (left panels, magenta) in coronal sections through the caudal dorsal cortex and hippocampus of P0.5 ActinCre+/−;Reln+/+ (top) and ActinCre+/−;Relnfl/fl (bottom) mice showing ectopic distribution of Reln-expressing cells. Immunohistochemical staining for Reln proteins using the mG10 antibody (N-terminal, green) and C17 antibody (C-terminal, magenta) shows efficient degradation of Reln protein isoforms in ActinCre+/−;Relnfl/fl mice. **c)** Western blot of liver and cerebellar lysates from P22 ActinCre+/− Reln+/+ (RELN-WT) control and ActinCre+/− Relnfl/fl mutant (RELN-KO) mice probed with the mG10 antibody. **d)** Coronal sections through the P0.5 caudal dorsal cortex and hippocampus stained for upper-layer Satb2+ neurons (red), future layer IV Ctip2+ neurons (magenta), intermediate progenitors (Tbr2+), and newly differentiated post-mitotic Tbr1+ neurons; **e)** Transverse view of P0.5 Reln-WT and Reln-KO cerebellum immunostained for Calbindin showing dysmorphic folia and aberrant Purkinje neuron positioning. **f)** Coronal sections through the rostral-medial dorsal cortex of E14.5 ActinCre+/−;Reln+/+ (Reln-WT, left) and ActinCre+/−;Relnfl/fl (Reln-KO, right) mouse forebrains immunostained for IBA1 and cleaved-caspase 3 (CC3, green). Graphs depict total numbers of IBA1+ cells +/− SEM quantified in 300 um width bins within the dorsal-lateral cortex. **g)** Barplots mean values ± SEM of morphological parameters scored in cortical microglia, with dots denoting number of mice analysed (n= total of 806-831 cells from n=4-5 mice Reln-WT, n= 559-801 cells from n=4 mice Reln-KO). **h)** Hippocampal microglia show no changes in morphology or expression of CD68 in P22 Reln-KO mice. Panels show x40 high magnification of the CA1 hippocampus of P22 ActinCre+/−;Reln+/+ (Reln-WT, top) and ActinCre+/−;Reln−/− (Reln-KO, bottom) mice, immunostained for the ionised calcium binding protein IBA1 (red) and the endolysosomal marker CD68 (magenta). **i)** Graphs show mean values +/− SEM of scored morphological parameters of hippocampal microglia with dots denoting individual P22 mice (morphology: 217-326 microglia from n=3 Reln-WT mice, 119-251 microglia from n=3 Reln-KO mice for each parameter).

**Supplementary Figure 8.** Principal-component analysis of the microglial-state / RELN-axis NES landscape — PC3 resolves homeostatic/cytokine-responsive from antigen-presenting/disease-associated programmes. PCA of the fgsea-NES matrix (same 10 features as Fig.6: RELN→DAB1, RELN→DAB2, MG_HM, MG_CRM, MG_IRM, MG_DAM, MG_HLA, Haney-LDAM, Victor APOE4-lipid, LXR-cholesterol efflux across *n* = 28 contrasts) drawn from six contrast groups: AD Progression Atlas (red circle, n=6), Haney e al., 2024^24^ AD snRNA of microglia in the prefrontal cortex (dark-red triangle, n=3) or iPSC-MG APOE (orange square, n=6), experimental APOE-iPSC microglia in this study (BIONIO37A/A-4) genotype (blue diamond, n=2) or following DAB1/DAB2 KD (blue circle, n=6) and developmental, adult and ageing microglia (green triangle, n=5 Kracht et al., 2020^27^; Galatro et al., 2017^28^; Ostrem et al., 2024^124^). Companion to the Fig 6 PC1-vs-PC2 biplot, here showing the third component. **a)** PC2 (19.7 % variance) vs PC3 (14.0 %) biplot. Points = contrasts; arrows = feature loadings (scaled). PC2 separates a CRM·HM pole (left) from a DAM·LDAM pole (right); PC3 separates an HM·CRM pole (top) from an HLA·DAM pole (bottom). The iPSC-APOE (BIONIO37A/A-4) DAB1-KD and DAB2-KD samples (blue, dashed ellipses) occupy the negative-PC1 “DAB-KD pole”, opposite the AD-progression cluster (red ellipse); the dashed arrow marks the hypothesised DAB-KD ↔ APOE4/AD-progression axis. **b)** PC3 loadings (bar chart). Positive (red): MG_HM (+0.58), MG_CRM (+0.57), Victor et al., 2022^48^ APOE4-lipid (+0.26), Marschallinger-LDAM^47^ (+0.20); near-zero: RELN→DAB2 (–0.03), RELN→DAB1 (−0.10); negative (blue): MG_IRM (−0.14), MG_DAM (−0.21), MG_HLA (−0.33). PC3 thus contrasts homeostatic/cytokine-responsive/lipid signatures against antigen-presenting (HLA) and disease-associated/interferon (DAM/IRM) signatures.

**Supplementary Figure 9.** WGCNA of n = 39 APOE iPSC-microglia (BIONIO37A / BIONIO37A-4) resolves fourteen co-expression modules (seven trait-significant); APOE-axis genes map to discrete modules. **a**) PCA of the n = 39 batch-corrected samples: PC1 (27.3 % of variance) separates samples by APOE genotype (E3 blue, E4 red), while siRNA class (Ctrl ●, DAB1 ▴, DAB2 ▪) does not drive a major axis. PC2 = 12.6 %. b) Gene dendrogram and module assignment (minModuleSize = 30 genes, mergeCutHeight = 0.25, deepSplit = 2). WGCNA was performed on the variance-stabilised (VST; DESeq2 v1.46), limma-batch-corrected expression matrix of n = 39 APOE3/3 and APOE4/4 iPSC-microglia. Low-expression genes were removed (retaining genes with ≥10 counts in ≥3 samples; 23,320 of 63,187 genes), and the top 10,000 most-variable genes were carried forward. A soft-thresholding power β = 12 (scale-free R² = 0.85) was applied to build a signed-hybrid weighted adjacency matrix; TOM was computed and genes clustered by dynamic tree cut, merging modules whose eigengenes correlated r > 0.75. Fourteen modules were obtained — turquoise (1,514 genes), blue (1,001 genes), brown (927 genes), yellow (615 genes), green (589 genes), red (325 genes), black (279 genes), pink (242 genes), magenta (232 genes), purple (150 genes), greenyellow (130 genes), tan (126 genes), salmon (101 genes), cyan (85 genes) — with 3,684 genes (36.8 %) unassigned (grey). Module membership (MM) is the Pearson correlation between each gene and its module eigengene (the first PC of the module’s expression across the 39 samples); intramodular connectivity (kWithin) is the summed adjacency to all other module genes. In module-trait analysis (Fig 7), DAB2 knockdown correlated with seven modules — brown and green (up) and purple, greenyellow, salmon, magenta and blue (down) — whereas the APOE-genotype and DAB1-knockdown trait columns showed no significant module correlations; the lower tracks show each module’s APOE-ε4 direction and DAB2-KD effect. c) Module membership (MM = signed kME) of RELN/APOE-axis genes and the top hub genes of each module. LRP8 (MM 0.84, hub) and LRP1 (0.76) are members of the blue module; DAB2 (0.71) of the purple module; ITGB8 (0.54) of the brown module; RELN (0.60) of the black module; and ITGB1 (0.74) and VLDLR (0.38) of the turquoise module; APOE (MM ≈ 0) and DAB1 (0.11) passed the variance filter but were unassigned (grey). d) Projection onto human AD microglia. Projection of the seven significant n = 39 BIONIO37A/A-4 iPSC-microglia WGCNA modules — brown (CRM), green (no microglial state), magenta (HM), blue (RM/HLA), greenyellow (DAM), salmon (HM) and purple (DAM/LDAM) — onto human microglial pseudobulk from the AD Progression Atlas (Wachter et al., 2024^29^) across four brain regions: n=113 total donors: entorhinal cortex (EC, n = 23 donors), inferior-temporal gyrus (ITG, n = 31), prefrontal cortex (PFC, n = 28) and primary visual cortex (V1, n = 31). Within each region, genes were ranked by the Spearman correlation of their expression with CERAD scores (amyloid) across donors, and each module’s gene set was tested against that ranking with a correlation-aware competitive gene-set test (CAMERA). Heatmap colour = rank-biserial effect (module genes vs background; + = up with CERAD, − = down), clipped to ±0.5. After accounting for inter-gene correlation, no module is significantly associated with CERAD pathology in any region (all BH-FDR > 0.05; Stars “*” marks cells significant only by the naive rank-sum). Braak (tau pathology) associations were tested in parallel and were no association with pathology in specific brain regions. **e) Overlap enrichment of the DAB2 siRNA knockdown WGCNA eigengene modules (purple DAB2/TREM2, blue LRP8/LRP1, brown ITGB8, salmon MRC1/P2RY12) against independently defined microglial-state signatures from mouse**^46,47,62,65^ **and human brain**^61,63,64^ **AD transcriptomic studies.** Enrichment was scored as the Fisher’s exact test odd’s ratio of gene-overlap (Benjamini-Hochberg FDR; *FDR<0.05, **<0.01, ***<0.001). The purple DAB2/TREM module was significantly enriched in disease-associated (DAM)/Microglial neurodegenerative (MgND) TREM2-dependent signatures, whereas the salmon module was enriched in homoestatic microglial signatures across mouse and human studies.

## Supplementary Material

Supplementary Table1_Reln-pathwaylists

Supplementary Table2_Pathway enrichment analysis of APOE4/E4 versus APOE3/E3 Vehicle control APOE iPSC-microglia

Supplementary Table3_Pathway enrichment analysis of DAB2 siRNA knockdown APOE iPSC-microglia

Supplementary Table4_Pathway enrichment analysis of DAB1 siRNA knockdown APOE iPSC-microglia

Supplementary Table5_PCA analysis of DAB2 siRNA knockdown APOE iPSC-microglia with human brain datasets

Supplementary Table6_WGCNAmodules of DAB2 siRNA knockdown APOE iPSC-microglia

Supplementary Table7_DAB2 WGCNA module enrichment with Bellenguez dementia-risk loci

## Data Availability

The microglial RNA sequencing data sets analysed in this study are available under the following Gene Expression Omnibus (GEO) accession numbers: GSE254205 from Haney et al., 2024^24^,PMID: 38480892; GSE99074 from Galatro et al., 2017^28^ PMID: 28671693; and GSE141862 from Kracht et al., 2020^27^ DOI: 10.1126/science.aba5906. Microglial RNA sequencing processed data from the AD Progression Atlas was downloaded from https://ad-progression-atlas.partners.org Wachter et al., 2024^29^ DOI:10.1007/s00401-024-02704-2. The GWAS dementia-risk loci used in this study were obtained from Supplementary Table 5 & Supplementary Table 20 from Bellenguez et al., 2022 DOI:10.1038/s41588-022-01024-z. The microglial signatures used in this study were obtained from: Mancuso et al. (2024)^26^ DOI:10.1038/s41593-024-01600-y, Mathys et al. (2019)^106^ DOI:10.1038/s41586-019-11952-2, Keren-Shaul et al. (2017)^136^ DOI:10.1016/j.cell.2017.05.018, Sala-Frigerio et al. (2019)^45^ DOI:10.1016/jcelrep.2019.03.099 and Marschallinger et al., (2020)^47^ DOI:10.1038/s41593-019-0566-1. The bulk RNA sequencing data generated in this study from iPSC-APOE microglia with DAB1/DAB2 gene perturbation studies have been deposited under the Gene Expression Omnibus accession number: GEO#### RNASeq_BIONIO37A/A-4. All code and scripts used in this study have been deposited in Github.DOI 10.5281/zenodo.23214621.

