## Supplementary Fig. 1 for "The adaptor protein DAB2 cell-autonomously links APOE with disease-associated microglia (DAM)/TREM2 state dysregulation associated with Alzheimer’s disease"

**a**

| Gene | Composite Stage | | | | Braak | | | | CERAD | | | |
|---|---|---|---|---|---|---|---|---|---|---|---|---|
| | EC | ITG | V1 | PFC | EC | ITG | V1 | PFC | EC | ITG | V1 | PFC |
| RELN | E3 -0.59 * | +0.27 ns | -0.01 ns | -0.03 ns | E3 -0.62 * | +0.11 ns | -0.05 ns | -0.07 ns | -0.08 ns | +0.40 ns | +0.06 ns | +0.19 ns |
| DAB1 | E3 -0.72 ** | E3 -0.48 * | -0.31 ns | ε4 -0.87 * | E3 -0.73 ** | E3 -0.54 * | -0.34 ns | ε4 -0.87 * | E3 -0.64 * | -0.39 ns | -0.13 ns | -0.21 ns |
| LRP8 | +0.14 ns | -0.12 ns | -0.07 ns | -0.09 ns | +0.19 ns | -0.17 ns | -0.09 ns | -0.06 ns | +0.18 ns | -0.04 ns | -0.06 ns | -0.05 ns |
| ITGB1 | +0.38 ns | E3 +0.47 * | +0.25 ns | -0.03 ns | +0.38 ns | +0.33 ns | +0.20 ns | -0.03 ns | +0.16 ns | +0.38 ns | +0.20 ns | -0.18 ns |
| DAB2 | -0.14 ns | +0.26 ns | +0.26 ns | -0.17 ns | -0.13 ns | +0.12 ns | +0.23 ns | -0.21 ns | ε4 -0.88 * | +0.15 ns | +0.16 ns | ε4 -0.87 * |
| LRP1 | -0.38 ns | -0.13 ns | ε4 -0.87 * | ε4 -0.87 * | -0.34 ns | -0.31 ns | ε4 -0.87 * | ε4 -0.87 * | -0.21 ns | -0.07 ns | +0.41 ns | -0.46 ns |
| APOE | +0.24 ns | +0.51 ns | E3 +0.48 * | +0.48 ns | +0.26 ns | +0.49 ns | E3 +0.63 * | +0.50 ns | +0.21 ns | +0.50 ns | +0.51 ns | +0.40 ns |

E3/E3-driven   ε4-driven   Both   ns

Significance: *** p<0.001 ** p<0.01 * p<0.05

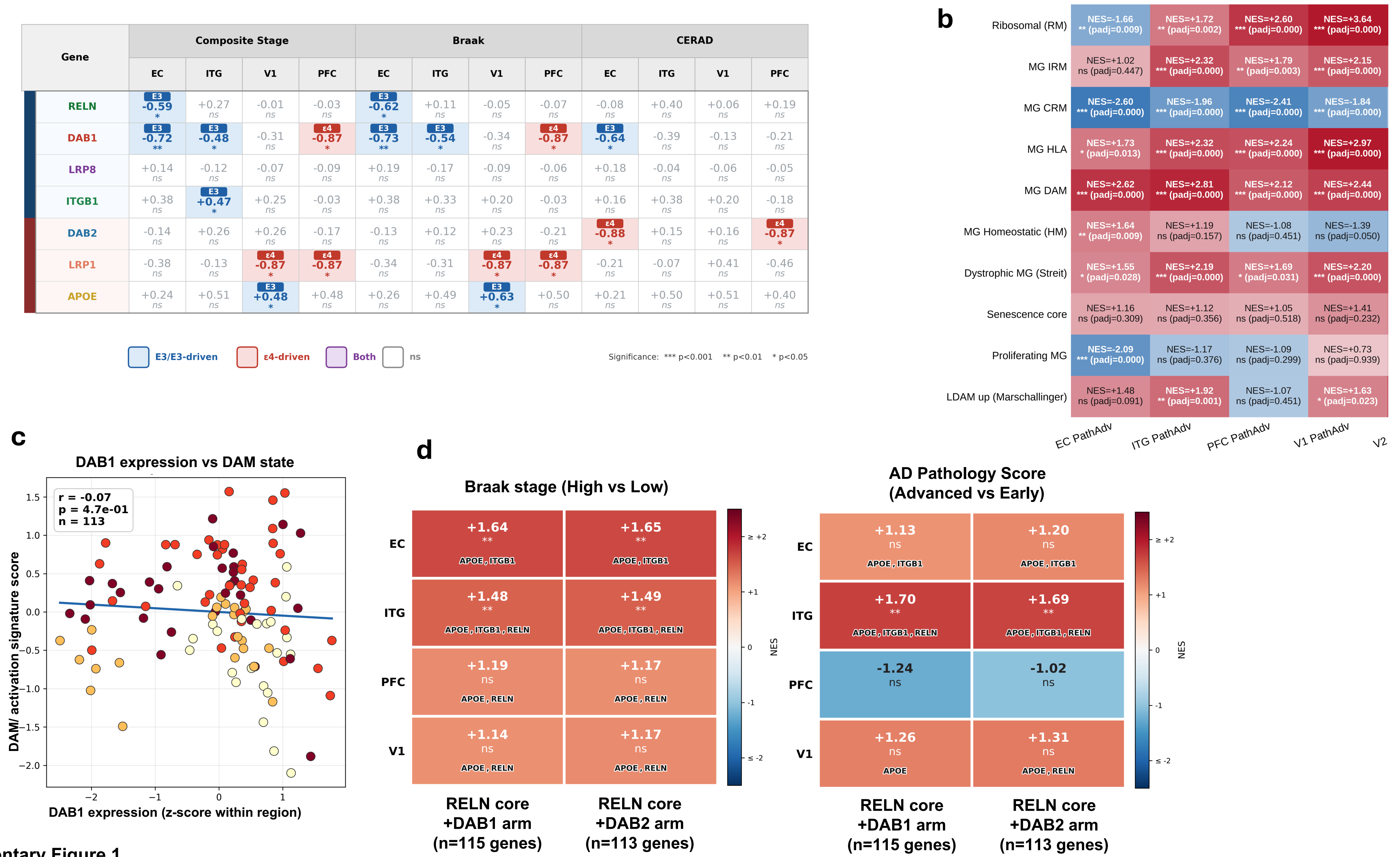


Supplementary Figure 1.
