## Supplementary figures and images for "The adaptor protein DAB2 cell-autonomously links APOE with disease-associated microglia (DAM)/TREM2 state dysregulation associated with Alzheimer’s disease"

### Supplementary Fig. 2

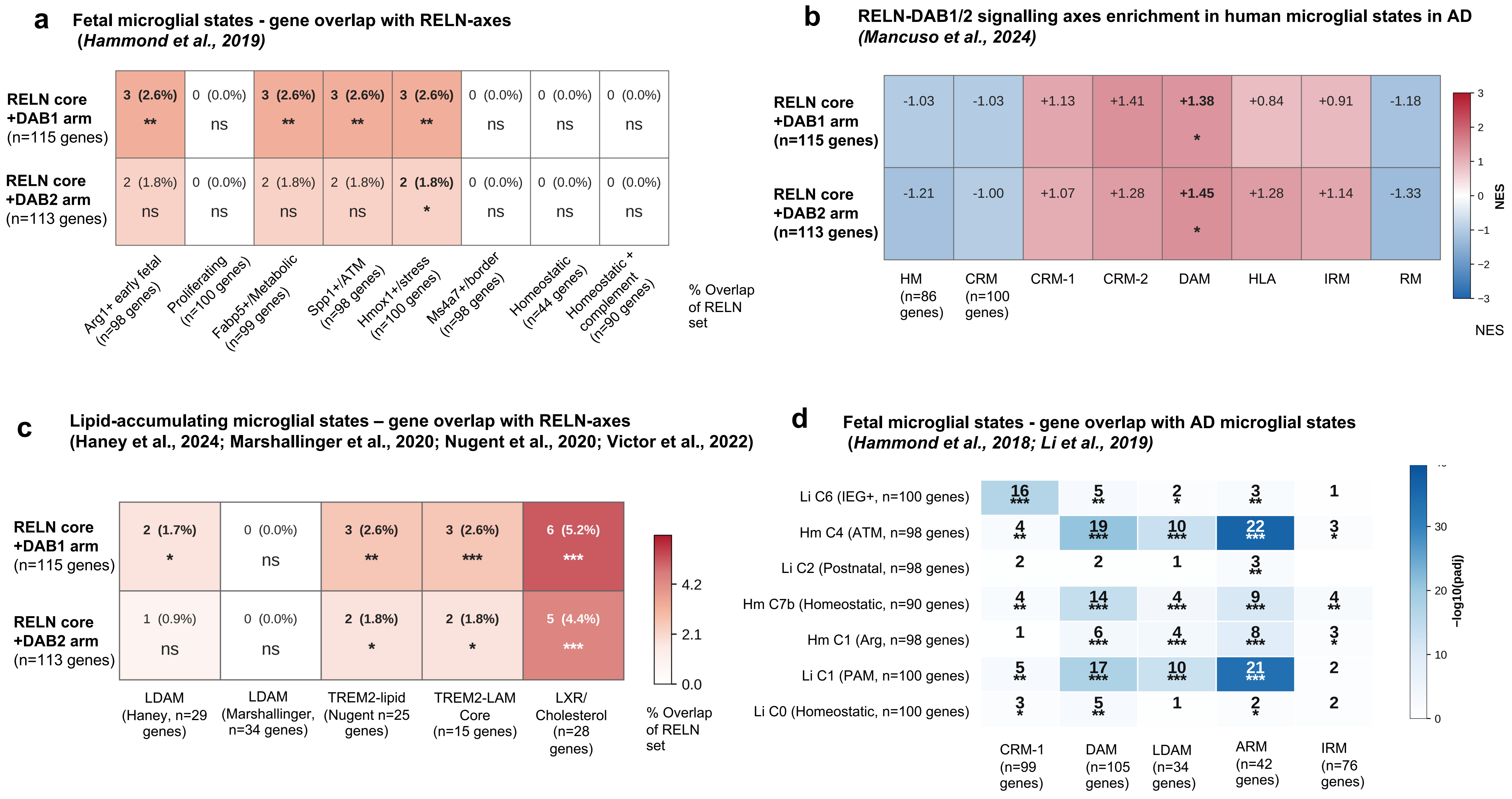


Supplementary Figure 2.

### Supplementary Fig. 3

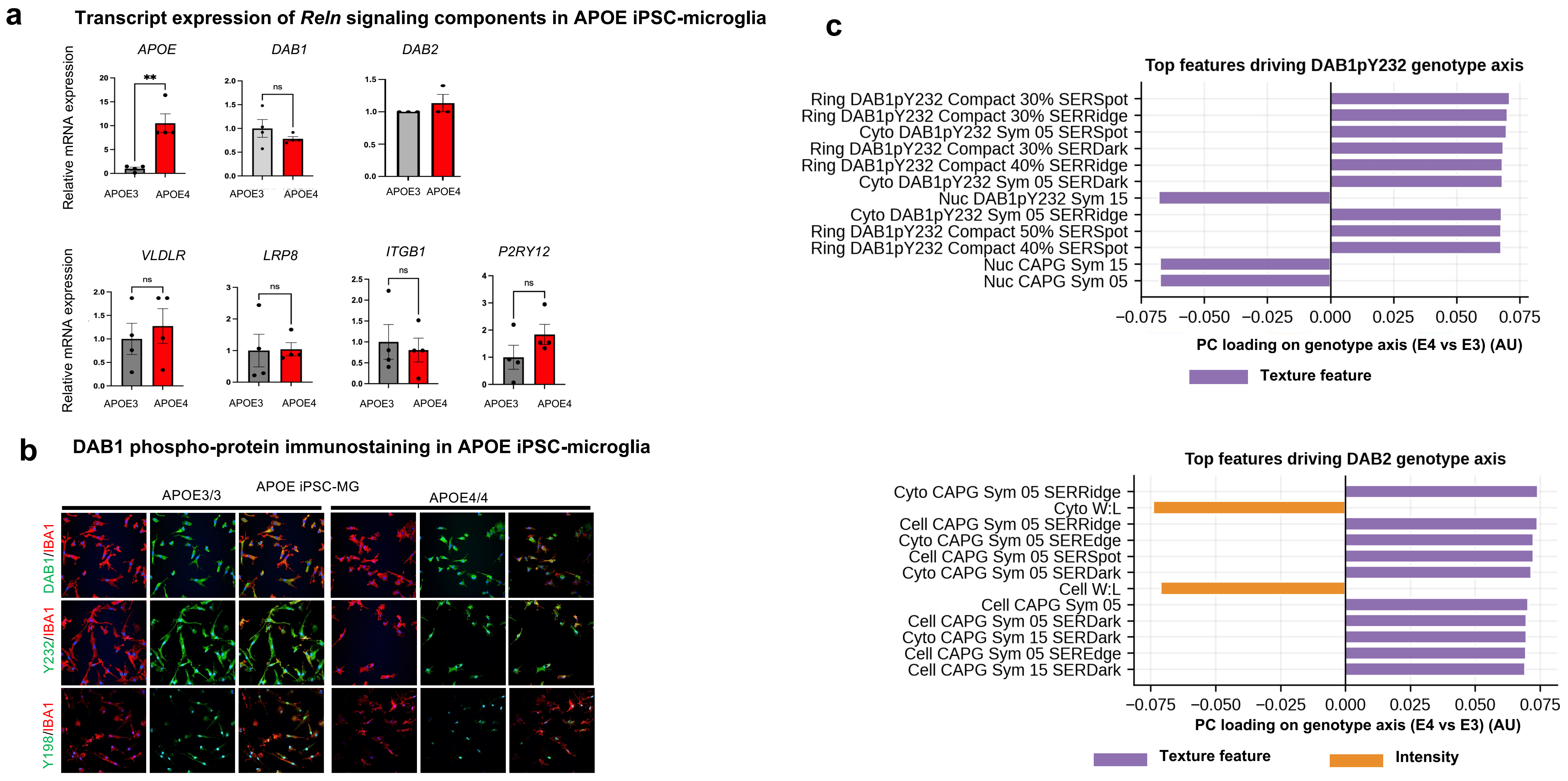


**Supplementary Figure 3.**

### Supplementary Fig. 4

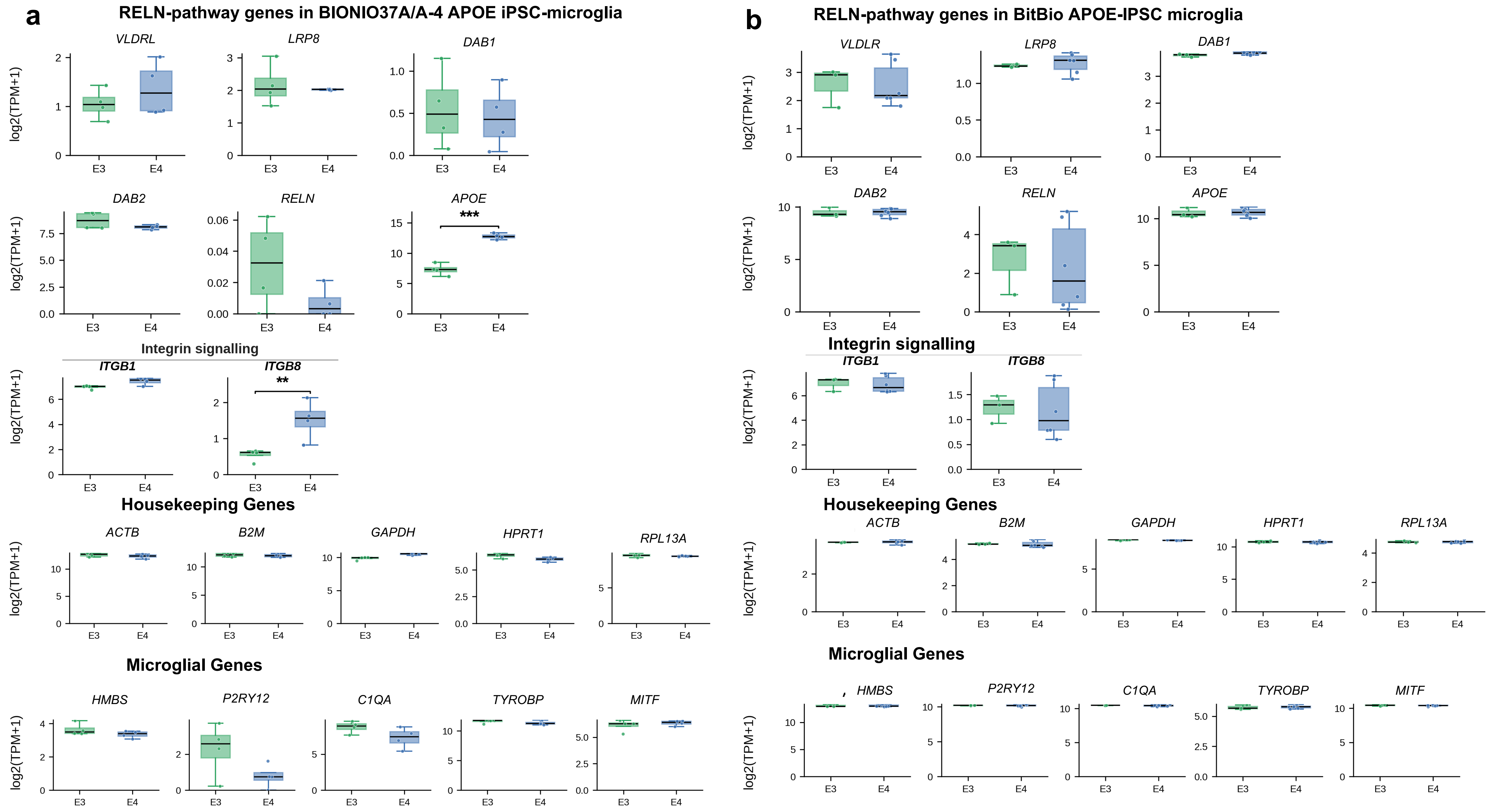


**Supplementary Figure 4.**

### Supplementary Fig. 5

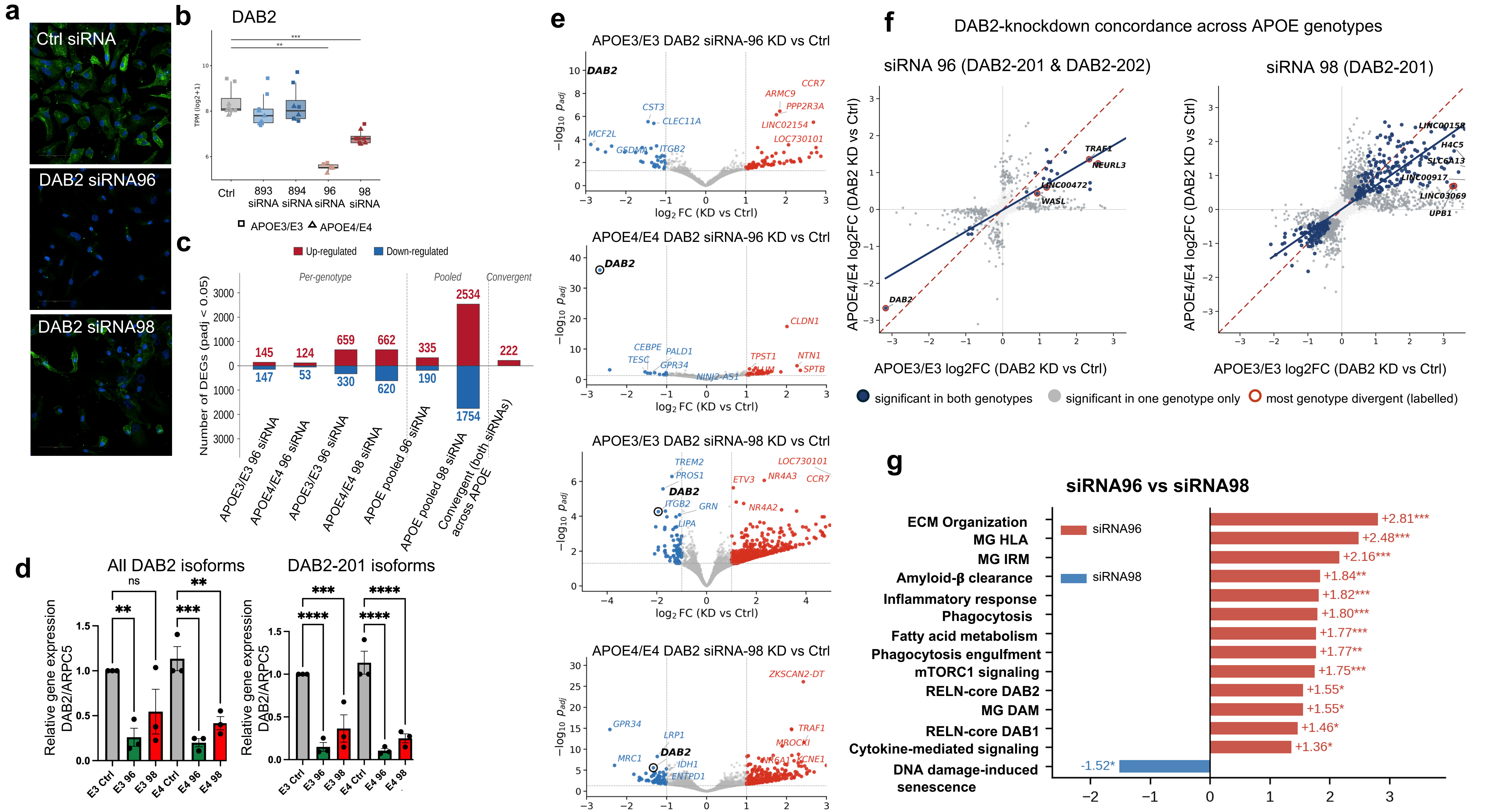


**Supplementary Figure 5.**

### Supplementary Fig. 6

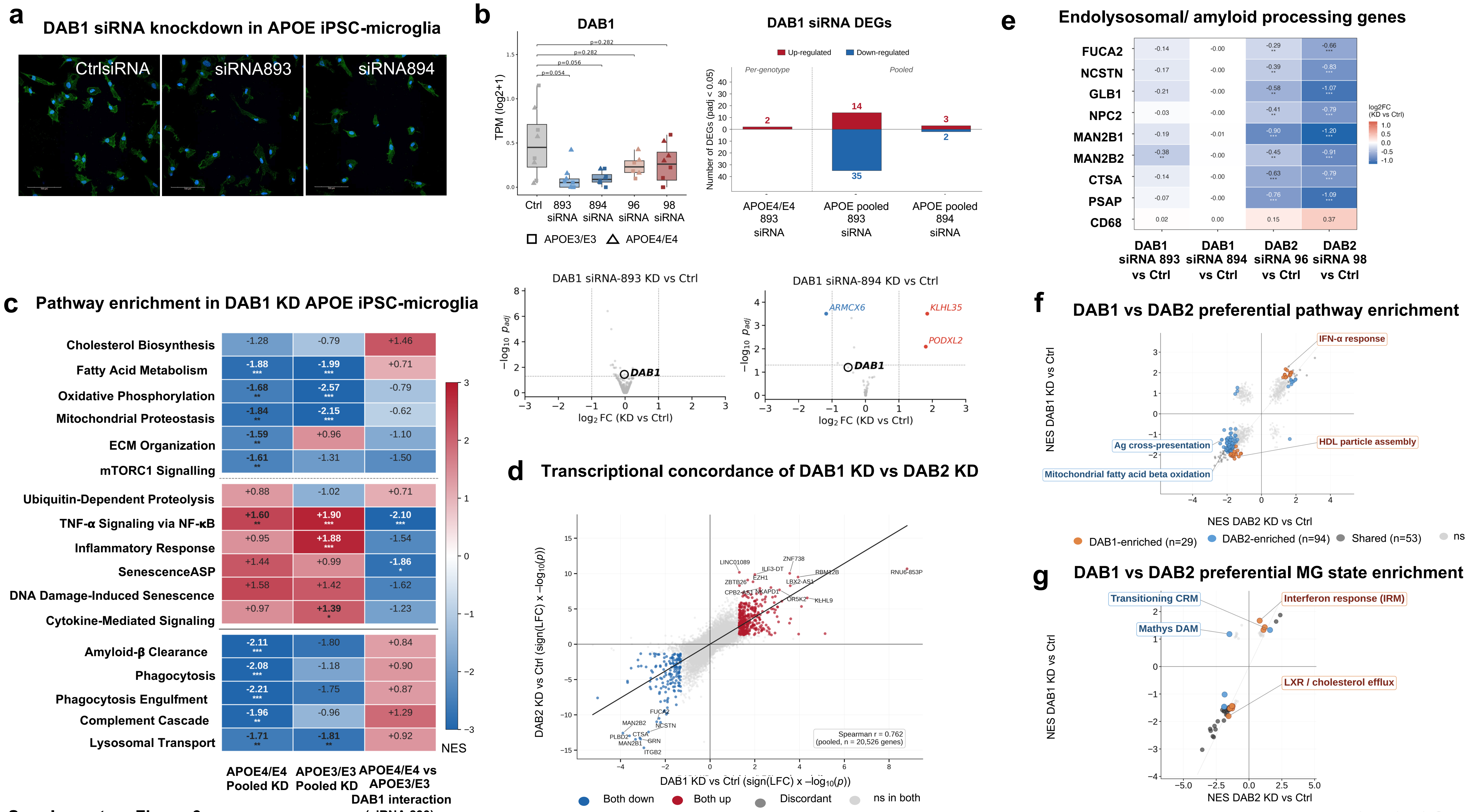


Supplementary Figure 6.

### Supplementary Fig. 7

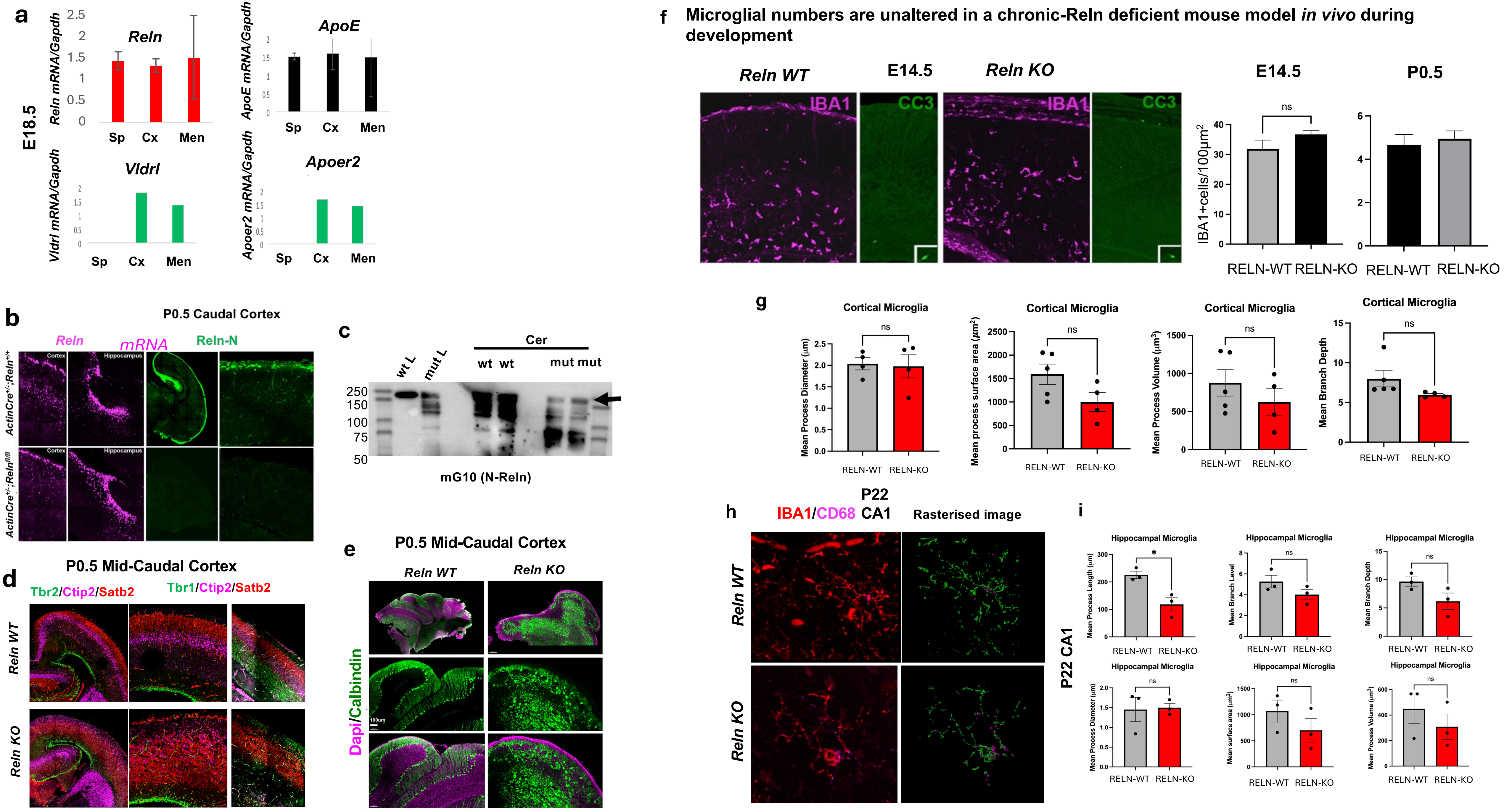


**Supplementary Figure 7.**

### Supplementary Fig. 8

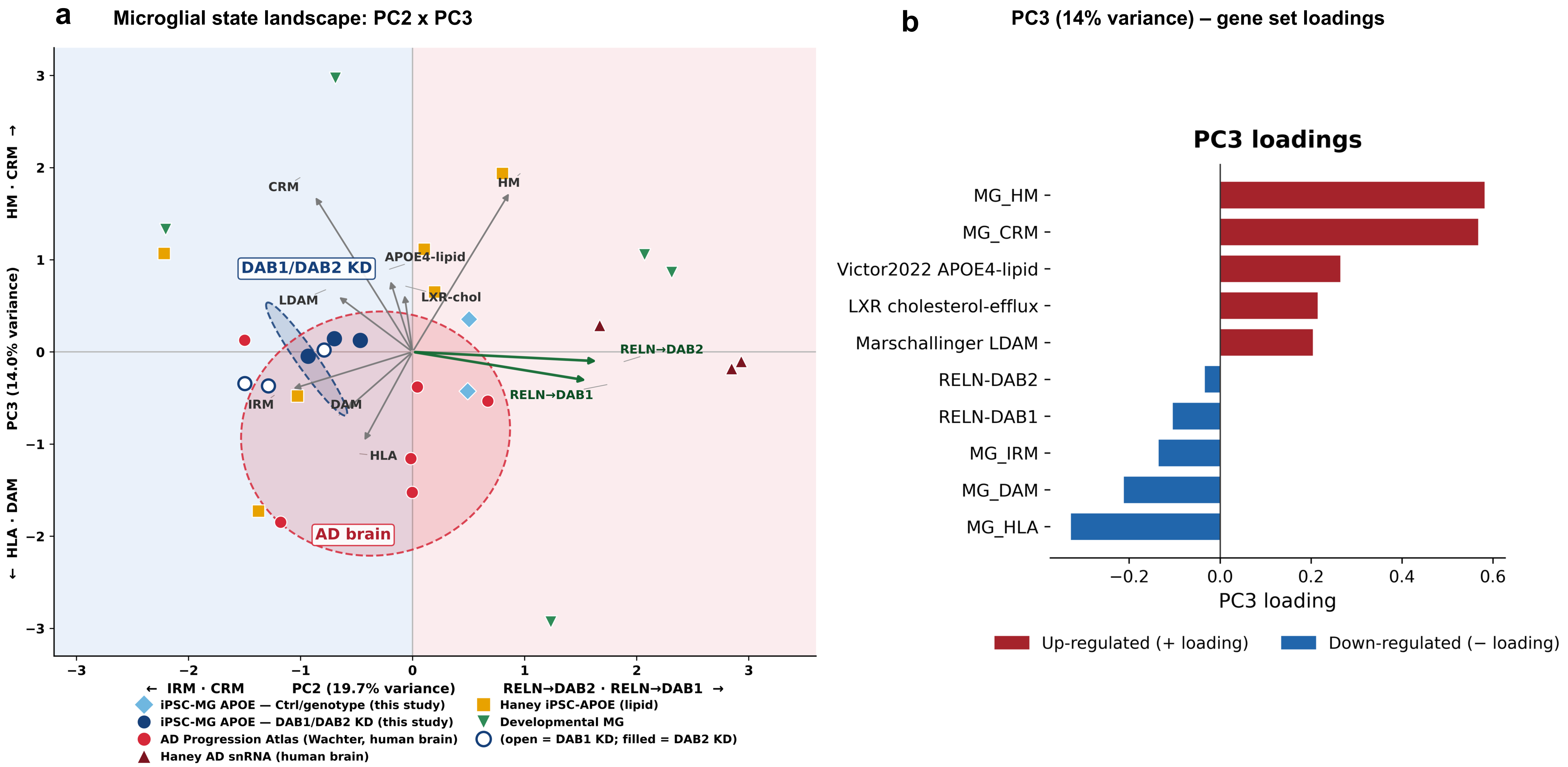


Supplementary Figure 8.

### Supplementary Fig. 9

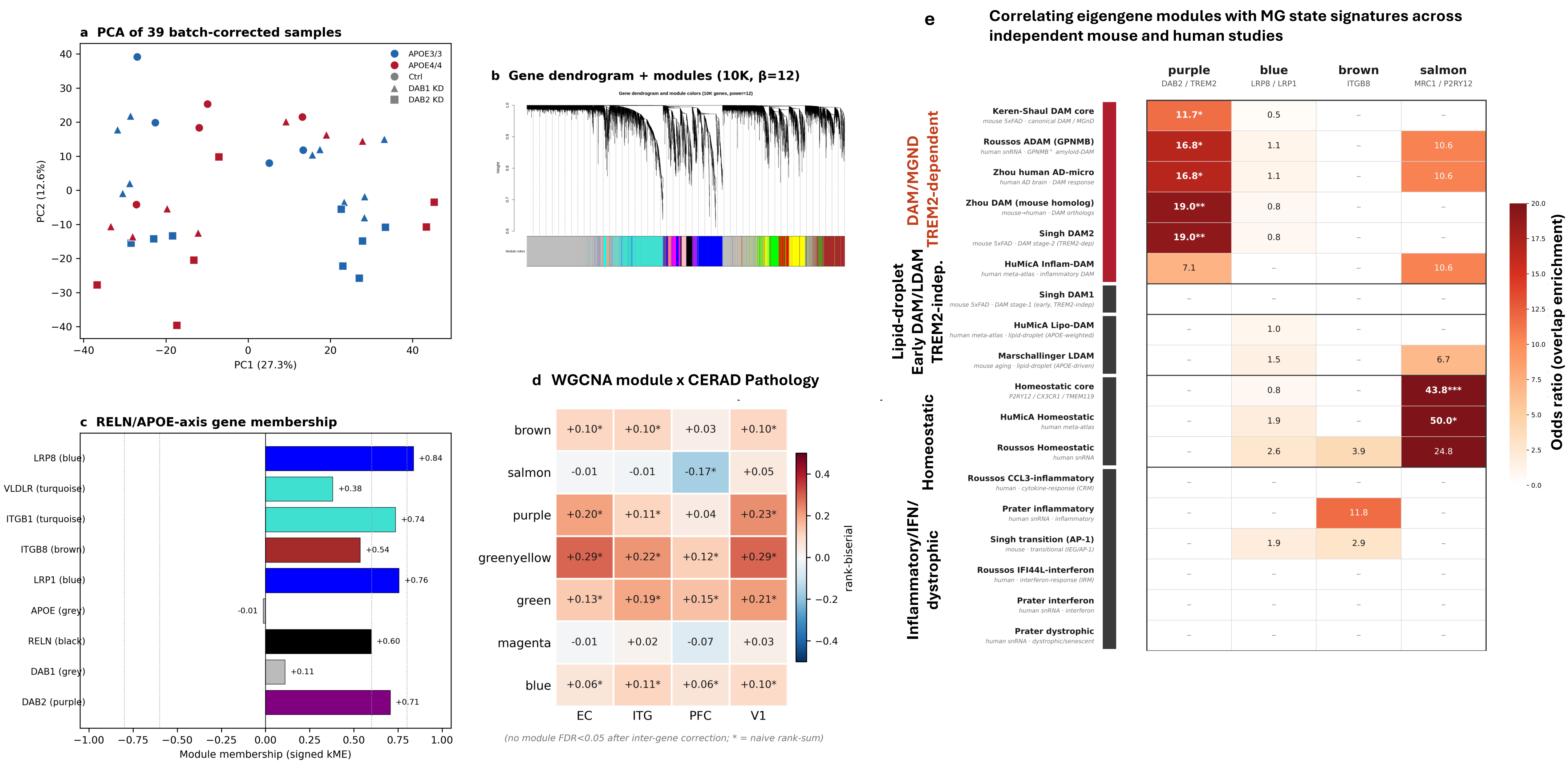


**Supplementary Figure 9.**
